# A ubiquitin chain-feeding mechanism for BRCA1-A

**DOI:** 10.64898/2026.06.05.730370

**Authors:** Andrea G Murachelli, Farid El Oualid, Titia K. Sixma

## Abstract

The BRCA1-A complex is a multi-subunit, metallo-deubiquitinating enzyme (metallo-DUB) involved in genome maintenance. BRCA1-A displays strict specificity for K63-linked ubiquitin, with a strong preference for long chains, but the mechanistic basis for this selectivity has remained unclear.

To address this, we developed a novel activity-based probe that is specific for metallo-DUBs and mimics di- or polyubiquitin chains of any linkage (di- and poly-ubiquitin_ATA_). We solved cryoEM structures of BRCA1-A bound to K63-linked probe chains of various length, capturing multiple conformational and catalytic states. The structures reveal how allosteric regulation of catalysis occurs within the complex and how BRCA1-A uses auxiliary ubiquitin-binding sites to engage substrate by avidity and to trigger processive cleavage. Crucially, avidity and processivity can only apply to long polyubiquitin chains, explaining BRCA1-A’s substrate preference. Together, these results establish BRCA1-A as a chain-shortening DUB specialised for trimming extended K63-linked polyubiquitin chains.

## Introduction

The BRCA1-A complex is a multi-subunit deubiquitinating enzyme (DUB) that regulates repair of DNA double-strand breaks (DSBs) arising from insults such as ionising radiation^1,2^, interstrand crosslinks^3,4^, or stalled replication forks^5^. BRCA1-A comprises ten subunits assembled into a two-armed, dimer-of-pentamers architecture, with each arm (pentamer) composed of ABRAXAS (also known as ABRAXAS1)^1,2^, BRCC36 (the catalytic subunit)^6,7^, BRE^8,9^ (also known as BABAM2), BABAM1^9–11^ (also known as MERIT40 or NBA1), and RAP80^1,6,12^ (also known as UIMC1)^13^.

The cellular response to DSB (reviewed in ref ^14^) induces the activation of ATM (Ataxia Telangiectasia Mutated) and phosphorylation of histone H2AX (γH2AX) at the damage locus, followed by the recruitment of the E3 ligases RNF8 and RNF168. This results in the ubiquitination of H2A on K13/K15, and the accumulation of Lys63-linked polyubiquitin chains (K63-polyUb) at the damage site^15–18^. This cascade ultimately recruits two key repair factors, 53BP1 and the BRCA1/BARD1 heterodimer, which channel repair towards non-homologous end joining (NHEJ) or homologous recombination (HR), respectively^18–20^. The BRCA1-A complex is recruited to the break site through binding of the tandem ubiquitin interacting motifs (UIMs) of RAP80 to K63-polyUb^1,6,10,12,21–25^, and counteracts excessive accumulation of K63-polyUb deposited by RNF8^26,27^. In response to yet incompletely defined signals, ABRAXAS can be phosphorylated by Ataxia Telangiectasia Mutated (ATM) kinase at Ser404 and Ser406, enabling BRCA1-A to bind the BRCT repeats of BRCA1^2,12,24,28^, thereby sequestering BRCA1 away from the damage site and promoting NHEJ^24,29–31^.

Disruption of BRCA1-A’s DUB activity—for example, by expressing a catalytically inactive BRCC36—leads to elevated HR levels^32^, suggesting that deubiquitination is required for BRCA1-A’s HR-suppressive function. Consistently, BRCC36 knockout or catalytic inactivation sensitizes cells, mice^5^, and zebrafish^33^ to genotoxic agents such as mitomycin C and ionizing radiation, although BRCC36-null or -inactive animals exhibit no overt phenotype under normal conditions. While the physiological targets of BRCA1-A at DSBs remain unclear, it was shown that BRCA1-A engages in auto-deubiquitination, cleaving inhibitory K63-polyUb chains from the N-terminal regions of RAP80 and facilitating its own recruitment at DSBs^5^.

Studies of BRCA1-A are complicated by the existence of a cytoplasmic paralogue of BRCA1-A, BRISC, which comprises BRCC36, BRE, and BABAM1, but features ABRO1 (also known as ABRAXAS2) instead of ABRAXAS, and lacks RAP80^1,8,13,34^. BRISC shares the same super-dimeric architecture and has roles in antiviral response, mitosis, and hematopoiesis (reviewed in ^35^).

*In vitro*, BRCA1-A shows high specificity for K63-linked polyubiquitin chains^36^, with a preference for longer chains^13,36^, and is essentially inactive on mono-ubiquitin (Ub) substrates such as Ub-AMC, requiring at least a diubiquitin (diUb) for activity^13,36^. Although no structure of BRCC36 bound to substrate has been solved, comparison of the apo structures^13,34,37^, with that of the homolog AMSH-LP (also known as STAMBPL1) bound to K63-diUb^23^ show that only K63 linkages can thread into the active site. However, unlike AMSH-LP, BRCA1-A has two active sites positioned such that simultaneous binding of two K63-diUb appears sterically incompatible^37^, raising questions about substrate stoichiometry and mode of engagement.

Unlike most DUBs, which utilise a His-Asp-Cys catalytic triad^38,39^, BRCC36 is a zinc-dependent metalloprotease of the JAB1/MPN/Mov34 metalloenzyme (JAMM) family^9,36,40^. JAMM DUBs use a His-His-Asp triad to chelate a catalytic zinc ion, and a Glu residue to activate a water molecule that hydrolyses the isopeptide bond^35,37,41,42^. A conserved Ser residue stabilises the tetrahedral intermediate during catalysis^42^. In JAMM domain DUBs, substrate positioning is achieved by two characteristic insertions (insertion 1, Ins-1 and insertion 2, Ins-2) compared to the prototype AfJAMM (*Archaeoglobus fulgidus* JAMM)^43^. Ins-1 generally positions the C-terminus of the distal ubiquitin in the active site whereas Ins-2 positions the proximal ubiquitin. In human BRCC36, Ins-1 involves residue 80-93 and Ins-2 residues 175-211.

Activity-based probes^44–47^ have been instrumental in understanding the specificity of cysteine-based DUBs. However, only a mono-ubiquitin activity-based probe for metallo-DUBs has been reported to date^48^, and it is poorly active against BRCC36.

Here, we developed a polyubiquitin-based, metallo-DUB-specific activity probe and used it to determine cryo-EM structures of BRCA1-A in complex with substrate, in order to elucidate the molecular basis of its specificity and catalytic mechanism.

## Results

### BRCA1-A preferentially acts on long polyUb chains

To study how BRCA1-A acts on K63-linked ubiquitin chains, we expressed and purified the human BRCA1-A core complex, consisting of full-length ABRAXAS, BRCC36, BABAM1, and BRE, with RAP80 truncated to residues 265–413 (Fig. 1A). We then profiled its activity against polyubiquitin chains. K63-linked chains were prepared using UBC13/MMS2 and RNF8 as E2 and E3, respectively. Long polyUb chains (Ub_15+_) were isolated by gel filtration and then digested. BRCA1-A-mediated chain processing was monitored over time by following the abundance of individual ubiquitin species (Fig. 1B).

**Figure 1:**
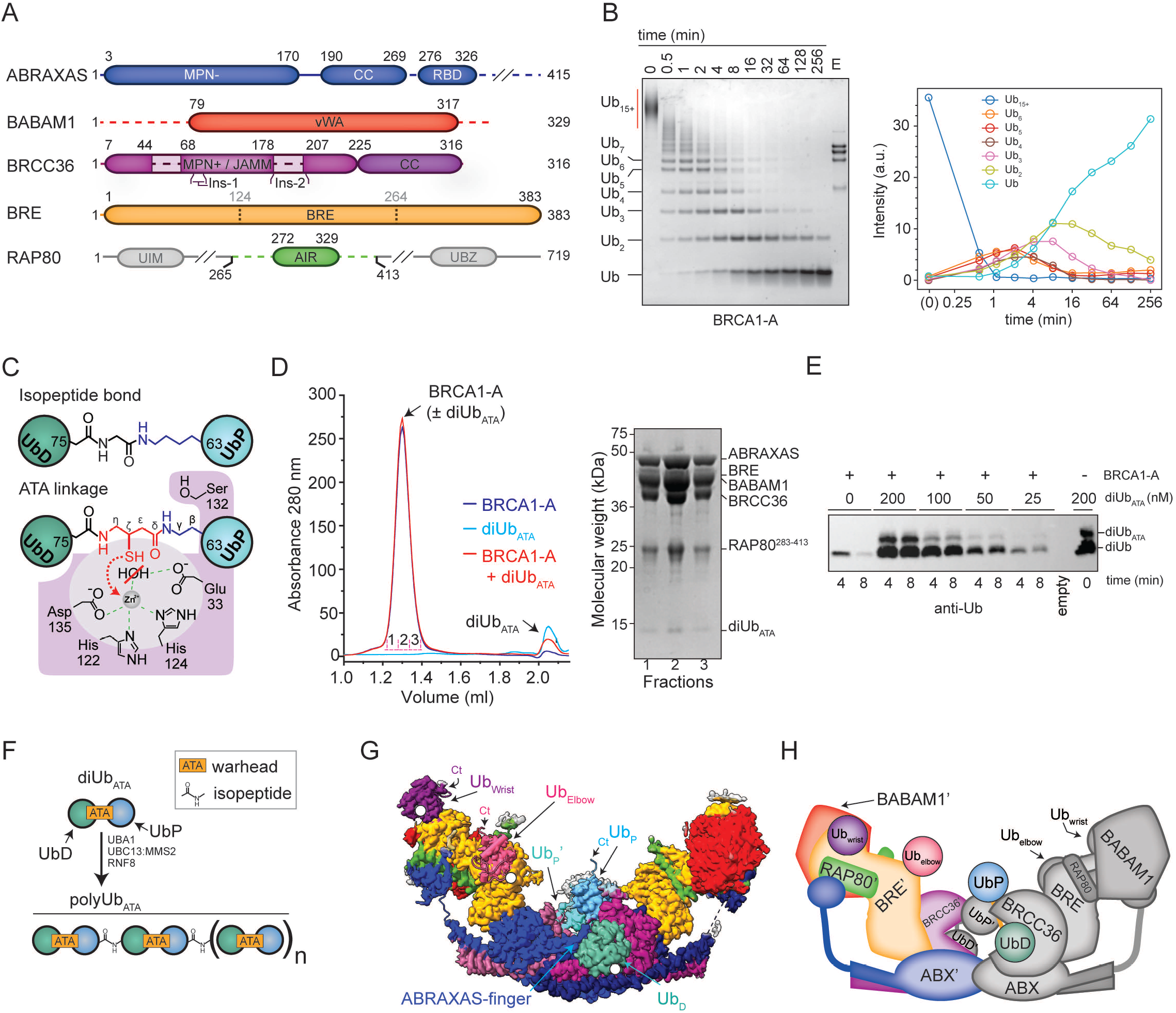
A diUb_ATA_ probe for metallo-DUBs enables analysis of BRCA1-A substrate recognition. **(A) Domain organisation of the BRCA1-A complex components used in this study**. Folded domains are shown as rectangles, and disordered regions as lines, with residue boundaries indicated by numbers. Dashed segments represent regions that could not be traced in the cryo-EM density. Grey segments in RAP80 indicate portions absent from the expression construct. The BRE domain of BRE is formed by the ancestral fusion of three domains from the ubiquitin-conjugating enzyme superfamily^50^, whose approximate boundaries are indicated by dashed lines and grey numbers. Ins-1, Ins-2: JAMM domain insertion-1, -2. **(B) BRCA1-A is more efficient toward long polyUb chains**. Left: long K63-linked polyubiquitin chains (Ub_15+_) were digested with 5 nM BRCA1-A in a time course, and products at the indicated time points were resolved by SDS–PAGE and stained with Coomassie. Five microliters of 1 μM BRCA1-A stock were loaded as a control (lane E). Bands corresponding to various Ub species are indicated on the left of the gel. Right: band intensities corresponding to each species were quantified and plotted. Ub_15+_ levels were quantified by densitometry of the horizontal gel region indicated by the red bar. As long chains (Ub_15+_) are processed, shorter species (Ub_2–6_) accumulate before being further digested in order of decreasing length. Time is shown on a log_2_ scale. A.u., arbitrary intensity units. Time 0 was plotted at 2^-4^ minutes and denoted as (0). **(C) Design of a di-ubiquitin probe for metalloproteases** Top: chemical structure of a native isopeptide bond. Bottom: schematic of the generic active site organisation of a JAMM metallo-DUB. A zinc ion is coordinated by a His–His–Asp triad and, together with a Glu residue, activates a water molecule for isopeptide bond hydrolysis. A Ser residue forms an oxyanion hole that stabilises the reaction intermediate^38^. DiUb_ATA_ shortens the lysine side chain, shifting the isopeptide bond and replacing it with a thiol warhead, which is expected to displace the catalytic water and chelate the zinc ion. DiUb_ATA_ is illustrated with a K63 linkage, with the portions of the warhead mimicking Gly76 and Lys63 coloured red and blue, respectively. ATA carbons are labelled using Greek letters. Residue numbering corresponds to BRCC36. **(D) diUb**_**ATA**_ **binds BRCA1-A**. BRCA1-A was incubated with a five-fold molar excess of diUb_ATA_ and analyzed by size-exclusion chromatography using a Superose 6 Increase 3.2/300 column. BRCA1-A and diUb_ATA_ were run separately as controls. SDS-PAGE of the fractions corresponding to the BRCA1-A:diUb_ATA_ main peak are shown. diUb_ATA_ (17 kDa) co-migrates with BRCA1-A (368 kDa), demonstrating that they form a complex. **(E) diUb**_**ATA**_ **inhibits diUb cleavage by BRCA1-A**. Fifty nanomolar BRCA1-A was used to digest diUb in the presence or absence of diUb_ATA_ for the indicated time points. Reaction products were resolved by SDS-PAGE and visualised by anti Ub western blot. Cleavage of diUb is inhibited in the presence of diUb_ATA_. **(F) Probe chains can be generated enzymatically**. The diUb_ATA_ probe can be assembled in chains using an appropriate E1 E2 E3 combination. K63-linked probe chains used in this study were assembled using UBA1, UBC13:MMS2, and RNF8. These probe chains alternate between an ATA warhead and a native, cleavable isopeptide bond. **(G) CryoEM structure of the BRCA1-A core complex bound to poly-Ub**_**ATA**_ **chains**. A side view of the structure is shown, with sharpened electron density segmented and coloured by subunit (Blue: ABRAXAS; Light purple: BRCC36; Gold: BRE; Red: BABAM1; Green: RAP80; Light grey: unmodelled density). The visible ubiquitin moieties of one probe chain bound to the complex are labelled: UbD (distal ubiquitin, teal), UbP (proximal ubiquitin, light blue), Ub_elbow_ (pink), and Ub_wrist_ (purple). The corresponding Ub moieties for the second probe chain are hidden in this view, except for the proximal ubiquitin (UbP’, cyan). The density for Ub_elbow_ and Ub_wrist_ is weak in the global reconstruction due to arm flexibility but is recovered by focused local refinement. The positions of the C-terminus (Ct) and free Lys63 (white dot) of the bound ubiquitins are indicated. **(H) Schematic representation of the structure**. Core subunits and visible ubiquitin moieties are shown in solid colours matching Fig. 1C for one arm, in grey for the second arm. Subunit of the first arm are denoted with primes. ABX: ABRAXAS.

BRCA1-A cleaves ubiquitin chains at markedly different rates depending on chain length. Long chains (Ub_7+_) are processed rapidly, leading to early accumulation of shorter species (Ub_2–6_). Intermediate chains (Ub_4–6_) are then cleaved, followed by Ub_3_ and finally Ub_2_. A similar length-dependent processing order is observed when shorter chains (Ub_2-5_) are used as substrates (Supplementary Fig. 1A). Apparent half-lives (t_1_/_2_) estimated from the decay phase in Fig. 1B (Supplementary figure 1B) support a tiered substrate preference, with long chains (Ub_15+_, t_1_/_2_ 0.16–0.18 min, 95% confidence interval) processed more rapidly than intermediate-length chains (Ub_5-6_, t_1_/_2_ = 2.4–3.6 min; Ub_4_, t_1_/_2_ = 3.7 - 4.6 min) and short chains (Ub_3_, t_1_/_2_ = 8–29 min; Ub_2_, t_1_/_2_ = 19 – 266 min).

### A new polyUb probe for metallo-DUBs

We wanted to use an activity-based probe to study the catalytic activity and specificity of BRCA1-A, but such probes exist for metallo-DUBs only in the monoUb form^48^. As BRCC36, is active only on diUb or longer chains, we developed a diUb-based probe (diUb_ATA_) by inserting an uncleavable, zinc-chelating thiol warhead between two ubiquitins (Fig. 1C). This was obtained by reacting a Ub where K63 was substituted with 2,4-diaminobutyric acid (UbD_ab_) with a Ub bearing 4-amino-3-thiobutyric acid in position 76 (Ub_ATA_) through native chemical ligation (Supplementary Fig. 1C-E). DiUb_ATA_ can bind to the BRCA1-A complex (Fig. 1D) and inhibits its activity (Fig. 1E).

We also enzymatically polymerized diUb_ATA_ into ubiquitin chains with alternating engineered, uncleavable, and natural cleavable bonds (oligo- and polyUb_ATA_; Fig.1F). Although we used K63 linkages to meet the specificity of BRCC36, diUb_ATA_ can be synthesised with any lysine linkage and polymerised into all chain types with known E2s, offering a versatile tool for probing the activity and specificity of other JAMM-family metallo-DUBs.

### CryoEM structure of BRCA1-A in complex with probe chains

In order to understand how BRCA1-A gains its specificity for K63-polyUb, we determined cryo-EM structures of the human BRCA1-A core complex alone, in complex with diUb_ATA_, oligoUb_ATA_ (Ub_4_-Ub_8_) and polyUb_ATA_ (Ub_6_-Ub_12+_) (Supplementary Table 1). Collectively, these structures offer a coherent view of BRCA1-A’s catalytic mechanism, capturing multiple conformational and substrate-bound states distinguishable by classification.

The structure of BRCA1-A bound to polyUb_ATA_ (BRCA1-A:polyUb_ATA_ State C+P, 3.2 Å, Supplementary Table 1) provides the most complete picture and will be described first. In this state, the BRCA1-A core complex adopts a two-armed, dimer-of-pentamers architecture that closely resembles the previously solved crystal structure (Fig. 1G and 1H, Supplementary Fig. 1F). No density is observed for the N-terminus of BABAM1 (residues 1– 88) or the C-terminus of ABRAXAS (residues 327–405), indicating that these regions are disordered (Fig. 1A).

The substrate chains appear partially disordered, with eight ubiquitin moieties visible: a diUb_ATA_ in each active site and two single ubiquitins per arm. Following standard nomenclature, we refer to the diUb_ATA_ components in the active site as distal ubiquitin (UbD), and proximal ubiquitin (UbP), with the C-terminus of UbD connecting to position 63 of UbP via the ATA probe. We term the ubiquitins binding to BRE as “wrist ubiquitin” (Ub_wrist_) and “elbow ubiquitin” (Ub_elbow_) based on their relative positions on the BRCA1-A arm (Fig. 1G). These sites are likely the anchor points for two partially flexible polyUb_ATA_ chains, with each chain binding at wrist and elbow of one arm before entering the active site at the base of the opposite arm.

### Active sites bind simultaneously, with alternate conformations

As a super-dimer, BRCA1-A has two active sites, which surprisingly bind diUb_ATA_ simultaneously, but in distinct conformations (Fig. 2A, 2B): one adopts a catalytically competent arrangement (“State C”), while the diUb_ATA_ in the opposing site is not fully engaged, representing what is likely a precatalytic configuration (“State P”). In state C, the BRCC36 active site is correctly aligned for catalysis (Fig. 2C, Supplementary Fig.2A), with the catalytic zinc coordinated by His122, His124, and Asp135. The ATA probe, which substitutes for Gly76 of UbD and Lys63 of UbP, is positioned directly in front of the active site, and coordinates the catalytic zinc through its thiol (Fig. 2C, Supplementary Fig. 2A). UbD engages BRCC36 via its Ile44 hydrophobic patch (Leu8, Ile44, His68, Val70) and threads its C-terminus into the active site, where it forms a β-sheet with Ins-1 of BRCC36. Leu71 and Leu73 of UbD interact with the hydrophobic core of the active site, while Arg72 and Arg74 point outwards, forming a basic patch alongside Arg42. UbP engages BRCC36 through an unusual hydrophobic interface centred on Met1, Glu18, Pro19, and Ser20. Additionally, two key interactions with Ins-1 stabilize UbP binding: Gln2 of UbP interacts with Lys87 and Asp88 of BRCC36, and Glu64 of UbP connects with His133 and Arg89 of BRCC36 (Fig. 2A). Together, UbP Glu64 and BRCC36 Arg89 act as a lid that seals the reaction chamber, likely shielding the catalytic water required to hydrolyse the isopeptide bond under *wild-type* conditions. Comparison of State C with the structure of inactive AMSH-LP (BRCC36’s closest homolog) in complex with K63-diUb shows that the ATA probe sterically mimics the native isopeptide bond (Supplementary Fig. 2A).

**Figure 2:**
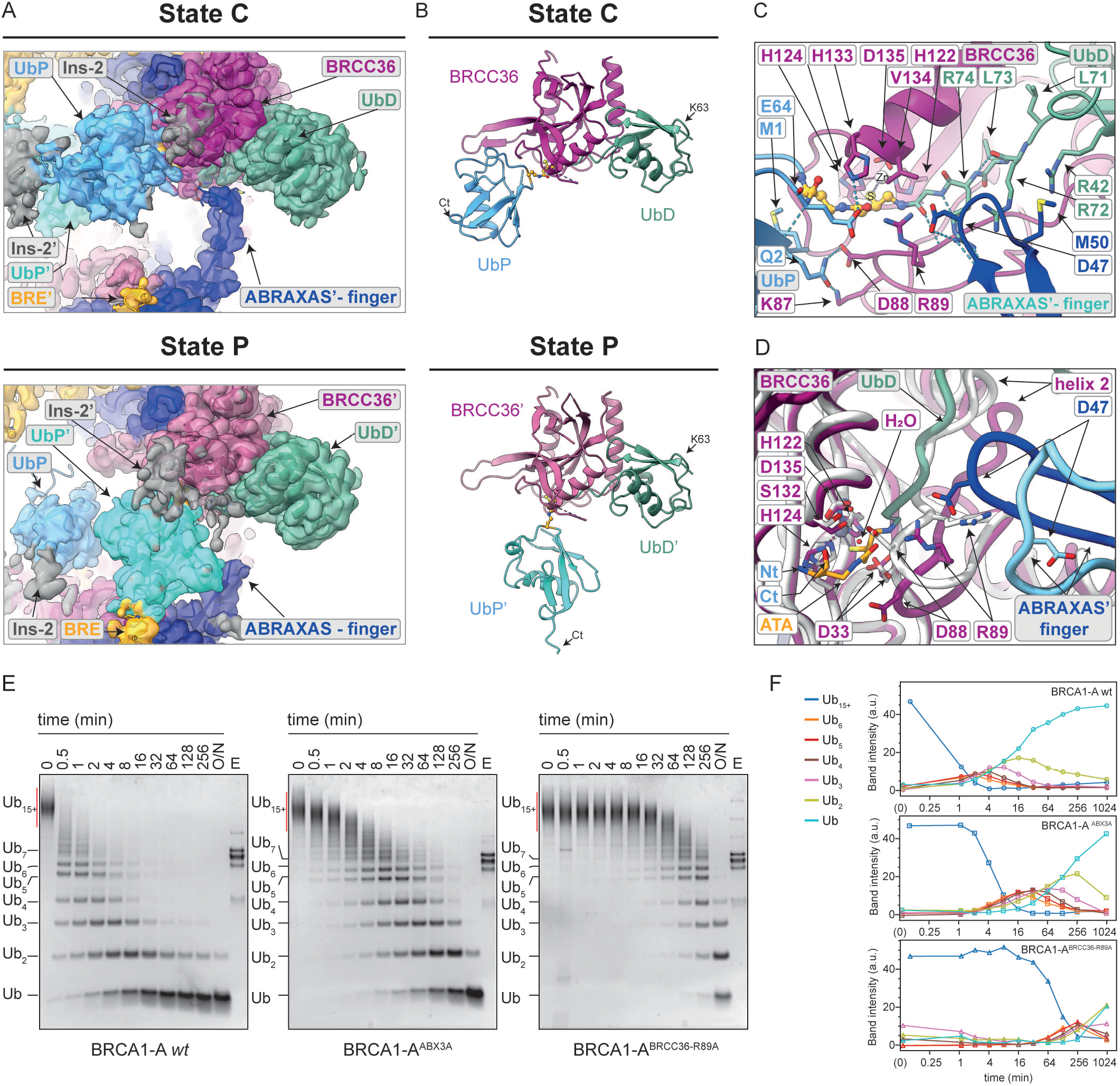
ABRAXAS facilitates catalysis in trans. **(A) The two active sites of BRCA1-A bind diUb**_**ATA**_ **in different conformations**. The two active sites in BRCA1-A:PolyUb_ATA_ (State C+P, Supplementary Table 1) are shown, rotated to provide the same orientation. Sharpened cryo-EM density is shown, in the top panel for a state where diUb_ATA_ is bound in a catalytically competent conformation (State C); in the bottom panel, the other active site is shown, displaying a pre-catalytic conformation (State P), in which proximal UbP’ is displaced from the active site. Subunits of the pentamer containing BRCC36 in State P and the chain bound to it are denoted with a prime (⍰), e.g. BRE’, UbP’. Residues 44-52 of the ABRAXAS finger (opposite State P BCC36) are disordered. Density is coloured by subunit: BRCC36, purple; UbD, teal; UbP, light blue; UbP’: cyan; ABRAXAS, blue; BRE, gold. Unmodelled electron density corresponding to BRCC36 Ins-2 is shown in grey. **(B) State C and State P differ in the conformation of UbP**. BRCC36 and the bound diUb_ATA_ from each active site in the BRCA1-A:polyUb_ATA_ structure (State C, top; State P, bottom) are shown in cartoon representation and in the same orientation. While BRCC36 and UbD adopt similar conformations in both states, the position of proximal UbP differs between State C and State P. Colours as in A. Ct: C-terminus. **(C). Ins-1 positions both Ub moieties of the substrate and seals the reaction chamber**. Cartoon representation of the active site in State C is shown, where BRCC36 Ins-1 (residues 80-93) positions the substrate, with Arg89 sealing the reaction chamber. The ABRAXAS-finger contacts BRCC36 Arg89 and the basic patch formed by arginines 46, 72 and 74 of UbP. BRCC36 Lys87, Asp88 and His133 position UbP by interacting with UbP Gln2 and Glu64. Key residues are displayed as sticks and labelled. Colours as in A. **(D) The ABRAXAS-finger coordinates with Ins-1 during substrate engagement**. Superposition of apo BRCC36 (white) and polyUb_ATA_-bound BRCC36 (purple, State C) reveals that Ins-1 shifts downward to accommodate the C-terminus of UbD (teal), displacing BRCC36 Asp88 from its position covering the active site^42,50^. The ABRAXAS-finger (light blue in the apo structure; blue in State C) interacts with Arg89 of Ins-1 and stabilises this conformational transition. H_2_O indicates the catalytic water present in the apo structure. UbP residues connecting to the amino (Nt) and carboxyl (Ct) groups of the ATA warhead are omitted for clarity. **(E) The ABRAXAS-finger and Arg89 of BRCC36 are necessary for efficient catalysis**. Ubiquitin chain cleavage assay where 5 nM wild type BRCA1-A, BRCA1-A^ABX3A^ or BRCA1-A^BRCC36 R89A^ were incubated with K63-linked polyubiquitin chains at 37°C. Aliquots were collected at the indicated time points, resolved by SDS-PAGE, and stained with Coomassie blue. E: five microliters of 1 µM BRCA1-A stock were loaded as control. O/N: overnight. **(F) Kinetics of digestion of the ubiquitin species in D**. Band intensities corresponding to the different ubiquitin species in (E) were quantified by densitometry and plotted over time. For Disappearance of Ub _15+_ and peak maxima of Ub_2-6_ for BRCA1-A^ABX3A^ (red) and BRCA1-A^BRCC36 R89A^ (blue) are delayed by approximately 4 and 7–8 time points compared to *wt* BRCA1-A (black), corresponding to ~2^4^ (16-fold) and ~2^7^–2^8^ (128–256-fold) reductions in catalytic efficiency, respectively. Ub_15+_ levels were quantified by densitometry of the horizontal gel region indicated by the red bar. Time is shown on a log_2_ scale. A.u., arbitrary intensity units; O/N: overnight. Time 0 was plotted at 2-4 minutes and denoted as (0). Overnight digestion was approximated to 1024 min (~17 h).

### ABRAXAS facilitates catalysis *in trans*

Interestingly, in State C, a loop of ABRAXAS (residues 39–57, hereafter referred to as the “ABRAXAS-finger”) extends *in trans* from the opposite pentamer of the complex. At its tip, a negatively charged Thr-Asp-Ser motif (residues 46-48) interacts with Arg89 of Ins-1 in BRCC36, and with Arg72 and Arg74 of UbD (Fig. 1G, 2A).

The ABRAXAS-finger undergoes a striking conformational change during substrate engagement. In our apo BRCA1-A structure (“Apo”, 3.2 Å, Supplementary Table 1) and in the mouse BRCA1-A crystal structure^13^, it rests against Ins-1 of BRCC36, with Glu88 from Ins-1 coordinating the catalytic water, and Arg89 oriented outward (Fig. 2D). Upon substrate binding, UbD entry pushes helix 2 and Ins-1 of BRCC36 downward, opening the active site. The ABRAXAS-finger then extends forward to stabilise this new configuration, holding Ins-1 down, positioning Arg89 over the catalytic centre, and securing UbD’s C-terminus within the active site. Upon substrate binding, engagement of the ABRAXAS-finger with Ins-1 shortens the distance between the MPN domains of opposing ABRAXAS and BRCC36, resulting in partial arm closure and slight arching of the coiled coil in BRCA1-A (Supplementary movie 1, Supplementary Fig.2B).

Notably, both the length of the ABRAXAS-finger and the Asp residue at its tip are strictly conserved across evolution for ABRAXAS (present in metazoans) and its paralogue ABRO1 (present in all eukaryotes; Supplementary Fig.2C). Furthermore, BRCC36 Arg89 is invariant in eukaryotes (Supplementary Fig.2D), and among JAMM metallo-DUBs, only BRCC36 bears an arginine at this position, suggesting co-evolution of BRCC36 Arg89 and ABRAXAS/ABRO1 Asp47 and supporting an important, conserved structural role for the ABRAXAS-finger in regulating active site configuration.

We therefore tested whether the ABRAXAS-finger allosterically regulates BRCC36 DUB activity. To this end, we generated a mutant in which residues 46–TDS–48 of ABRAXAS were mutated to alanines (BRCA1-A^ABX3A^) and assessed its ability to cleave K63-linked polyUb chains (Fig. 2E). BRCA1-A^ABX3A^ exhibited the same length-dependent substrate processing order observed for *wild type* BRCA1-A, but showed an approximately 16-fold reduction in catalytic activity (Fig. 2E, 2F). Mutation of Arg 89 of BRCC36 to alanine (BRCA1-A^BRCC36-R89A^) caused an even stronger impairment (~128-256-fold), consistent with a dual role for Arg89 in engaging the ABRAXAS finger and in shielding the reaction chamber (Fig. 2D). Together, these mutations indicate an essential role of the ABRAXAS-finger in catalysis.

Notably, since ABRAXAS is exclusive to BRCA1-A, BRCA1-A^ABX3A^ may serve as a useful separation-of-function mutant to distinguish BRCA1-A activity from its paralog BRISC, while the equivalent ABRO1 mutation could report on BRISC activity.

### Active sites engage simultaneously, but alternate in cleavage

The second active site of BRCA1-A binds a diUb_ATA_ molecule in a different configuration (State P, Fig. 3A). Subunits from the pentamer in State P, and the diUb_ATA_ bound to it, are hereafter denoted with primes. In state P, diUb_ATA_’ adopts a bent conformation: UbD’, its C-terminus, and the ATA’ probe until the thiol-bearing carbon (corresponding to Gly76 of UbD’ and the mimetic scissile bond), retain the conformation observed in state C. Beyond the thiol, however, ATA’ bends by ~53°, allowing UbP’ to extend towards the arm opposite the active site, where its C-terminus interacts with Trp165 of BRE (Fig. 3A; Supplementary Fig. 3A). Additional interactions between UbP’ and BRCC36’ involve Gln62 of UbP’ contacting His133 and Arg208 of BRCC36’ (Figure 3A).

**Figure 3:**
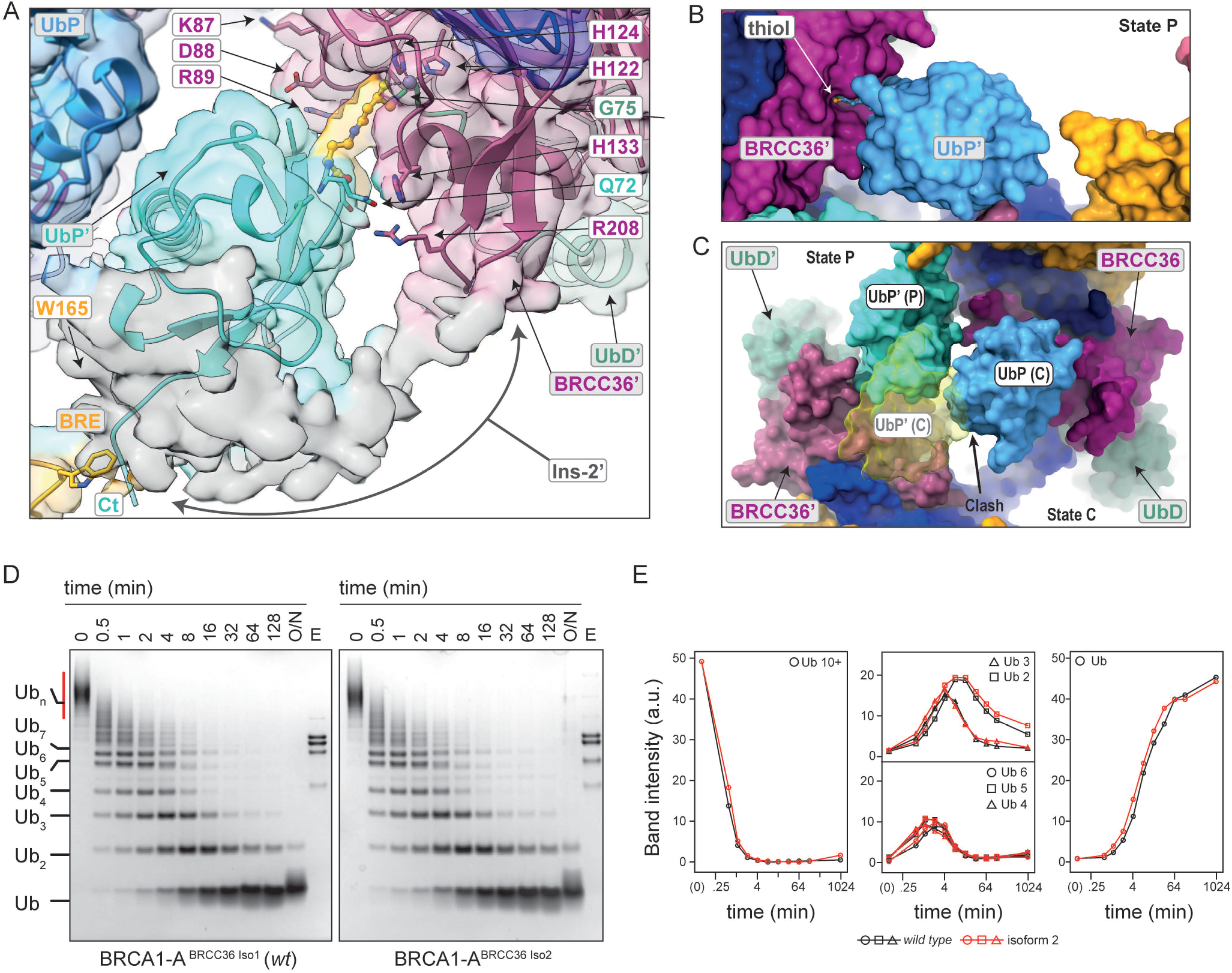
Active sites cleave in alternation. **(A) In stateP, UbP is displaced towards BRE and interacts with Ins-2**. Sharpened electron density and fitted model of the BRCA1-A:PolyUb_ATA_ (State C+P) structure, focused on the active site in state P. Although BRCC36’ (purple) and UbD’ (teal) are in the same conformation as state C, UbP’ (light blue) is displaced towards BRE (gold), dragging the probe (gold, ball representation) out of the active site. Density for BRCC36’ Ins-2 (grey, unmodelled) overlays UbP’ and the UbP’:BRE interface. **(B) State P is likely cleavage incompetent**. A *s*pace filling representation of the active site in State P is shown, viewed along the axis of the UbD C-terminus. Pulling of UbP towards BRE on the opposite arm leaves the thiol of ATA (mimicking the catalytic water) exposed to bulk solvent. In native conditions, positioning and chemical activation of the catalytic water would likely be perturbed despite the presence of a properly aligned active site. Colours as in A. **(C) Active sites cannot be simultaneously in state C**. The two active sites in the BRCA1-A:PolyUb_ATA_ (State C+P) structure are shown, with BRCC36 (State C) on the right and BRCC36’ (State P) on the left. A yellow cartoon indicates the position of UbP’ if the left active site were also in State C, demonstrating that it would sterically clash with the opposite UbP. Representation and colouring are as in (A). **(D) A longer Ins-2 is not necessary for catalysis**. BRCA1-A^iso2^, assembled with BRCC36 isoform 2, bearing a shorter Ins-2, was assayed for K63-Ub chains cleavage and exhibits catalytic activity comparable to BRCA1-A^wt^. **(E) BRCA1-A activity is not influences by BRCC36 isoform choice**. Band intensities corresponding to the different ubiquitin species in (D) were quantified by densitometry and plotted over time. No difference is apparent between BRCA1-A bearing BRCC36 isoform 1 (*wild* type, black) or isoform 2 (red). Ub_10+_ levels were quantified by densitometry of the horizontal gel region indicated by the red bar. Time is shown on a log_2_ scale. A.u., arbitrary intensity units; O/N: overnight. Time 0 was plotted at 2-4 minutes and denoted as (0). Overnight digestion was approximated to 1024 min (~17 h).

State P appears to be cleavage incompetent. Bending of ATA’ pushes BRCC36’ Arg89 towards the solvent (Fig. 3B), and the ABRAXAS-finger remains disordered, or, in a subset of the population, interacts with the C-terminus of UbD’ (Fig. 2A, Supplementary Fig.3B). In addition, the entire backside of the reaction chamber is exposed to bulk solvent, likely interfering with activation of the catalytic water (mimicked here by the thiol of ATA) by the zinc ion (Fig. 3B).

For State P to become catalytically competent, UbP⍰ would need to rotate and engage BRCC36’. However, this is sterically incompatible with the presence of UbP in the opposing state C active site (Fig. 3D). This suggests that, for an active site to transition from State P to State C, UbP must first be cleaved and released from the opposing site. Consistent with this, while we observed a configuration in which both active sites adopt State P (BRCA1-A:polyUb_ATA_ State double P, 3.1 Å, Supplementary Table 1, Supplementary Fig. 3C), we never observed a structure in which both were simultaneously in State C. Therefore, although both active sites of BRCA1-A can engage substrate at the same time, cleavage seems only possible in alternation (Supplementary movie 2).

### Extended Insertion 2 of BRCC36 is dispensable for catalysis

In the BRCC36 homologs AMSH and AMSH-LP, UbP is positioned by Ins-2, and Ins-2 also contributes a phenylalanine residue to form the lid of the reaction chamber^42^. Our structures show that in BRCC36, Ins-1 can fulfil the functions that in AMSH and AMSH-LP are fulfilled by Ins-2, namely UbP recognition (via Lys87 and Asp 88) and sealing of the reaction chamber (via Arg 89). Meanwhile, density for Ins-2 – while too weak overall to allow modelling – is visible above UbP in both state C and state P (Fig. 3A), suggesting BRCC36 Ins-2 might still play some role in UbP recognition.

In most placental mammals (mouse being a notable exception, Supplementary Fig. 3D and 3E), BRCC36 has two splicing isoforms: the canonical isoform 1, used in this work, with a long Ins-2 (residues 175-211), and an alternative splicing isoform 2, where Ins-2 is shortened by the deletion of residues 184-208 and cannot physically reach UbP. We compared the chain cleavage activity of BRCA1-A complexes assembled with either BRCC36 isoform 1 (BRCA1-A^*WT*^) or isoform 2 (BRCA1-A^iso2^). When Ins-2 cannot contact UbP (BRCA1-A^iso2^) we observe no difference in catalytic activity (Fig. 3D, 3E), demonstrating that this contact in Ins-2 is not required for UbP recognition or efficient catalysis *in vitro*.

### BRCA1-A binds ubiquitin chains through the arms

BRCA1-A displays a clear preference for ubiquitin chains as substrates compared to diUb^13^ (Fig. 1B), suggesting that ubiquitin binding sites must exist beyond the active site. Consistent with this, the BABAM1:BRE complex was shown to bind ubiquitin^35,49^. In our BRCA1-A:polyUb_ATA_ structure, two Ub binding sites on BRE are revealed on the arm, at the wrist and elbow positions (Fig. 1G).

At the elbow, Ub engages BRE through its canonical Ile44 hydrophobic patch, which contacts a hydrophobic pocket formed between helices 4 and 5 of BRE, centred on Arg137 and Phe140 (Fig. 4A). At the wrist, ubiquitin binds via an interface involving the tip of ubiquitin helix 1 and the loop connecting to the third β-strand (residues 31–37). On the BRE side, residues 350–SPRW–353 constitute the primary interaction surface (Fig. 4B). The key BRE residues involved in both the elbow and wrist interactions are conserved across eukaryotes (Supplementary Fig. 4A and B). BRE derives from the ancestral fusion of three ancestral ubiquitin conjugating enzyme folds^50^ (Fig. 1A). Neither the elbow nor the wrist ubiquitin-binding sites correspond to those used by active E2 enzymes to engage ubiquitin.

**Figure 4:**
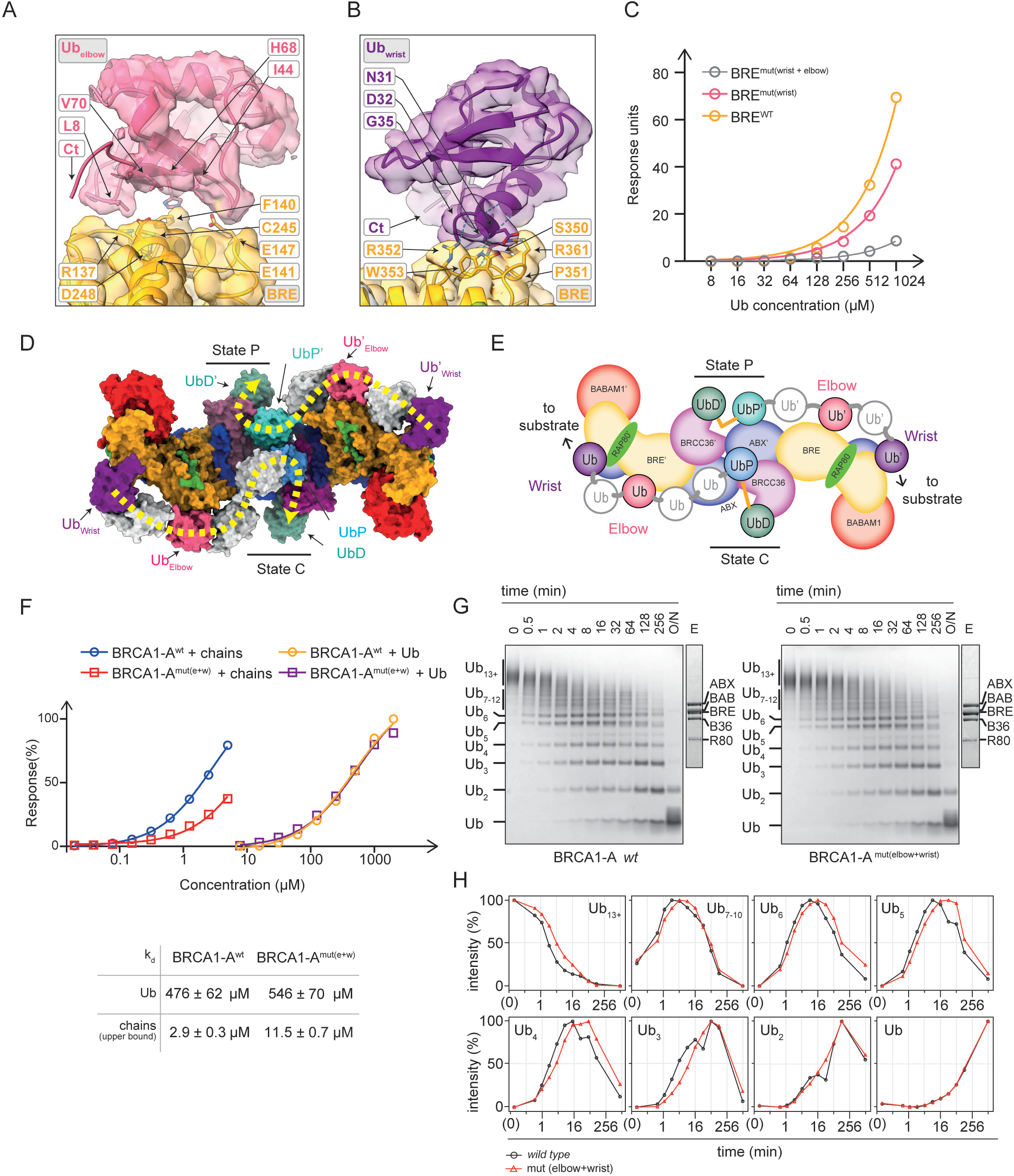
BRCA1-a recognizes ubiquitin chains through avidity. **(A) Ub**_**elbow**_ **binds via Ile44 face**. Particles from the BRCA1-A:polyUb_ATA_ (State C+P) and BRCA1-A:poluUb_ATA_ (double State P) structures were combined, and 3D classification was performed to enrich for particles bearing Ub_elbow_, followed by local refinement. Models for BRE and Ub were fitted as rigid bodies. Sharpened electron density and cartoon representation of the interface between Ub_elbow_ (pink) and BRE (gold) are shown, with key interacting residues highlighted. Map deposited as EMDB: EMD-55052. **(B) Ub**_**wrist**_ **binds via a novel interface centred on G35**. As in (A), particles were classified and refined with a focus on Ub_wrist_. Models for BRE and Ub were first fitted as rigid bodies, then real-space refined in Phenix^68^. The interface between BRE and Ub_wrist_ is shown. Map and model deposited as PDB: 9SMS and EMDB: EMD-55041. **(C) The BRE wrist site accounts for most of the binding affinity of BRE for ubiquitin**. GST-BRE, GST-BRE^mut(wrist)^ or GST-BRE^mut(wrist+elbow)^ were produced recombinantly in *E. coli*, immobilized through anti-GST antibody and assayed for their association with monoUb by surface plasmon resonance (SPR). Although the data do not allow robust fitting of a dissociation constant (k_d_), both mutants display reduced affinity. **(D) Ubiquitin chains require a minimal length of five moieties to span the arms of BRCA1-A**. The minimal number of ubiquitin moieties (white) required to connect Ub_wrist_ and Ub_elbow_ to UbP was modelled on the polyUb_ATA_ State C+P structure (colour). At least five ubiquitins are needed to span the shortest path between the wrist and the active site in State P, and six are required for the active site in State C. Each chain binds at the wrist and elbow of one arm and feeds into the active site at the base of the opposite arm. **(E) Schematic representation of the structure in D**. Each chain binds along one arm and feeds into the BRCC36 active site of the opposite arm. Accordingly, the chain bound to BRCC36’ in State P is denoted with primes (e.g. UbP’, UbD’, etc.) and engages BRE from the non-prime pentamer containing BRCC36 (in State C), and vice versa. **(F) BRCA1-A binds chains through avidity**. The dissociation constant (k_d_) of His-tagged BRCA1-A^wt^ or BRCA1-A^mut(elbow+wrist)^ for monoUb or K63-linked polyubiquitin chains were measured by SPR (immobilization: anti His antibody). Affinities are reported as K_d_ ± standard error (SE). The k_d_ reported for polyUb chains is based on an estimated chain concentration of ≤5 μM (see Materials and Methods) and therefore represents an upper-bound value. **(G) Avidity mutants are impaired in catalysis**. The activity of BRCA1-A WT or BRCA1-A^mut(elbow+wrist)^ against K63-polyubiquitin chains was assayed in a time course and resolved via SDS PAGE. BRCA1-A was used at 1 nM. Lane E: 5 μL of the 1 μM enzyme stocks used in the experiment were loaded as a control on a separate gel (shared for both BRCA1-A variants). O/N: overnight. **(H) BRCA1-A** ^**mut(elbow+wrist)**^ **displays a 2-4 fold catalysis defect**.. Band intensities corresponding to the different ubiquitin species in (D) were quantified by densitometry and plotted over time. For all species except Ub_2_ peak maxima for BRCA1-A^mut(elbow+wrist)^ (red) are delayed by one to two time points compared to wild type BRCA1-A (black), corresponding to ~2-4-fold catalytic defect. Time is shown on a log_2_ scale. A.u., arbitrary intensity units; Overnight digestion was assumed to correspond to 1024 min (~17 h). All curves were independently normalised to their respective minima and maxima (scaled between 0 and 1).

To validate these binding sites, we expressed BRE in *E. coli* as a GST fusion protein to measure Ub binding affinity using surface plasmon resonance (SPR, Fig. 4C and Supplementary Fig. 4C). BRE binds Ub with low affinity (estimated k_d_ ≈ 850 μM). Disruption of the wrist site by mutation of BRE residues 350-SPRW-353 to four alanines (BRE^mut(wrist)^) reduces the affinity (estimated > 3000 μM). Further mutation of the elbow site (R137D F140A, BRE^mut(elbow+wrist)^) essentially abolishes the interaction of BRE with Ub. Together, these data validate the observed binding sites and indicate that the majority of BRE’s affinity for ubiquitin originates from the wrist site. Consistent with this, in a structure of BRCA1-A bound to diUb_ATA_ (BRCA1-A:diUb_ATA_, 3.1 Å, Supplementary Table 1, Supplementary Figure 4D), we observed an ordered ubiquitin moiety bound at each wrist site, but no binding at the elbow (Supplementary Fig. 4D).

Since our polyUb_ATA_ is uncleavable and does not contain detectable monoUb species (Supplementary Fig. 4E), the ubiquitins we observe bound to BRE must be parts of disordered longer chains. Thus, we interpret Ub_wrist_ and Ub_elbow_ as anchoring points for two probe chains. The C-terminal residues (72-76) and the side chain of K63 are disordered in both Ub_wrist_ and Ub_elbow_, indicating conformational flexibility. Nevertheless, their positioning indicates that each chain runs along an arm of the complex and feeds into the active site at the base of the opposite arm, where it appears as diUb_ATA_. The elbow and wrist ubiquitin connecting to diUb_ATA_’ and BRCC36’ are hereafter marked with primes (e.g. Ub_elbow_’). The ubiquitin moieties connecting the visible molecules at the active site and the wrist via the elbow are conformationally flexible and not resolved in our cryo-EM maps. Assuming an extended K63-linked polyubiquitin conformation (~70 Å per moiety), one ubiquitin appears sufficient to bridge Ub_wrist_ and Ub_elbow_, and one or two are required to connect Ub_elbow_ to the diUb in the active site, depending on the active site conformation (State P or State C, Fig. 3D and 3E). Thus, Ub_6_ (State P) and Ub_7_ (State C) represent the minimal chain lengths that can span all ubiquitin-binding sites; Ub_4_/Ub_5_ can engage only the elbow site, whereas Ub_3_ and Ub_2_ can reach neither.

### BRCA1-A binds long Ub chains through avidity

Ubiquitin-binding proteins that specialize in chain recognition typically exploit avidity, i.e., multivalent interactions, where multiple low-affinity binding sites engage different ubiquitin moieties within the chain (reviewed in ref ^51^). The presence of low-affinity ubiquitin-binding sites along the arm suggests that BRCA1-A follows a similar strategy. To test this, we compared the affinity of our BRCA1-A (lacking the UIMs of RAP80) for monoUb and *wild type* polyUb chains using SPR (Fig. 4F and Supplementary Fig. 4F). BRCA1-A binds monoUb with an affinity of 476 ± 62 μM. The chains vary in length, precluding an exact determination of their concentration. If we estimate an upper-bound concentration of 5 μM for the chains (see Materials and Methods and Supplementary Fig. 4G), BRCA1-A has at minimum an affinity of 2.9 ± 0.3 μM for polyubiquitin chains (Fig. 4F), i.e. at least 100-fold higher than that of monoUb. A BRCA1-A variant in which the elbow and wrist sites on BRE were mutated (BRCA1-A^mut(elbow+wrist)^) exhibits a ~5-fold lower apparent affinity 11.5 ± 0.7 for polyUb compared to wild-type while retaining similar affinity for monoUb (546 ± 70 μM, Fig. 4F). Taken together these data shows that BRCA1-A exploits avidity to bind chains, and that the sites on the arms contribute to the effect. Avidity-driven substrate selection appears to explain the three-tiered cleavage kinetics observed in Fig. 1B. Chains longer than Ub_6–7_ fully engage avidity and are preferentially bound and processed, leading to early accumulation of Ub_5–6_. Intermediate chains (Ub_4–6_), which engage avidity only partially via the elbow site, are then processed. Short chains (Ub_2–3_), which interact only with the active site, are cleaved last.

To evaluate the role of avidity in BRCA1-A catalysis, we tested the avidity mutant for chain cleavage. Compared to wild-type BRCA1-A, BRCA1-A^mut(elbow+wrist)^ exhibits a modest but reproducible 2-to 4-fold overall delay in the digestion of long ubiquitin chains (Ub_15+_, Fig. 4G), which translates to a delay in appearance and processing of shorter substrates (Ub_3-10_, Fig. 4G, compare peak maxima). When shorter chains (Ub_2-5_) are used as substrate, a similar delay for BRCA1-A^mut(elbow+wrist)^ is observed with Ub _4,5_ but not Ub _2,3_(Supplementary Fig. 4H,4I). Based on the structure, a length of Ub_4-5_ is required to engage the elbow and the active site, resulting in partial avidity and moderate affinity. Lengths of Ub_6-_ or greater are required to reach the wrist site, enabling full avidity and highest affinity. This mirrors the three-tiered substrate half-lives observed in Fig. 1B (t_1/2_ Ub_15+_ < t_1/2_ Ub_4–6_ << t_1/2_ Ub_3_, Ub_2_). Thus, the substrate preference of BRCA1-A is likely driven by avidity-based recognition of longer over intermediate and short chains.

The elbow and wrist binding sites do not contact the C-terminus or Lys63 of the bound ubiquitin, indicating that they are not intrinsically K63 linkage-specific. However, most other diUb linkage types, with the exception of K27 and M1, appear sterically hindered from engaging at one or both sites. Thus, the arm as a whole appears capable of displaying avidity only toward some chain types, reducing the likelihood of off-target binding or feeding incompatible substrates into the active site. The SIM/UIM domains of RAP80 that are absent from our constructs, may provide an additional layer of regulation and influence avidity, although they have been reported not to robustly affect cleavage of long SUMO-conjugated ubiquitin chains^13^.

### BRCA1-A exists in an equilibrium between open and closed form

Across all reconstructions, independently of substrate length or occupancy in the active sites, we observed variability in the angle between the arms of the BRCA1-A complex. Thus far, we have described the open form, in which the arms are splayed apart. However, in most datasets, BRCA1-A also adopts a closed conformation, where the arms fold inward (Fig. 5A). These closed states form a conformational ensemble rather than a single well-defined structure, resulting in blurred 3D reconstructions. Nevertheless, we applied focused refinement and 3D classification to the closed form, enabling reconstruction of a closed conformation of the complex between BRCA1-A and oligoUb_ATA_ (BRCA1-A:oligoUb_ATA_, 3.4 Å, Supplementary Table 1, Fig. 5B) that bears sufficient detail to build a pseudo-atomic model. The model was built by flexible fitting of an Alphafold3 (AF3) prediction of BRCA1-A (deposited in ModelArchive^52^ with ID: ma-yb0lm, Supplementary Fig. 5A), followed by manual rebuilding in the dimerization interface area, which is better defined (Supplementary Fig. 5B). Interestingly, AF3 consistently predicts human BRCA1-A in a closed form with arms even more closed than in our closed structure. The AF3 prediction fits the density of other lower-resolution subclasses of this closed form (Supplementary Fig. 5A), implying a broad range of possible arm movements. The transition from open to closed form appears to be a natural extension of the movement triggered by substrate engagement (Fig. 5C, Supplementary Movie 3), in which the ABRAXAS-finger advances the JAMM domain of ABRAXAS toward the opposing BRCC36. This motion rotates the BRE elbow forward and inward, pulling upward on the C-terminal end of the coiled coil through the ABRAXAS– RAP80–BABAM1 connection at the wrist (Fig. 5D). As the coiled coil rises, it tugs on ABRAXAS helix 3, which loops back to the MPN domain—thereby completing a rope-and-pulley mechanism entirely driven by ABRAXAS. Strain within the BRCC36 coiled coil is absorbed by two conserved proline residues (Pro236 and Pro282), which allow the segments flanking the catalytic domain to bend in different directions (Fig. 5D). Additionally, the forward shift of the ABRAXAS JAMM domain from one arm is met with compression of loosely packed residues of the opposing ABRAXAS helix 3 and BRCC36 coiled coil at the dimerization interface (Supplementary Fig. 5C). Thus, the dimerisation interface appears to act as a buffer to absorb conformational strain, while mediating coupling between the movement of one arm and the state of the opposing active site.

**Figure 5:**
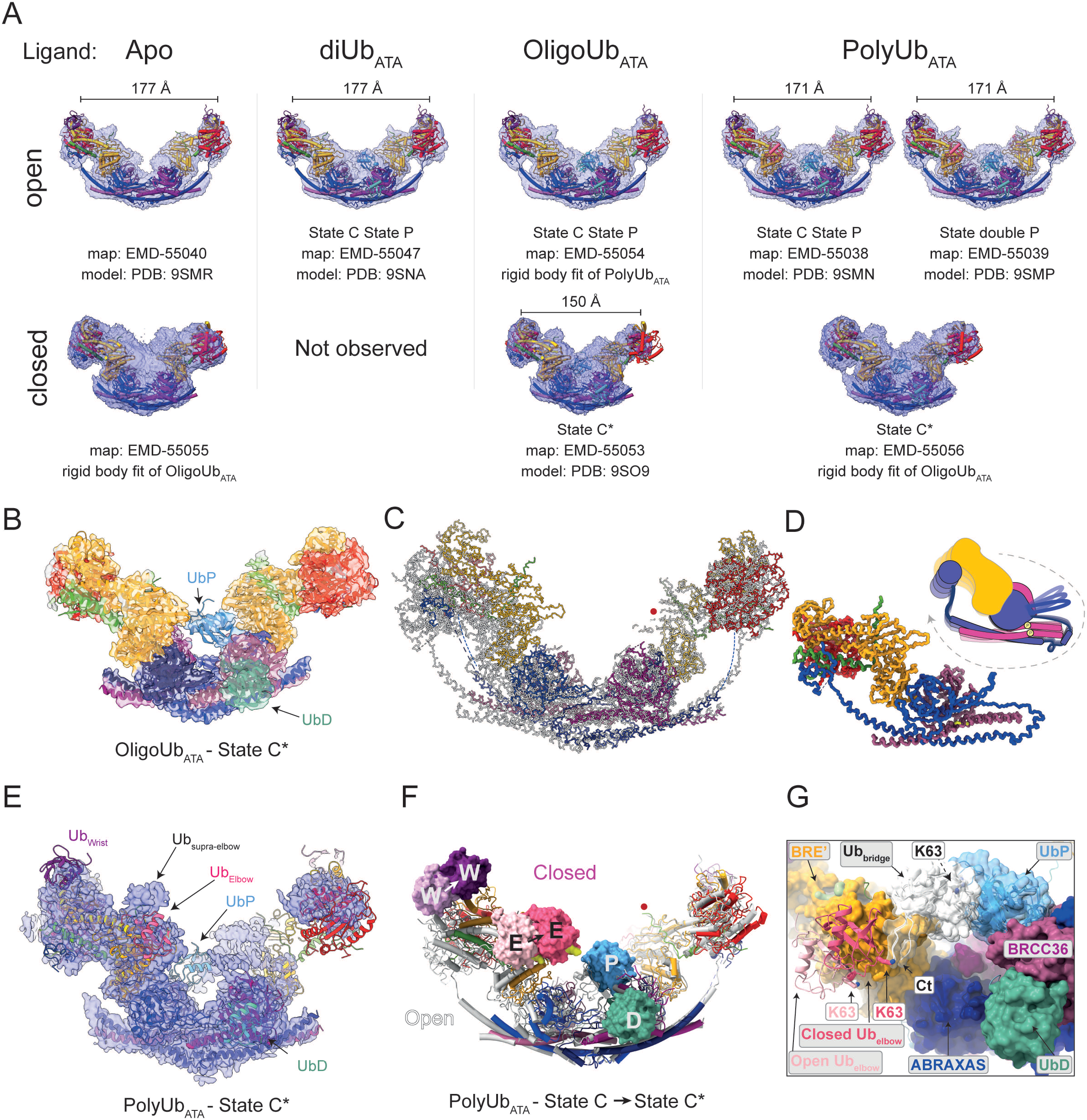
BRCA1-A shifts between open and closed forms to advance ubiquitin. **(A) BRCA1-A adopts both open and closed conformations**. Raw cryo-EM maps and fitted models are shown for the open (top) and closed (bottom) forms, determined in the presence of the indicated substrate (see also Supplementary Table 1 and Supplementary EM information). In most datasets—regardless of active-site engagement—a closed conformation is detected alongside the open form. For high-resolution models, the distance between the Cα atoms of Trp353 in the two BRE subunits (at the centre of the Ub_wrist_ binding interface) is indicated. Because closure of the BRCA1-A core involves a complex three-dimensional conformational change, the effective distance between the arm ubiquitin-binding sites and the active site can decrease by more than is suggested by the arm-to-arm distance. Active site state, EMDB map IDs and corresponding PDB IDs are indicated for each conformation (see also Supplementary EM information). For datasets where model building was not possible, the indicated model was fitted as a rigid body. “Apo” refers to a sample containing oligoUb_ATA_, with structurally intact but unoccupied active sites and Ub bound at the wrist sites. No closed conformations were observed in the diUb_ATA_ dataset. **(B) Structure and cartoon representation of a closed conformation in State C\***. Particles of BRCA1-A:oligoUb_ATA_ in closed conformation were 3D classified with a mask on BRCC36 and UbD. Of the four resulting subclasses, the one with the best-resolved coiled coil was used for model building. The sharpened density and fitted model of this class are shown. In this conformation, both active sites are occupied by UbD; one UbP adopts a conformation similar to State C (State C*), while the second UbP is disordered. Occupancy at the elbow and wrist ubiquitin binding sites is low. Map: EMD-55053; PDB: 9SO9 **(C) Superposition of Open and Closed state reveals complex transition**. Open BRCA1-A:polyUb_ATA_ (State C+P from Fig. 1D, white) and closed BRCA1-A:oligoUb_ATA_ (State C* from Fig. 5B, colour) structures—shown as backbone traces—were superposed on the BRE subunit indicated by the red dot. Ubiquitins are omitted for clarity. Arm closure brings each BRCC36 active site into closer proximity with the opposing ABRAXAS, bends the coiled coil, and pushes the BRE elbows inward. Dashed lines indicate regions of ABRAXAS that are disordered in the closed form. **(D) ABRAXAS controls BRCA1-A arm shape**. Backbone of one BRCA1-A pentamer is shown. The C-terminal RAP80-binding region and N-terminal MPN domain of ABRAXAS are rigidly connected via BRE. The schematic illustrates how movement of the ABRAXAS MPN, potentially triggered by ABRAXAS-finger engagement with BRCC36 of the opposite pentamer, can pull on the arm tip and bend the coiled coil. The forward and rear halves of the BRCC36 coiled coil bend independently due to two conserved prolines (yellow), acting as hinges. **(E) Closure of BRCA1-A shifts ubiquitin binding on the elbow region**. Particles corresponding to the closed form of BRCA1-A:polyUb_ATA_ (Fig. 5A, Supplementary Table 1) were 3D-classified with a spherical mask on the left arm (four classes). Density for a class with high chain occupancy on the arm is shown, with the model from Fig. 5B fitted. Ub_wrist_ and Ub_Elbow_ were modeled using the Alphafold3 model (ModelArchive ID: ma-yb0lm) as reference. Additional density consistent with ubiquitin appears above the elbow (Ub_supraelbow_), and Ub_elbow_ density shifts slightly forward (see close-up in Supplementary Fig. 5D). Ub_wrist_ remains unchanged. The right arm appears partially disordered due to variations in arm angle. UbD is present in both sites one UbP adopts a State C*–like conformation, while the second UbP is disordered. Map: EMD-55055 **(F) Arm closure pushes the engaged ubiquitin chain toward the opposing active site**. Open (State C+P from Fig. 1D, grey) and closed (State C* from Fig. 5E, colour) BRCA1-A structures bound to poluUb_ATA_ were superposed on the BRE subunit indicated by the red dot. Ubiquitins from the chain feeding into the State C / State C* site are shown as solids. Lys 63 of Ub_elbow_ is marked in yellow. Arm closure displaces the chain toward the active site. W: Ub_wrist_; E: Ub_elbow_; P: UbP; D: UbD. **(G) Closure brings Ub**^**elbow**^ **within one ubiquitin length of UbP**. Close-up of panel G, with BRCA1-A shown as a surface, Ub_elbow_ in cartoon (pink: open; magenta: closed), and Lys63 displayed as sticks. A bridging ubiquitin (Ub_bridge_, white) was manually placed with its Lys63 linked to the C-terminus of UbP. Closure brings Lys63 of Ub_elbow_ into proximity with the Ub_bridge_ C-terminus. To assess reachability, all known PDB structures of ubiquitin were aligned and used to model Ub_bridge_ and the possible reach of the C-terminus of UbP; three conformations with suitably positioned termini are shown.

### Closing of BRCA1-A: a chain feeding mechanism

The interactions of closed-form BRCA1-A with either oligoUb_ATA_ or polyUb_ATA_ are similar to each other, but we focus on polyUb_ATA_, as it shows higher chain occupancy along the arms. In the BRCA1-A:polyUb_ATA_ complex, we observe multiple ubiquitin interaction states. Focused 3D classification revealed distinct subclasses, each characterised by different combinations of three main features: firstly, some subclasses exhibit additional density in front of BABAM1 (Supplementary Fig. 5D). BABAM1 has previously been reported to bind ubiquitin^13,49^, and the observed density aligns with the binding mode predicted by AlphaFold3, in which the ubiquitin Ile44 patch contacts a shallow surface on BABAM1 centred on Cys169 (Supplementary Fig. 5D). We confirmed that BABAM1 binds ubiquitin with low affinity, and that this interaction is disrupted by mutating Ala154 and Cys169 to lysine (Supplementary Fig. 5E). However, a BRCA1-A complex carrying the BABAM1^A154K,C169K^ mutant shows wild-type affinity for ubiquitin and ubiquitin chains and no catalytic impairment (Supplementary Fig. 5F), leaving the functional relevance of this site unresolved.

Additionally, all subclasses display clear density for distal ubiquitin (UbD) in both active sites, whereas proximal ubiquitin (UbP) density is often missing, indicative of conformational heterogeneity. In a subset of classes, we identify a configuration—designated State C*—in which one active site binds diUB_ATA_ in a conformation similar to State C (Fig. 5B and 5E). Within the active site, though, ATA appears disordered and the resolution is insufficient to confirm cleavage competence. Arm closure shifts the BRE elbow region closer to the active site. This suggests that State C* may represent a transitional configuration between State P and State C, potentially driven by (or resulting in) arm closure.

Finally, closure slightly alters ubiquitin chain engagement along the arms: binding remains detectable at both the wrist and elbow sites, and in some subclasses, additional density appears just above the elbow. This could correspond to the disordered C-terminus of RAP80, or alternatively, to an additional ubiquitin moiety that becomes ordered or more stably bound upon arm closure (Ub_supra-elbow_, Fig. 5E).

Notably, in the closed form, the Ub^elbow^ is shifted slightly forward toward the active site relative to its position in the open state (Supplementary Fig. 5D). Although the density does not allow precise modelling, an AF3 model of BRCA1-A in complex with ten monoUb copies predicts a closed conformation with ubiquitin binding at the distal, elbow, wrist, and BABAM1 positions on each arm (Supplementary Fig. 5A). The Ub^elbow^ predicted by AF3 matches the forward-shifted density observed in the experimental map of the closed form more closely than Ub_elbow_ from the open structures (Supplementary Fig. 5G).

Together, closure of the arms and forward shifting of the elbow ubiquitin substantially reduce the spatial separation between the elbow ubiquitin and the active site (Fig. 5G, Supplementary Movie 3). In the open form, the distance from Lys63 of the elbow ubiquitin to the catalytic zinc is 64 Å, reducing to 54 Å in the closed form. Crucially, in the open conformation (State C), the distance between UbP and the elbow-bound ubiquitin is too great to be spanned by a single ubiquitin moiety, requiring at least two to bridge the gap. Upon arm closure, in State C*, this distance is reduced such that a single ubiquitin is sufficient to connect the two (Fig. 5G). Thus, arm closure appears to push the chain forward by one moiety, effectively feeding the next ubiquitin into the active site.

## Discussion

We developed a diUb-based probe for metallo-DUBs featuring a zinc-chelating thiol warhead, diUb_ATA_, which can be produced with all linkage types and used to probe specificity and mechanisms of action of metallo-DUBs (Fig. 1C). DiUb_ATA_ appears to effectively mimic the native isopeptide linkage (Supplementary Fig. 3A), allowing characterization of enzymes in a near-native state without resorting to inactive mutants, which often display disordered active sites. Using chain-specific E2-E3 and DUB combinations, diUb_ATA_ can also be enzymatically extended into any linkage-specific polyUb chains (except K27, for which no E2 is yet known), thus enabling mechanistic studies of long chain processing. Alternatively, di- or polyUb_ATA_ can be conjugated to specific substrates to enable investigation of the effect of chain conjugation and length on the regulation of metallo-DUB substrates. The inclusion of alternating cleavable and uncleavable bonds allows post-catalytic states to be captured. Together, the diUb and polyUb formats offer a generalisable toolset for dissecting the activity of JAMM-family DUBs across diverse linkage contexts.

Our structural data on K63-linked diUb_ATA_ and polyUb_ATA_ in complex with BRCA1-A support a BRCC36 cleavage mechanism that is broadly consistent with prior predictions based on the structure of AMSH-LP^13,37,42^. However, BRCC36 diverges from AMSH-LP in its use of insertion 2 (Ins-2). In AMSH-LP, Ins-2 provides the primary interaction surface for UbP and contributes directly to catalysis by forming a lid over the reaction chamber, with Phe407 interacting with UbP Gln2.

By contrast, our structures show that Ins-2 plays no direct role in BRCC36 catalysis. Instead, Ins-1 fulfils both substrate positioning and capping functions: Glu88 and Lys97 position UbP through interactions with UbP Gln2, while Arg89 forms a lid over the catalytic centre by interacting with UbP Glu64 (Fig.2C). Disruption of this lid by mutation of Arg89 to Ala results in a pronounced catalytic defect (Fig. 2D).

Consistent with this shift in function, the binding mode of UbP also differs between AMSH-LP and BRCC36. In BRCC36, UbP is rotated relative to AMSH-LP and binds directly to the JAMM domain via Met1, rather than primarily engaging Ins-2. This reorientation shifts UbP Gln2 away from the lid role, which is instead fulfilled by Glu64 (Fig. 2C).

The lack of a catalytic role for Ins-2 is further supported by our biochemical data: both long and short Ins-2 isoforms of BRCC36 are catalytically active (Fig. 3D,E). Why mammals retain two distinct splicing isoforms of BRCC36 remains unclear, it could depend on some yet unknown interactor or PTM. Alternatively it could be primarily relevant for the assembly or regulation of the paralogous BRISC complex^53^.

A novel feature of BRCA1-A catalysis is the role of the ABRAXAS-finger, a loop that extends from ABRAXAS’ to Ins-1 of BRCC36 in the opposite pentamer. Upon substrate binding, it interacts with the lid arginine (Arg89) and engages the C-terminus of UbD (Fig. 2A and Supplementary Fig. 3D). Mutation of the tip of the loop results in a severe loss of activity (Fig. 2E), highlighting its essential role in catalysis. The arrangement of this loop is conserved in both ABRAXAS and ABRO1, offering an opportunity to develop BRCA1-A- and BRISC-specific catalytic mutants—valuable tools for dissecting the distinct cellular roles of each complex.

BRCA1-A is a K63-specific DUB with a clear preference for polyubiquitin over diUb. This preference arises from a combination of avidity and processivity. In addition to the diUb binding in the active site, BRCA1-A has two low-affinity ubiquitin binding sites on each arm (Fig. 4D and 4E), and all of these together confer high-affinity binding—but only when the chain is long enough to span them (Fig. 4D). Thus, long chains are recognised more effectively (Fig. 4F).

BRCA1-A can engage two chains simultaneously (one per active site) but cleavage occurs in alternation (State C+P, Fig. 2A, Supplementary Movie 2). After cleavage at the State C site, UbP must dissociate from BRCC36 to allow the opposing site to transition from state P to state C. If the cleaved chain remains bound to the arm binding sites after cleavage, BRCA1-A can close its arms, and feed the chain forward for subsequent cleavage, enabling processivity. This mechanism applies only to chains that fully span the arm binding sites prior to cleavage (i.e. Ub_6+_, Fig. 4D and 5F). Accordingly, processivity, together with avidity, is expected to accelerate cleavage of long chains (Ub_7+_) whereas intermediate (Ub_4–6_) and short (Ub_2–3_) chains are processed more slowly and largely in a distributive manner (Fig. 2B). Functionally, this positions BRCA1-A as a chain-shortening DUB, acting on shorter chains only after longer ones have been trimmed.

*In vivo*, BRCA1-A’s recruitment at DSB sites depends on K63-linked polyubiquitin chains deposited in an RNF8/Ubc13-dependent manner^7,18,26^. At the same time, BRCA1-A activity has been shown to oppose RNF8/Ubc13-dependent ubiquitination at sites of DNA double-strand breaks^27^. This paradox—recruitment by K63 chains and suppression of their accumulation— may be explained by BRCA1-A’s intrinsic propensity to act as a chain trimmer rather than eraser. Given that RAP80’s tandem UIM domains can bind as little as diUb^23^, short K63-linked chains may be sufficient for BRCA1-A recruitment, while longer chains would be selectively targeted for degradation.

In DNA damage tolerance, choice of the pathway used to bypass DNA is influenced by the length and type of Ub chains conjugated to PCNA (reviewed in^54^), with multiple E3s and DUBs dynamically determining the balance^55–60^. How K63-polyUb chain length influences pathway choice in DSB repair is currently unknown. DUB-inactive mutants of BRCC36 lead to accumulation of K63-Ub chains^27^, increased DNA resection and hyperactivation of HR repair^32^, outlining a possible connection for future investigation. Extended K63-linked polyubiquitin chains are associated with phase separation behaviours, particularly when interacting with proteins containing multiple ubiquitin-binding domains (reviewed in ref ^51^). BRCA1-A might therefore also influence the local physical properties of DNA damage sites— not only by modulating ubiquitin signalling, but also by regulating the solubility and diffusion behaviour of associated repair factors.

### Arm dynamics coordinate substrate feeding and cleavage

The structural states we described suggest a working model for the catalytic mechanism of BRCA1-A (Fig. 6A, Supplementary Movie 5). In presence of abundant substrate, both active sites can engage chains, quickly reaching a configuration with, e.g. site 1 in State C and site 2 in State P (Fig. 6A, intermediate 1, corresponding to our open structure). Cleavage at site 1 leads to release of UbD and likely dissociation of UbP from the active site, though the chain remains anchored to the arm through the avidity provided by the combined elbow and wrist binding sites (Fig. 6A, intermediate 2). At site 2, UbP’ needs to move forward in order to reach state C. If the chain has slack, i.e. more than the minimal number of one ubiquitin connects UbP’ to Ub’_elbow_, UbP’ can slot into the active site and be cleaved. However, if the chain is taut, UbP’ cannot yet enter the active site, as the chain is too short to span the required distance and is being held back by the arm. Arm closure resolves this limitation by moving UbP’ closer, effectively feeding the chain into the active site (Fig. 6A, intermediate 3, our polyUb_ATA_ closed structure Fig. 5F): site 2 transitions into State C*, with UbP’ now positioned in the active site. At site 1 the previously cleaved chain, still bound to the arm, is brought closer to BRCC36 and can presumably re-engage the active site. With the ends of both chains now held by the active sites, the arms must release their grip to swing back to the open configuration, and then re-engage the chains further along (Fig. 6A, intermediate 4), thereby restoring a State C + State P arrangement with the roles of the two sites reversed, and one chains advanced by (at least) one moiety. Chain release by the arm upon opening likely depends on reduced chain-binding affinity in the closed conformation, consistent with our observation that the elbow and wrist sites are more highly occupied in the open state than in the closed state. Future studies, including single molecule FRET analysis, and/or trapping of additional intermediates are needed to resolve the details of this mechanism.

**Figure 6:**
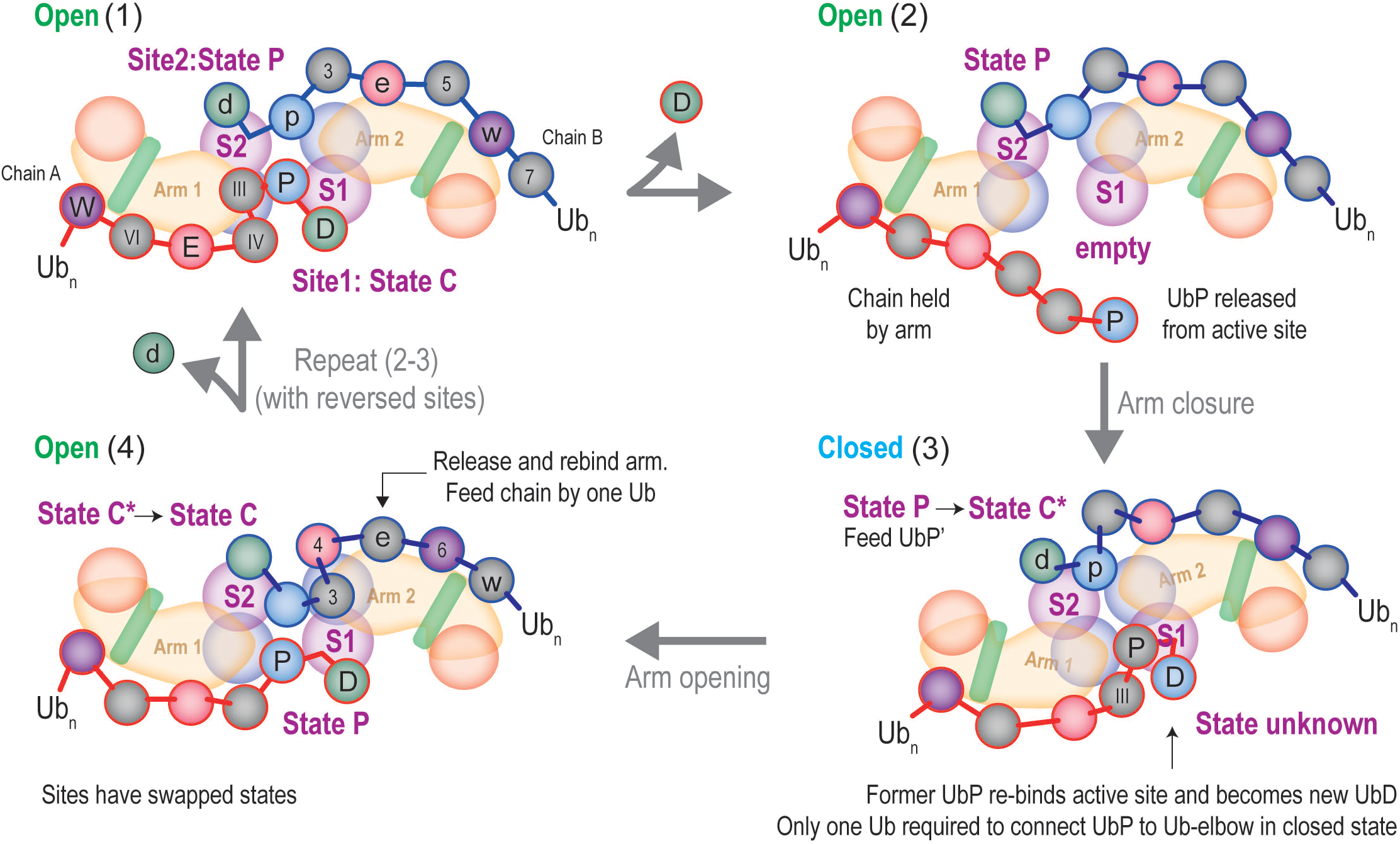
A model for chain feeding in BRCA1-A. Proposed mechanism for processive polyUb cleavage by BRCA1-A. (1) (1) Two polyUb chains are bound without slack: for example, active site 1 is in State C and active site 2 in State P. Ubiquitin moieties are represented as circles, with their positions indicated as *w* (Ub_wrist_), *e* (Ub_elbow_), *p* (UbP), *d* (UbD), or numbered by position relative to UbD. Chains are shown with no slack. Moieties from chain A (bound to arm 1 and feeding into active site 2) are labelled with uppercase letters and Roman numerals; moieties from chain B (bound to arm 2 and feeding into active site 1) are labelled with lowercase letters and Arabic numerals. (2) UbD is cleaved from chain A at site 2, releasing the chain from the active site but not from the arm due to avidity. (3) Arm closure enables UbP from chain B (marked p) to transition from State P to State C*, entering active site 1. Meanwhile, chain A re-engages active site 2, with its former UbP and Ub III moieties now serving as UbD and UbP, respectively (4) As the arms reopen, chain A and site 2 return to State P. For active site 1 to re-enter State C, chain B— which is now one Ub too short—must dissociate from arm 2 and rebind further downstream. The complex thus returns to the initial configuration, but with the states of the active sites reversed. Repeated cycles of (2)–(4) would return the complex to step (1), enabling processive cleavage. Arm alternation is compatible with this model. Slack in one or both chains would permit multiple cleavage cycles without requiring arm closure.

Overall, this model outlines a potential mechanism by which BRCA1-A combines selective avidity and processivity to achieve its substrate preference for long over short polyubiquitin chains.

## Materials and Methods

### Chemical synthesis

#### General Fmoc SPPS Strategy

Solid phase peptide synthesis (SPPS) is based on El Oualid et al.^44^ and Mulder et al.^61^ In brief, SPPS is performed in N-methylpyrrolidone (NMP) on a Syro II MultiSyntech Automated Peptide synthesizer using Fmoc-Gly-trityl resin (0.18 mmol/g, RA1213, Rapp Polymere GmbH), standard 9-fluorenylmethoxycarbonyl (Fmoc) based solid phase peptide chemistry (25 – 50 μmol scale) and a 4-fold excess of amino acids relative to resin. The following amino acids were used: Fmoc-L-Ala-OH, Fmoc-L-Arg-(Pbf)-OH, Fmoc-L-Asn(Trt)-OH, Fmoc-L-Asp(OtBu)-OH, Fmoc-L-Gln(Trt)-OH, Fmoc-L-Glu(OtBu)-OH, Fmoc-L-Gly-OH, Fmoc-L-His(Boc)-OH, FmocL-Ile-OH, Fmoc-L-Leu-OH, Fmoc-L-Lys(Boc)-OH, Fmoc-L-Met-OH; Fmoc-L-Phe-OH; Fmoc-L-Pro-OH; Fmoc-L-Ser(tBu)-OH; Fmoc-L-Thr(tBu)-OH, Fmoc-L-Tyr(tBu)-OH, Fmoc-L-Val-OH. The following pseudoproline and DMB dipeptide building blocks were used at the indicated dipeptide sequences: Fmoc-L-Leu-L-Thr(Ψ^Me,Me^pro)-OH at position 8-9, FmocL-Ile-L-Thr(Ψ^Me,Me^pro)-OH at position 13-14, Fmoc-L-Ala-(Dmb)Gly-OH at position 46-47, Fmoc-L-Asp(OtBu)-(Dmb)Gly-OH at position 52-53, Fmoc-L-Leu-L-Ser(Ψ^Me,Me^pro)-OH at position 56-57 and Fmoc-L-Ser(tBu)-L-Thr(Ψ^Me,Me^pro)-OH at position 65-66. Cycles 1–30 were performed using single couplings in NMP for 45 min with PyBOP (benzotriazol-1-yloxytripyrrolidinophosphonium hexafluorophosphate, 4 equiv) and DiPEA (N,N-diisopropylethylamine, 8 equiv). Fmoc deprotection was carried out using 20% piperidine in NMP (2 × 2 min and 1 × 5 min). Dipeptides were coupled for 2 h. After cycle 30, the coupling time was extended to 60 min, and Fmoc deprotection was performed using 20% piperidine in NMP (4 × 3 min). For residues P37 and T12, incomplete incorporation following single coupling was observed; this was mitigated by performing double couplings (2 × 90 min). Dipeptides were coupled for 2 h throughout. Double couplings were additionally applied for cycles 62–69 (35 min each).

#### LC-MS analysis

LC-MS measurements were performed on a system equipped with a Waters 2795 Seperation Module (Alliance HT), Waters XSelect CSH C18 (4.6 x 100mm, 5 μm) and Waters LCT Premier XE Mass Spectrometer. Samples were run (column temperature= 40°C, flow= 0.8 mL/min) using 2 mobile phases: A= 1% CH_3_CN and 0.1% formic acid in water and B= 1% water and 0.1% formic acid in CH_3_CN. Data processing was performed using Waters MassLynx Mass Spectrometry Software 4.1.

#### RP-HPLC purification

RP-HPLC was performed on a system equipped with a Waters 600 HPLC and controller, Waters XSelect CSH C18 (30 x 250mm, 5 μm) column and Waters Dual Absorbance Detector 2487 detector. Samples were run (flow= 25 mL/min) using 2 mobile phases: A= 5% CH_3_CN and 0.05% trifluoroacetic acid in water and B= 5% water and 0.05% trifluoroacetic acid in CH_3_CN. Data analysis was performed using Waters MassLynx Mass Spectrometry Software 4.1.

#### Ub(1-75)-MMP

All compounds referred to in the following procedures are numbered as shown in Supplementary Fig. 1. Fully side-chain protected Ub(1-75) *(1)* was synthesized on a 50 umol scale on Fmoc-Gly-trityl resin following the general procedure described previously^44,61^. The N-terminus was protected with a tert-butyloxycarbonyl (Boc) group by treatment with 5 equiv di-tert-butyl dicarbonate and 10 equiv DiPEA in 10 mL CH_2_Cl_2_ for 45 min. The resin was washed with CH_2_Cl_2_, treated with 20 mL CH_2_Cl_2_/hexafluoro-2-propanol (4:1 v/v) for 45 min and filtered. The resin was rinsed with 1,2-dichloroethane (10 mL) and the combined filtrates were concentrated. Residual hexafluoro-2-propanol was removed by co-evaporation with 1,2-dichloroethane (3×25 mL). The fully protected protected Boc-Ub(1-75) (*2*, 1 equiv, 50 µmol) was dissolved in CH_2_Cl_2_ (20 mL) and reacted overnight with PyBOP (5 equiv), methyl 3-mercaptopropionate (MMP, 5 equiv) and DiPEA (10 equiv). The solvent was removed in vacuo and the residue treated for 3 hrs with 10 mL trifluoroacetic acid/H_2_O/iPr_3_SiH (92/5/3 v/v/v), followed by precipitation in 90 mL cold ether. The product was pelleted (5 min at 4000 rpm) and washed with ether (3 times). The pellet was next dissolved in 5 mL warm dimethylsulfoxide and added slowly while stirring to 100 mL HPLC buffer A. RP-HPLC purification (gradient: 20 – 30%B over 20 min) and lyophilization of the LC-MS pooled HPLC fractions afforded Ub(1-75)-(methyl 3-mercaptopropionate) thioester (*Ub(1-75)-MMP*) as a white powder. This was lyophilized and additional 2 times in 25 mL CH_3_CN/H_2_O/formic acid (50/50/1, v/v/v). Yield: 172 mg, 20 μmol, 40%.

#### Ub K63Dab(ATAStBu) mutant

Fully side-chain protected Ub(1-75) (50 µmol) with a K63-to-Dab(Alloc) mutation and free N-terminus (*3*, Fig. 1S) was synthesized on Fmoc-Gly-trityl following the general procedure described previously^44,61^. The N-terminus was protected with a Boc group by treatment with 5 equiv di-tert-butyl dicarbonate and 10 equiv DiPEA in 10 mL CH_2_Cl_2_ for 45 min. After washing the resin with NMP and CH_2_Cl_2_, the allyloxycarbonyl (Alloc) group was removed by treating the resin with Pd(PPh_3_)_4_ (0.5 equiv) and Ph_3_SiH (20 equiv) in CH_2_Cl_2_ (2×20 min), affording resin *4*. After washing with NMP and CH_2_Cl_2_, the resin was treated for 1 hr with 2 equiv 4-((tert-butoxycarbonyl)amino)-3-(tert-butyldisulfanyl)butanoic acid (*5*, Fig. 1S) (Mulder et al.)^61^, PyBOP (5 equiv) and DiPEA (10 equiv). After washing with NMP and CH_2_Cl_2_, the resin was treated with 20 mL CH_2_Cl_2_/hexafluoro-2-propanol (4:1 v/v) for 45 min and filtered. The resin was rinsed with 1,2-dichloroethane (10 mL) and the combined filtrates were concentrated. Residual hexafluoro-2-propanol was removed by co-evaporation with 1,2-dichloroethane (3×25 mL). The fully protected protected *6* (1 equiv, 50 µmol) was dissolved in CH_2_Cl_2_ (20 mL) and reacted overnight with PyBOP (5 equiv), H-Gly-OtBu (HCl salt, 5 equiv) and DiPEA (15 equiv). The solvent was removed in vacuo and the residue treated for 3 hrs with 10 mL trifluoroacetic acid/H_2_O/iPr_3_SiH (92/5/3 v/v/v), followed by precipitation in 90 mL cold ether. The product was pelleted (5 min at 4000 rpm) and washed with ether (repeat 3 times). The pellet was dissolved in 5 mL warm dimethyl sulfoxide and added slowly while stirring to 100 mL HPLC buffer A. RP-HPLC purification (gradient: 20 – 30%B over 20 min) and lyophilization of the LC-MS pooled HPLC fractions afforded *7* as a white powder. Lyophilization is repeated 2 times in 25 mL CH_3_CN/H_2_O/formic acid (50/50/1, v/v/v). Yield: 100 mg, 11.4 μmol, 22%.

#### diUb_ATA_ probe (K63-linked diUb K63Dab(ATA)

Ub mutant *7* (40 mg, 4.6 μmol) and **Ub(1-75)-MMP** thioester (54 mg, 6.3 μmol) were dissolved in 2 mL 6M Gdn·HCl, 0.15 M sodium phosphate, 200 mM 4-mercaptophenylacetic acid (MPAA), pH 8. Centrifugation at 4000 rpm was used to aid dissolution if the reaction proceeded slowly. Next, analysis with a micro pH probe showed the pH had dropped to 6.3, and this was carefully adjusted to 7.0 with 1 N NaOH. The reaction mixture was incubated overnight at 37ºC after which LC-MS analysis showed complete consumption of *7* and formation of product. The reaction mixture was diluted with 38 mL 6M Gdn·HCl, 0.15 M sodium phosphate, 25 mM tris(2-carboxyethyl)phosphine pH 7, mixed for 30 min and purified by RP-HPLC (gradient: 18 – 25%B over 30 min). After lyophilization of the LC-MS and SDS-PAGE pooled HPLC fractions, the **diUb**_ATA_ probe was lyophilized and additional 2 times in 25 mL CH_3_CN/H_2_O/formic acid (50/50/1, v/v/v). Yield: 28 mg, 1.6 μmol, 36%, white powder.

### DNA sequences

DNA constructs encoding human ABRAXAS (Gene name: ABRAXAS1, Uniprot id: Q6UWZ7), BRCC36 (BRCC3, Uniprot id: P46736), BABAM1 (Uniprot id: Q9NWV8), BRE (BABAM2, Uniprot id: Q9NXR7) were synthesized as codon-optimised sequences (see Synthetic DNA sequences). Human RAP80 (UIMC1, Uniprot id: Q96RL1) was amplified from human cDNA, with final DNA/protein sequence corresponding to Genbank AF349313.1. This natural variant carries a Glu347->Gly mutation compared to the canonical sequence in Uniprot Q96RL1.

### Expression of GST-BRE in *Escherichia coli*

Full length, *wild type* BABAM2 was subcloned into vector pGEXNKI-his-GST-3C-LIC^62^ (Addgene id: 108709) using Ligation Independent Cloning (LIC). Primers were designed using CCD2^63^. BABAM2 mutants in “wrist” position (350-SPRW-353 -> AAAA), “elbow” position (R137D F140A), or the combined “wrist” + “elbow” mutant were generated by site directed mutagenesis using the *wild type* construct as template.

For expression, the appropriate constructs were transformed in *E. coli* BL21 (DE3) Rosetta PlysS. Selection was carried out on with ampicillin (50 µg/mL) and chloramphenicol (37 µg/mL) at all stages. A single colony was picked from a fresh agar plate and used to start an overnight preculture in Luria Bertani (LB) broth. The preculture was then used at 1:500 ratio to inoculate a large-scale culture (typically 2L, with 500 mL Terrific Broth in each 3.2-liter, baffled Fernbach flask). The culture was grown at 37°C until OD 1.5 was reached, then shifted to 20°C. Upon reaching that temperature, expression was induced with 0.25 mM Isopropyl β-d-1-thiogalactopyranoside (IPTG) and carried out overnight at 20°C.

After harvesting, the cells were resuspended in lysis buffer (300 mM NaCl, 20 mM HEPES, pH 7.8) with 0.5 mM 4-(2-aminoethyl)benzenesulfonyl fluoride (AEBSF) as protease inhibitor and 2 mM dithiothreitol (DTT). Lysis was carried out by sonication, and the lysate was clarified by centrifugation at 49000 RCF for 45’. The clarified lysate was applied with a peristaltic pump (1 mL/min) to 5 mL of Glutathione Sepharose High Performance resin (Cytiva Life Sciences) packed in a XK-16/20 column (Cytiva). Contaminants were washed with 8 column volumes of 1M NaCl, 20 mM HEPES pH 7.8, followed by re-equilibration to gel filtration buffer (150 mM NaCl, 20 mM HEPES pH 7.8). Elution was carried out with gel filtration buffer supplemented with 20 mM reduced glutathione. The eluate was pooled, supplemented with 2 mM DTT and 0.5 mM AEBSF, and concentrated by ultrafiltration with Amicon Ultra Centrifugal filter units (30 kDa molecular weight cutoff). GST-BRE was then separated from the remaining contaminants (free GST and an unknown band of about 65 kDa) by size exclusion using a Superdex 75 16/60 column (Cytiva) and gel filtration buffer. Note that owing to its intrinsic dimerization properties, free GST always co-purified with GST-BRE. GST-BRE was then concentrated to about 5 mg/mL, supplemented with 5 mM (tris(2-carboxyethyl)phosphine (TCEP), flash frozen in liquid nitrogen and stored at −80°C in small aliquots. The identity of the purified protein was confirmed by western blotting (Anti BRE, sc-376453, Santa Cruz Biotechnology). Point mutants were purified with the same protocol and behave similarly during purification.

### Expression of BRCA1-A complex in insect cells

#### Generation of Constructs

In order to achieve co-expression, the genes corresponding to the five members of the human BRCA1-A complex were subcloned into three separate DNA constructs: a bicistronic pFastDual (Invitrogen) vector carrying ABRAXAS (gene symbol: ABRAXAS1) and BRCC36 (BRCC3); a pFastDual vector carrying BRE (BABAM2) and 6xHis-tagged-BABAM1; and a pFastBac-derived, monocistronic vector carrying StrepII-tagged-RAP80 260-413 (UIMC 260-413). The vectors were constructed as follows:

#### pFASTDual:ABRAXAS:BRCC36 and mutants

Synthetic sequences (see Synthetic DNA sequences) coding for ABRAXAS and BRCC36 were amplified by PCR using oligos carrying restriction sites for SalI (5’) and SpeI (3’), then inserted by standard restriction cloning into, respectively, the pH cassette and p10 cassette of the pFASTDual vector (Invitrogen). For this operation, the SalI/SpeI of the pH cassette and the NheI/XhoI sites of the p10 cassette were used. This strategy exploited the fact that SalI and SpeI form compatible ends with NheI and XhoI respectively, and that the p10 cassette is on the antisense strand. Point mutations in ABRAXAS were engineered by site-directed mutagenesis of the pFASTDual:ABRAXAS:BRCC36 vector.

#### pFASTDual:BRE:His-3C-BABAM1 and mutants

A synthetic sequence coding for full-length BABAM1 was cloned into pETNKI-his-3C-LIC-kan (NKI-1.1^62^ Addgene # 108703) for *E. coli* expression, yielding a construct that carries a 3C-cleavable, N-terminal, hexahistidine tag. This construct, and a synthetic sequence encoding BRE were then cloned, respectively, into the p10 and pH cassette of pFastDual using the same strategy as BRCC36/ABRAXAS above. Point mutations in BRE were substituted into the pH cassette by restriction cloning from the *E. coli* expression vectors.

#### NKI2.9:StrpII-3C-RAP80^260-413^

Residues 260-413 of RAP80 were cloned into a pFastBac-derived ligation independent cloning vector bearing a 3C-cleavable, N-terminal StrepII^64^ (NKI 2.9^62^, NKI protein facility).

#### Baculovirus generation

Baculoviruses for all three constructs were generated using the Bac-to-Bac protocol (Invitrogen), with either *E. coli* DH10Bac (Invitrogen) or *E*.*coli* EMBacY^65^ as transposition strain, and *Spodoptera Frugiperda* SF9 cells for virus production. The choice of transposition strain did not have an effect on protein expression or protein quality; however, ABRAXAS:BRCC36 viruses generated using EMBacY-derived bacmids had much shorter shelf life than those generated by DH10Bac-derived bacmids, for reasons unknown.

#### BRCA1-A Protein expression

The BRCA1-A complex was expressed by simultaneously infecting SF9 cells (1 million cells/mL in ESF 921 medium, Expression Systems) with pFASTDual:ABRAXAS:BRCC36, pFASTDUal:BRE:His-3C-BABAM1 and NKI2.9:StrpII-3C-RAP80^260-413^, either *wild type* or mutants where appropriate. The relative amounts of each virus required for optimal expression were batch dependent and were determined by small scale expression tests. After infection, cells were grown until viability dropped below 90% (approximately 60 hours). The cells were then harvested by centrifugation, resuspended in lysis buffer supplemented with EDTA free protease inhibitor cocktail (Pierce) and either used for protein purification or flash frozen in liquid nitrogen and stored at −20°C.

#### BRCA1-A protein purification

Insect cells expressing BRCA1-A were lysed by sonication, and the lysate was clarified by centrifugation (49000 RCF, 45 min). The clarified lysate was applied to 10 mL Talon resin (Takara Bio) in a gravity column. The resin was washed with two column volumes of 1M NaCl, 20 mM imidazole, 20 mM HEPES pH 8.0, followed by re-equilibration with 150 mM NaCl, 20 mM HEPES, 20 mM imidazole pH 8.0, and elution with 150 mM NaCl, 20 mM HEPES, 200 mM imidazole, pH 8.0 (five column volumes). The eluate was supplemented with DTT to a final concentration of 2 mM and immediately loaded onto a 10 mL Streptactin XT 4flow (IBA LifeSciences) column (1ml/min). The column was washed with gel filtration buffer (150 mM NaCl, 20 mM HEPES pH 7.8) before elution with gel filtration buffer plus 50 mM biotin (final pH 7.8). The eluate was concentrated with Amicon Ultra Centrifugal filter units (100 KDa cutoff) and buffer-exchanged by size exclusion chromatography (Superose 6 or Superose 6 Increase 10/300, Cytiva). The complex was finally concentrated to 1-10 mg/mL, supplemented with 5 mM TCEP, flash frozen and stored at −80°C. In preliminary experiments, the BRCA1-A variant bearing the BRCC36 R89A mutation precipitated in the high-imidazole elution buffer following the Talon affinity step. For this variant, purification was therefore performed using only the Strep affinity step, including a two-column-volume wash with 1 M NaCl prior to elution. This modification had minimal impact on sample purity.

Sample quality was assessed by size-exclusion chromatography, loading 30–100 μg of purified sample onto a Superose 6 Increase 3.2/300 column (Cytiva) equilibrated in gel filtration buffer. Chromatograms are shown in Supplementary Figure 6B.

#### Thermal Stability Assays

*Wild type* and mutant BRCA1-A complexes were diluted to 1-5 μM in 150 mM NaCl, 20 mM HEPES pH 7.8, 5 mM TCEP, and assayed for thermal stability using a Prometheus NT.48 (NanoTemper Devices). Protein complexes were denatured using a thermal ramp from 20 to 95 °C at 1 °C/min, and unfolding and aggregation were monitored by intrinsic tryptophan fluorescence at 330 and 350 nm, expressed as the fluorescence ratio (350/330 nm). Melting temperatures were calculated using Prometheus software with default parameters. Melting curves are shown in Supplementary Figure 6. Calculated melting temperatures (mean ± standard deviation from at least two independent measurements): *wild type*: 65.6 ± 0.9 °C; BRCA1-A^iso2^: 65.1 ± 1.4 °C; BRCA1-A^BRCC36-R89A:^ 66.0 ± 0.7 °C; BRCA1-A^ABX3A^: 64.0 ± 1.7 °C; BRCA1-A ^BABAM1 A154K C169K^: 65.7 ± 0.1 °C; BRCA1-A^BRE W165A^: 67.7 ± 0.7 °C; BRCA1-A^mut(wrist)^ 65.6 ± 1.4 °C; BRCA1-A^mut(elbow+wrist)^: 62.8 ± 0.1 °C.

### Ubiquitin chain digestion

#### Preparation of human ubiquitin

A construct bearing untagged, wild type human ubiquitin (pET3A-Ubiquitin, gift of Prof. Cecile Pickart) was transformed in *E. coli* E. coli BL21 (DE3) and used to inoculate 2L of Terrific Broth (Ampicillin 50 mg/mL). Cells were grown to OD 1.5, then induced with 0.5 mM IPTG and grown overnight at 25° C. Cells were harvested in 300 mM NaCl, 20 mM HEPES pH 7.5 and lysed by sonication. The lysate was clarified by centrifugation (49000 RCF, 30’) and 70% perchloric acid was added to the supernatant while stirring (final concentration 5%). Precipitate was removed by centrifugation (49000 RCF, 30’), and the supernatant was diluted 1:10 vol/vol with 50 mM ammonium acetate, pH 5.2. The sample was then loaded on a 3 mL PorosS cation exchange resin (Thermo Fischer) and eluted with a linear gradient of 12 column volumes (buffer A: 50 mM ammonium acetate pH 5.2; buffer B: 1M NaCl, 50 mM ammonium acetate pH 5.2). The peak corresponding to ubiquitin was collected, concentrated with Amicon Ultra Centrifugal filter units (3 kDa cutoff) and buffer exchanged into 150 mM NaCl, 20 mM HEPES pH 7.5 using a Superdex 75 16/60 gel filtration column (Cytiva). Ubiquitin was then concentrated by ultrafiltration to 1-10 mM and flash frozen in liquid nitrogen for storage at −80 °C. Concentration was measured by absorbance at 280 nM assuming Abs_280_(1 g/l) = 0.174.

#### Preparation of K63-Ub chains

0.5 µM each of His-Uba1 (E1), GST-Ubc13:MMS2 (E2) and GST-RNF8^351-485^ (E3) (preparation as in reference 15), and 0.5 mM human ubiquitin were incubated overnight (long chains) or 25’ (short chains) at 37°C in 150 mM NaCl, 20 mM HEPES pH 7.8, 5 mM ATP, 5 mM MgCl_2_, 2 mM DTT. After incubation, polymerization was stopped by addition of 20 mM iodoacetamide, and the mixture was dialyzed overnight against 50 mM ammonium acetate (pH 4.5), resulting in precipitation of E1, E2, and E3 enzymes. The chains were then concentrated by ultrafiltration (30 or 10 kDa molecular weight cutoff) and separated by in to pools with different lengths of chains using gel filtration (150 mM NaCl, 20 mM HEPES, pH 7.8; short chains: Superdex 75 10/300; long chains: Superose 6 10/300). Gel filtration fractions were flash-frozen and stored at −80 °C.

#### Preparation of K63-Ub probe chains

Probe chains were prepared using the same reaction conditions as long *wild type* chains, with 1µM diUb_ATA_ instead of Ub. Two preparations of different length (oligoUb_ATA_, shorter and polyUb_ATA_, longer; Supplementary Figure 4D) were used for structure determination. No size separation was performed.

#### Digestion of K63-Ub chains and K63-Ub2

BRCA1-A variants were prepared at 1 μM, then diluted in reaction buffer (150 mM NaCl, 20 mM HEPES, pH 7.5, 2 mM TCEP) to generate a 20× stock. The final enzyme concentration was 5 nM in all experiments except Fig. 3E (1 nM). K63-Ub chains were diluted in the same buffer. Both stocks were aliquoted into PCR tubes and prewarmed to 37 °C. Reactions were initiated by adding enzyme to the chain mixture and carried out at 37 °C in a PCR machine. At specified time points, 10 μL aliquots were removed and mixed with 5× SDS–PAGE buffer. The reaction proceeded too rapidly to obtain a true zero time point; instead, 9.5 μL of the K63-Ub chain mix was used. As a loading control, 5 μL of the 1 μM enzyme stock was included. All sample handling, except preparation of the 1 μM stock, was performed using a multichannel pipette. The samples were then resolved by SDS-PAGE using Bolt Bis-Tris Plus Mini Protein Gels in MES-SDS buffer (Thermo Scientific) and stained by Coomassie blue.

#### Quantification of Ub species from digestion experiments

Gel band intensities were quantified using a custom image analysis workflow (see Supplementary Code 1). A global background value was subtracted from all images prior to analysis. Gel lanes were manually defined and extracted as fixed-width (38-pixel) vertical stripes. For each lane, intensity profiles were generated by averaging pixel values across each row and baseline-corrected using asymmetric least squares (λ = 10^7^; p = 0.01). Peaks in the lane intensity profiles (corresponding to gel bands) were manually identified and numerically integrated to obtain band intensities. All experiments were repeated at least three times with comparable results.

#### Half Life Estimation of Ub species

Half-lives of Ub species were estimated by fitting a one-phase exponential decay with plateau model in GraphPad Prism [Plateau + (Y_0_−Plateau) * exp (−K(X−X0))] to the decay phase of each ubiquitin species, where X_0_ corresponds to the time point of maximal intensity. The decay phase was defined as all time points from the intensity maximum onwards. Fits were performed by nonlinear least squares, and half-lives (t_1_/_2_) are reported with 95% confidence intervals. Statistical significance of differences in half-lives between different Ub species was estimated using the Akaike information criterion in GraphPad Prism.

### BRCA1-A interaction with diUb_ATA_

#### Size exclusion chromatography

BRCA1-A complex (200 μg, 10 μM) was incubated with 100 μM diUb_ATA_ at 4 °C for 5 min in 150 mM NaCl, 50 mM HEPES pH 8.0, and 2 mM TCEP (total volume: 50 μL). The mixture was separated by size-exclusion chromatography using a Superose 6 Increase 3.2/300 column on an ÄKTA micro FPLC (originally manufactured by Amersham plc, now part of Cytiva), resolved by SDS–PAGE, and stained with Coomassie blue. Equal amounts of BRCA1-A or diUb_ATA_ alone were run as controls. The identity of the shifted diUb_ATA_ bands was confirmed by anti-ubiquitin western blot (Anti Ub antibody: sc-8017 HRP, Santa Cruz Biotechnology).

#### diUb_ATA_ inhibition

BRCA1-A (500 nM) was incubated with varying molar ratios of diUb_ATA_ for 10 min at 37 °C, then added as a 10× stock to K63 diUb (UbiQ BV; final concentration 10 μM) in 150 mM NaCl, 50 mM HEPES pH 8.0, and 2 mM TCEP. Digestion was performed at 37 °C, and aliquots were collected at the indicated time points. Reaction products were analysed by anti-ubiquitin western blot.

### CryoEM

#### Grid preparation

DiUb_ATA_: 5 µM BRCA1-A (1.9 mg/mL, 150 mM NaCl, 20 mM HEPES pH 8.0, 2 mM TCEP) was mixed with 25 µM diUb_ATA_ probe (same buffer, but 50 mM HEPES pH 8.0) and incubated 20’ at 37°C. Then 10x detergent buffer (0.4% Octyl β-D-glucopyranoside, 50 mM HEPES pH 7.5, 150 mM NaCl) was added to the sample before freezing. Grids were frozen with a Vitrobot (Thermo Fisher Scientific) as follows: 3 ul of sample were spotted onto Quantifoil 2/2, 400-mesh copper grids (glow discharge: 30 mA, 1 minute, at 0.2 mbar pressure). Excess sample was blotted with blot force 0 for 4.5 seconds and plunge frozen in liquid ethane. Five non-overlapping micrographs were collected per hole, shot close to the rim to account for particle lensing in the center of the hole (see Supplementary Table 1 and EM processing supplement for data collection statistics).

Apo: 1.8 μL of 10 μM BRCA1-A were incubated with 10 μL of oligoUb_ATA_ (Supplementary Fig. 4F) at 37 °C for 30 min. The mixture was then diluted with 8.2 μL of 150 mM NaCl, 50 mM HEPES pH 8.0, and 2 mM TCEP (no detergent) and used to freeze grids under the same blot and timing conditions as for diUb_ATA_. Despite the presence of oligoUb_ATA_—detectable as Ub_wrist_ moieties in the reconstructed density—the active sites appeared empty. The density in the active sites indicates that they are structurally intact and aligned, including density for the catalytic water. We speculate that an expired batch of TCEP may have allowed oxidation of the oligoUb_ATA_ thiol during processing, resulting in steric incompatibility with the active site.

OligoUb_ATA_: Prepared as for Apo, but using fresh TCEP stock.

PolyUb_ATA_: Prepared as for diUb_ATA_, but using polyUb_ATA_ (Supplementary Fig. 4F).

#### Data collection and processing

The structures presented in this study were derived from multiple datasets collected at the Netherlands Center of Nanoscopy (NeCen) on a 300 kV FEI Titan Krios microscope equipped with a K3 detector (Gatan) and an energy filter with a slit width of 20 eV. (Gatan). All datasets were processed using a consistent workflow (see EM processing supplement). Data processing was performed in cryoSPARC (v4+, Structura Biotechnology Inc.). Raw movies were motion-and CTF-corrected using default parameters, followed by manual curation to discard aberrant micrographs. Particle picking was performed using three approaches: (1) blob picking, (2) the general model in Topaz⍰(Topaz v0.2.6)^66^, and (3) the general model in crYOLO (v1.8.2)^67^. Resulting particle sets were deduplicated and merged. High-quality particles were selected through multiple rounds of 2D classification, ab initio reconstruction and heterogeneous refinement. With the exception of the diubiquitin dataset (Table S2), all datasets revealed co-existing open and closed conformations. We speculate that the lack of particles in closed conformation in the diUb_ATA_ structure might be due to the especially thin ice observed in the BRCA1-A:diUb_ATA_ grid. These unbiased initial models were used to (4) generate templates for 2D-based picking, and to train new picking models in (5) Topaz and (6) crYOLO. Newly picked particles were deduplicated, cleaned and merged with the initial stack.

All particles were subjected to 3D classification to separate open and closed forms, followed by global and local CTF refinement and reference-based motion correction. Global (i.e., microscope-dependent) CTF parameters were determined from the higher-resolution open-form reconstructions and then imposed on closed-form particles. After reference-based motion correction, particles were Fourier cropped to a pixel size of about 1/3 of gold standard Fourier shell correlation (GSFSC_0.143_) resolution (e.g.: GSFSC_0.143_ = 3Å would be Fourier cropped to ~1 Å/pixel). Maps deposited to EMDB are also cropped with this criterion.

Conformational heterogeneity in both open and closed classes was further resolved by focused 3D classification on the active sites. Due to intrinsic flexibility, one arm of the complex consistently displayed higher resolution than the other, and the arm tips remained poorly resolved. Detail in the low-resolution areas could be recovered by local refinement using masks on individual arms, followed by composite focused map generation in Phenix^68^. Maps used for model building were sharpened using DeepEMhancer^69^, employing the “high resolution” model. To recover maximum detail of elbow and wrist ubiquitin binding, densities in Fig. 3A and B were obtained by combining particles from the polyUb_ATA_ open structures (State C+P and double State P) and performing 3D classification to enrich for Ub occupancy at the elbow or wrist sites, followed by local refinement with a soft mask encompassing BRE, Ub_elbow_ and Ub_wrist_. C1 symmetry was used throughout the processing.

#### Model building

The final atomic model for the polyUb_ATA_, open State C+P structure was built in Coot using PDB entries 6GVW^50^ (mouse apo-BRCA1-A, open form) and 1UBQ^70^ (ubiquitin) as starting templates, with the DeepEMhancer-sharpened and/or composite focused map as the reference. Model refinement was performed against the unmasked, unsharpened 3D reconstruction using Refmac (Servalcat)^71,72^ with jelly-body restraints, and Phenix real-space refine^68^, with starting model restraints and secondary structure geometric restraints enabled.

The resulting State C+P model served as the starting point for building the polyUb_ATA_, open double State P, diUb_ATA_, and apo structures, using the same refinement workflow.

For the closed form, we performed focused 3D classification on the dimerization interface of the closed form of the BRCA1-A:oligoUb_ATA_, and obtained a class with sufficient detail to allow building of the coiled coil. We generated the pseudo atomic model by restrained flexible fitting of our AlphaFold3 prediction of BRCA1-A (ModelArchive^52^ ma-yb0lm) into the oligoUb_ATA_ map using Coot^79^, followed by local manual rebuilding where required (mainly in the dimerization interface and in the BABAM1:BRE:RAP80 interface. This model was then fitted as a rigid body in all other closed form reconstructions shown in Fig. 5 using ChimeraX^73^.

#### Pixel size correction

Some datasets displayed indications of incorrect pixel size, as evidenced by divergence of the CTF-refined spherical aberration coefficient (Cs) from the expected Titan Krios value of 2.7 mm. To estimate actual pixel size, we quantified the deviation of intra-helical hydrogen bond distances (atom N of residue *i* to atom O of residue *i+4*, Hbond_α_) compared to the expected value of 2.915⍰± ⍰0.096⍰Å^74^. We refined our atomic model without secondary structure restraints against the map at nominal pixel size using phenix real space refine, then calculated secondary structure using mkdssp^75^, and determined mean Hbond_α_ for all atoms classified as being part of alpha helix (DSSP code: H). To correct pixel scaling, we applied the formula:

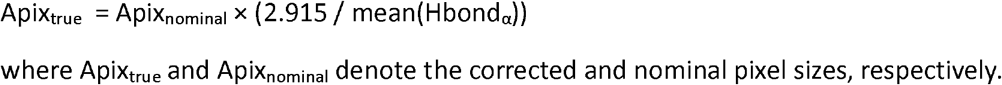

After reprocessing with new pixel size, refined Cs values for the updated reconstructions converged to 2.7 ± 1% mm, and the rescaled PDB models, refined against the corrected map, yielded a mean Hbond_α_ value within 0.5% of expected.

#### Probe restraints

Restraints for the _ATA_ probe (SMILES: NCC(S)CC(=O)NCC[C@H](N)C(=O)O) were generated using AceDRG^76^ allowing for both chiral configurations at the thiol-bearing stereocentre. Although both configurations appear geometrically compatible with either active site, Refmac Servalcat^71,72^ refinement converged on different chiralities in distinct states: the probe in State C refined to the S configuration (SMILES: NC[C@@H](S)CC(=O)NCC[C@H](N)C(=O)O), while the probe in State P refined to the R configuration (SMILES: NC[C@H](S)CC(=O)NCC[C@H](N)C(=O)O)

### AlphaFold modeling

Sequences for two copies each of ABRAXAS, BRCC36, BABAM1, BRE, and RAP80 (full-length or RAP80^271–329^), with or without up to 12 copies of ubiquitin, were provided as inputs to AlphaFold 3 (version AlphaFold-beta-20231127). Regardless of version or presence of ubiquitin, AlphaFold 2 (not shown) and Alphafold 3 predict closed structures. Residues 1– 270 and 330–777 of RAP80 are predicted to be disordered in relation to the structured core of the complex. The model with RAP80^271–329^ and 10 ubiquitin molecules was deposited in ModelArchive^52^ at https://www.modelarchive.org/doi/10.5452/ma-yb0lm

### Sequence alignments

Sequences for BRCA1-A components were retrieved from genomes of species representative of the main branches of the evolutionary tree using manual search on Uniprot or the taxon id filtering options of NCBI BLAST (https://blast.ncbi.nlm.nih.gov). Sequences were aligned using MSAProbs^77^ through the Jalview^78^ toolkit. Sequence ids: Supplementary Fig. 2C, ABRAXAS: Q6UWZ7, Q8BPZ8, Q1LVP6, Q6GR31, A0A6J0U9J7. ABRO1: Q15018, Q3TCJ1, A0A8M3AJY3, A0A8J0Q0E3, A0A6J0THY9, XP_035673382, XP_030841210, XP_006823808, XP_001813478, XP_001945617, ELT87605, XP_022288514, NP_187490, KAJ4779947, XP_004246297, AES86734, BDA41233, XP_026441550., Supplementary Fig. 2D, BRCC36: P46736, P46737, Q66GV6, A0A8M3B525, XP_072837618, XP_035694102, XP_030838197, XP_002740113, XP_064211341, E2AXC7, ELT99386, XP_022286615, NP_001117626, KAJ4771733, XP_069155468, XP_003597836, BDA48211, XP_026454415. Supplementary Fig. 3F (BRCC36 Ins-2): XM_045538942, XM_045538941, XM_045538943, NM_024332, NM_001018055, NM_001242640, NM_001166457,NM_145956, NM_001166459, NM_001358736, NM_001358737, XM_045141074, XM_045141075, XM_023248860, XM_023248861, XM_023248862, XM_023248863, XM_004285931, XM_033416850, XM_033416852, XM_033416851, NM_001435443, NM_001075790, XM_003421746, XM_003421747, XM_064277827, XM_023725641, XM_023725642, XM_004390876,XM_004390877, XM_007956641. Supplementary Fig. 4A and 4B (BRE): Q9NXR7, Q8K3W0, Q568D5, Q6GPL9, A0A6J0TNU0, B6NXD5, A0A7M7RHH1, XP_006817048, D6X0Y3, R7V964, A0A8B8DPM8, Q9FIH0, A0AAV8E530, A0A3Q7G848, I3SCW5, A0A8J9WQC4, A0AAD4X7P8.

### Surface Plasmon Resonance

#### BRCA1-A binding to Ub or polyUb chains

SPR was performed on Biacore S1+ device (Cytiva). His-BABAM1–tagged BRCA1-A complexes (WT or mutants, 0.5 mg/mL) were immobilised on a CM5 sensor chip (Cytiva) using the His Capture Kit (Cytiva) according to the manufacturer’s instructions. Runs were performed in 150 mM NaCl, 20 mM HEPES pH 7.8, 2 mM TCEP, and 0.05% Tween-20 (SPR buffer) at 25°C. Analytes (ubiquitin and K63-linked polyubiquitin chains) were buffer-exchanged into SPR buffer prior to the run using Zeba Spin Desalting Columns (0.5 mL, Thermo Fisher Scientific). Baseline correction was performed by direct subtraction of a blank cycle without analyte, followed by linear drift correction. Binding responses for Ub were normalised to the immobilisation level (RU) of each sample prior to fitting. Binding response for polyUb chains are averages from two independent experiments (normalized by expected B_max_ prior to averaging). Dissociation constants were determined in GraphPad Prism 8.0 using a one-site total binding model, imposing a shared B_max_ and setting the non-specific binding component (NS) to zero. Affinity values for chains were calculated using 5 μM as the estimated maximum concentration in the ubiquitin chain titration. Ub and polyUb chains binding curves in Fig. 5F were normalized to B_max_ to allow plotting on the same graph.

#### Estimation of Ub chain concentration

Ten microlitres of polyUb chains analyte, taken from the three highest-concentration injections of two separate SPR experiments, were digested with 1 μM BRCA1-A overnight at 37 °C. The resulting monoUb was resolved by SDS–PAGE and compared to a monoUb standard ladder. In both cases (Supplementary Fig. 4 and not shown), the highest chain concentration corresponded to ~1.3–2.6 μg/μL ubiquitin (15–30 μM). Assuming all chains are Ub_5_ (the smallest chain size in the mixture), this yields a concentration range of 3–6 μM. A value of 5 μM was used for affinity calculations. As chains are on average larger than Ub_5_, this estimate represents an upper bound for concentration and, accordingly, the derived dissociation constant is a lower-bound value.

#### GST-BRE and GST BABAM1 binding to Ub

SPR measurements were performed as described above, using a GST Capture Kit (Cytiva). Dissociation constant fitting did not converge, and the lower-bound K_d_ value was estimated from the inflection point of the binding curve.

### Structural Figures

Structural Figures were prepared with ChimeraX^73^ or Coot^79^. Cryo-EM densities shown (except in Fig. 5A) were sharpened with DeepEMhancer^69^ using the “high resolution” model and filtered to a resolution 0.2 Å lower than the corrected GSFSC resolution reported by cryoSPARC. Density colouring was performed using the Chimera “volume zone” command with the distance parameter set to 5 Å. In some cases, blur and vignette visual effects were added in Adobe Photoshop to improve clarity. ChimeraX does not render covalent bonds between the ATA probe and Gln62/Glu64 of UbP or Gly75 of UbD. Where needed, these connections were manually retouched in Adobe Photoshop to bridge the gap. These modifications are purely cosmetic and do not alter the underlying structural data.

### Declaration of AI-assisted technologies in the writing process

During the preparation of this work the authors used ChatGPT 5 (OpenAI) in order to assist in copy editing. After using this tool/service, the authors reviewed and edited the content as needed and take full responsibility for the content of the published article.

### Sequences

#### Synthetic DNA sequences

##### ABRAXAS1

ATGGAAGGCGAATCGACCTCGGCGGTTCTGAGCGGTTTCGTTCTGGGCGCACTGGCGTTTCAACATCTGAATACGGACTCGGATACGGAAGGCTTTCTGCTGGGTGAAGTGAAAGGCGAAGCGAAGAACAGCATTACCGATTCTCAGATGGATGACGTTGAAGTGGTTTATACGATTGACATCCAGAAATATATCCCGTGCTACCAACTGTTTAGTTTCTACAACAGCTCTGGTGAAGTCAATGAACAGGCCCTGAAAAAGATTCTGAGCAACGTGAAAAAGAATGTCGTGGGCTGGTATAAATTTCGTCGCCATTCTGATCAAATCATGACCTTCCGTGAACGCCTGCTGCATAAAAACCTGCAGGAACACTTTTCAAATCAAGACCTGGTGTTCCTGCTGCTGACCCCGTCGATTATCACGGAATCATGCTCGACCCATCGTCTGGAACACTCACTGTATAAACCGCAGAAGGGTCTGTTTCATCGCGTTCCGCTGGTTGTCGCAAATCTGGGTATGAGCGAACAACTGGGCTACAAAACGGTTAGCGGTTCTTGTATGAGTACCGGCTTCTCCCGCGCTGTCCAGACGCATAGTTCCAAATTTTTCGAAGAAGATGGCAGCCTGAAAGAAGTGCACAAGATTAACGAAATGTACGCAAGCCTGCAGGAAGAACTGAAATCTATCTGCAAAAAGGTTGAAGATTCTGAACAAGCTGTCGATAAACTGGTCAAGGACGTGAATCGTCTGAAACGCGAAATTGAAAAACGTCGCGGTGCGCAGATCCAAGCAGCGCGTGAAAAGAACATTCAGAAAGATCCGCAGGAAAATATCTTTCTGTGTCAGGCCCTGCGCACCTTTTTCCCGAACAGTGAATTCCTGCATTCATGCGTGATGTCGCTGAAAAATCGTCACGTTTCCAAATCATCGTGTAACTATAATCATCACCTGGATGTGGTTGACAACCTGACGCTGATGGTTGAACATACCGATATTCCGGAAGCATCACCGGCTTCGACCCCGCAGATTATCAAACACAAGGCGCTGGACCTGGATGACCGTTGGCAGTTTAAACGTAGTCGCCTGCTGGATACCCAAGACAAACGCTCCAAGGCGGATACGGGCAGCTCTAATCAGGACAAAGCCAGCAAGATGAGTTCCCCGGAAACCGATGAAGAAATTGAAAAAATGAAGGGCTTTGGTGAATACAGTCGTTCCCCGACGTT CTGA

##### BRCC3

ATGGCGGTTCAAGTCGTGCAAGCGGTTCAAGCCGTTCACCTGGAATCGGATGCGTTTCTGGTCTGTCTGAATCATGCTCTGTCTACGGAAAAAGAAGAAGTCATGGGCCTGTGCATTGGTGAACTGAACGATGACACCCGTTCTGATAGTAAATTTGCATATACCGGCACGGAAATGCGTACGGTGGCGGAAAAGGTTGACGCCGTCCGCATTGTGCATATCCACTCAGTTATTATCCTGCGTCGCTCGGATAAGCGTAAGGACCGCGTCGAAATCAGCCCGGAACAGCTGTCCGCAGCATCAACCGAAGCAGAACGTCTGGCTGAACTGACGGGCCGTCCGATGCGCGTGGTTGGTTGGTATCATAGCCACCCGCATATTACCGTTTGGCCGTCTCATGTCGATGTGCGCACGCAGGCGATGTACCAAATGATGGACCAGGGCTTCGTGGGTCTGATTTTTAGCTGCTTCATCGAAGATAAGAACACCAAGACGGGCCGTGTTCTGTACACCTGTTTTCAGAGTATCCAGGCCCAAAAAAGCTCTGAATCCCTGCACGGCCCGCGCGATTTCTGGAGTTCCTCACAACATATTAGTATCGAGGGTCAGAAAGAAGAAGAACGTTATGAACGCATTGAAATCCCGATTCACATCGTTCCGCATGTCACCATTGGTAAAGTGTGCCTGGAAAGCGCAGTTGAACTGCCGAAGATTCTGTGTCAGGAAGAACAAGATGCTTACCGTCGCATCCACAGTCTGACCCATCTGGACTCCGTCACGAAAATTCACAACGGCAGCGTGTTTACCAAGAATCTGTGTTCGCAGATGAGCGCAGTGTCTGGTCCGCTGCTGCAGTGGCTGGAAGATCGCCTGGAACAAAACCAGCAACATCTGCAGGAACTGCAGCAAGAAAAAGAAGAACTGATGCAGGAACTGTCGAGCCTGGAATG A

##### BABAM1

ATGGAAGTTGCCGAACCGTCAAGCCCGACCGAAGAAGAAGAAGAAGAAGAAGAACACTCAGCAGAACCGCGTCCGCGCACCCGCAGCAACCCGGAAGGCGCAGAAGATCGTGCTGTCGGTGCACAGGCTAGCGTGGGTAGCCGCTCTGAAGGCGAAGGTGAAGCGGCCTCTGCAGATGACGGTAGCCTGAATACGTCTGGCGCTGGTCCGAAAAGCTGGCAGGTGCCGCCGCCGGCACCGGAAGTTCAAATTCGTACCCCGCGCGTTAACTGCCCGGAAAAAGTCATTATCTGTCTGGATCTGAGTGAAGAAATGTCCCTGCCGAAACTGGAATCATTCAATGGCTCGAAGACGAACGCTCTGAATGTCAGTCAGAAAATGATTGAAATGTTCGTGCGTACCAAACATAAGATCGACAAGTCCCACGAATTTGCGCTGGTTGTGGTCAACGATGACACGGCCTGGCTGAGCGGTCTGACCTCTGATCCGCGCGAACTGTGCAGTTGTCTGTATGACCTGGAAACCGCAAGTTGCTCCACGTTCAACCTGGAAGGTCTGTTTAGCCTGATTCAGCAAAAAACCGAACTGCCGGTTACGGAAAATGTCCAGACCATTCCGCCGCCGTATGTGGTTCGTACGATCCTGGTTTACTCACGCCCGCCGTGCCAGCCGCAATTCTCGCTGACCGAACCGATGAAAAAGATGTTCCAATGTCCGTATTTCTTTTTCGATGTCGTGTACATCCACAACGGCACCGAAGAAAAAGAAGAAGAAATGAGCTGGAAGGATATGTTCGCGTTTATGGGCAGTCTGGACACCAAAGGTACGTCCTATAAGTACGAAGTGGCACTGGCAGGTCCGGCACTGGAACTGCATAATTGCATGGCAAAACTGCTGGCACACCCGCTGCAGCGTCCGTGTCAATCTCATGCCTCATACTCGCTGCTGGAAGAAGAAGATGAAGCAATCGAAGTGGAAGCTACCGTTTGA

##### BABAM2

ATGTCTCCGGAAGTGGCTCTGAACCGTATCTCGCCGATGCTGTCGCCGTTTATCTCCTCTGTCGTGCGTAATGGCAAAGTTGGCCTGGACGCAACCAACTGCCTGCGTATTACGGATCTGAAAAGCGGCTGCACCAGCCTGACCCCGGGTCCGAATTGTGACCGCTTTAAACTGCATATCCCGTATGCGGGTGAAACCCTGAAGTGGGATATTATCTTCAACGCCCAGTACCCGGAACTGCCGCCGGACTTTATTTTCGGCGAAGATGCTGAATTTCTGCCGGACCCGTCTGCACTGCAGAATCTGGCTAGTTGGAACCCGTCCAATCCGGAATGCCTGCTGCTGGTTGTGAAAGAACTGGTCCAGCAATATCACCAGTTTCAATGTAGCCGTCTGCGCGAAAGCTCTCGTCTGATGTTCGAATACCAGACCCTGCTGGAAGAACCGCAATATGGTGAAAACATGGAAATTTACGCAGGTAAAAAGAACAATTGGACCGGCGAATTTAGCGCTCGCTTCCTGCTGAAACTGCCGGTTGACTTTTCTAACATCCCGACGTATCTGCTGAAGGATGTCAATGAAGACCCGGGCGAAGATGTCGCACTGCTGTCAGTGTCGTTCGAAGATACCGAAGCTACGCAGGTTTATCCGAAACTGTACCTGAGTCCGCGTATTGAACATGCACTGGGCGGTAGTTCCGCACTGCACATTCCGGCATTTCCGGGCGGTGGCTGCCTGATCGATTATGTTCCGCAGGTCTGTCATCTGCTGACCAATAAAGTGCAGTATGTTATTCAAGGTTACCATAAGCGTCGCGAATATATCGCGGCCTTTCTGAGCCACTTCGGTACGGGCGTCGTGGAATACGATGCCGAAGGCTTTACCAAACTGACGCTGCTGCTGATGTGGAAGGATTTTTGTTTCCTGGTGCATATCGACCTGCCGCTGTTTTTCCCGCGCGATCAGCCGACCCTGACGTTTCAAAGTGTTTATCACTTCACCAACAGCGGTCAGCTGTACTCGCAGGCGCAGAAAAACTATCCGTACTCCCCGCGTTGGGATGGCAATGAAATGGCCAAACGCGCGAAGGCCTATTTTAAAACCTTCGTGCCGCAGTTTCAAGAAGCAGCTTTCGCGAATGGCAAGCTGTGA

## Supporting information

EM processing supplement

## Acknowledgements

We thank Dr Patrick Celie and the NKI Protein Facility for assistance with protein purification; Dr Xiaohu Guo and the NKI cryo-electron microscopy facility for support with cryo-EM sample preparation and screening; Ameer Alkhier, Torben Wriedt, and the NKI Research High Performance Computing Facility for help with cryo-EM compute infrastructure; Dr Alex Fish for assistance with SPR experiments; and Dr Robbie Joosten for help with refinement restraints. Dr. Gian-Luca McLelland, Dr Abdelghani Mazouzi, Dr Shun-Hsiao Lee for critical reading of the manuscript. We also thank the team at the Netherlands Center for Nanoscopy (NeCEN) for cryo-EM data collection, and all members of the NKI Department of Biochemistry for critical input and discussion. Research at the Netherlands Cancer Institute is supported by institutional grants of the Dutch Cancer Society and of the Dutch Ministry of Health, Welfare and Sport. This work was supported by Oncode Institute, the Gravity programme CGC and OCENW.M.21.288 from NWO to TKS, LSHM21048, LSHM19026 to TKS and UbiQ.

## Author contributions

F.E.O. conceived, developed, and synthesised the diUb_ATA_ and polyUb_ATA_ probes. A.G.M. conceived and performed protein purification, cryo-EM sample preparation, structure determination, and biochemical experiments. T.K.S. supervised and directed the project. A.G.M. and T.K.S. wrote the manuscript with input from all authors.

## Data availability

The cryo-EM maps and structure models were deposited to the EM Data Bank (EMDB) and Protein Data Bank respectively, under accession codes: EMD-55040, pdb_00009SMR (open Apo BRCA1-A); EMD-55055 (closed Apo BRCA1-A); EMD-55047, pdb_00009SNA (open BRCA1-A:diUb_ATA_ complex); EMD-55054 (open BRCA1-A:oligoUb_ATA_ complex); EMD-55053, pdb_00009SO9 (closed BRCA1-A:oligoUb_ATA_ complex state C*); EMD-55038, pdb_00009SMN (open BRCA1-A:polyUb_ATA_ complex, StateC+P); EMD-55039, pdb_9SMP (open BRCA1-A:oligoUb_ATA_ complex, state double P). The Alphafold3 model of full-length human BRCA1-A in complex with monoUb is available in ModelArchive with accession code ma-yb0lm.

