## Supplementary Figures for "A ubiquitin chain-feeding mechanism for BRCA1-A"

### Supplementary Figure 1

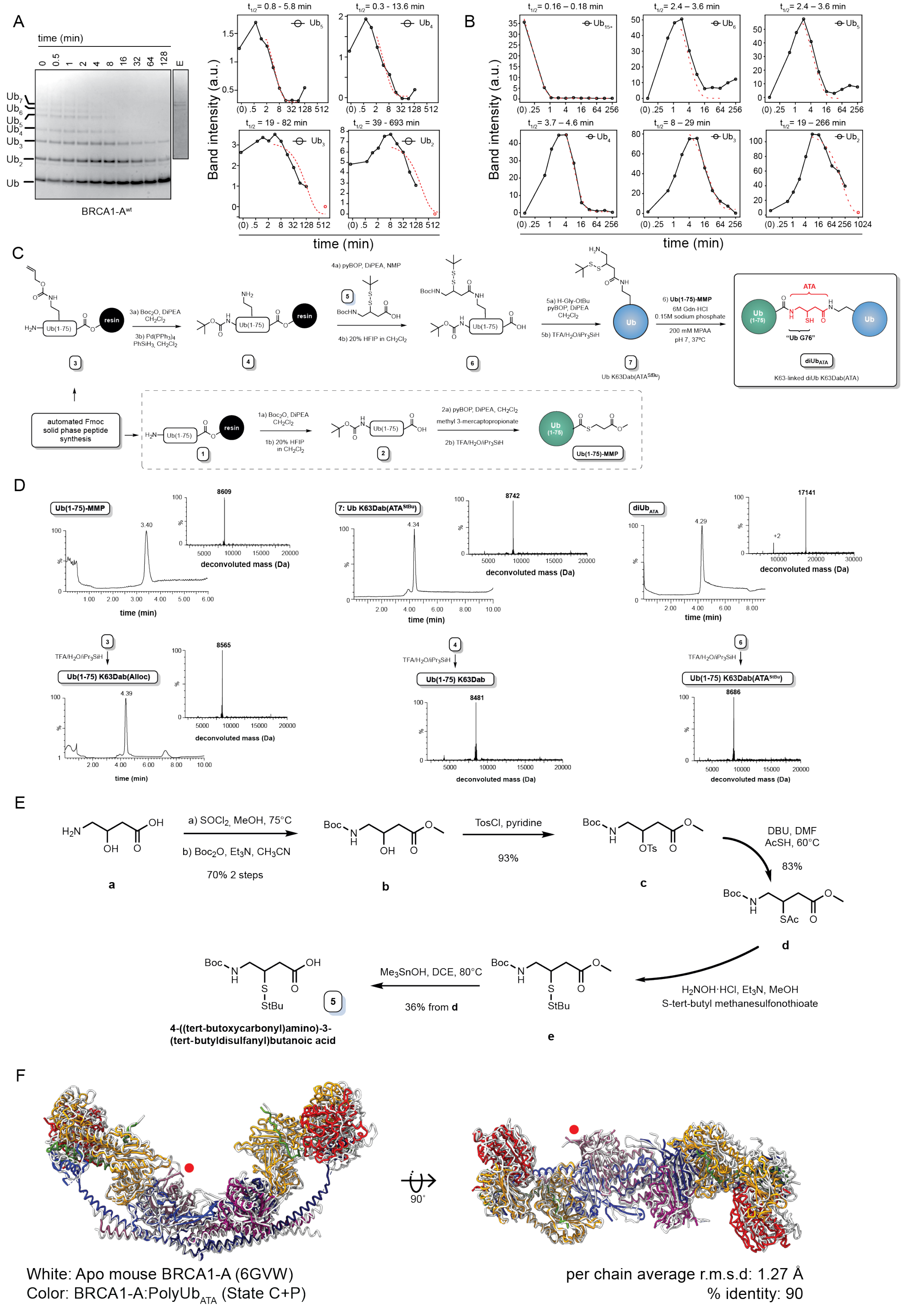

#### Supplementary Figure 1: Characterisation of BRCA1-A activity, synthesis of $diUb_{ATA}$ and comparison of human and mouse BRCA1-A.

**(A) BRCA1-A processes shorter K63-linked chains in order of decreasing length.** Left: intermediate- and short-length K63-linked Ub chains ($Ub_{2-5}$) were digested with 5 nM BRCA1-A in a time course, and products at the indicated time points were resolved by SDS–PAGE and stained with Coomassie. Lane E: 1 µl of 1 µM enzyme stock was loaded as control. Right: band intensities corresponding to each species were quantified by densitometry and plotted. $Ub_{4-5}$ chains are digested first, followed by $Ub_3$ and $Ub_2$ at a slower rate. To estimate half-lives, data points at and after the maximum of each curve were fitted with a single-exponential decay model with plateau, with the simplifying assumption that accumulation rates for each species are null after reaching plateau. In order to aid fitting of $Ub_3$ and $Ub_2$ curves, levels were assumed to reach zero at t = 1024 min (~17h). (B). Apparent half-life of each species (t½) is shown as 95% confidence interval. Black: experimental data. Red dashed line: fitted curve.

**(B) Estimation of decay rates for ubiquitin species in Fig. 1B.** The resulting half-lives ($t_{1/2}$) are reported with 95% confidence intervals. The Akaike information criterion was used to assess whether $t_{1/2}$ differed significantly between Ub species. Long ($Ub_{15+}$), intermediate ($Ub_{4–6}$), and short chains ($Ub_{2–3}$) were significantly different (long vs. intermediate: >99.9%, ΔAIC = 68.21; intermediate vs. short: >99.9%, ΔAIC = 31.37). Within the intermediate group, $t_{1/2}$ $Ub_4$ was significantly different from $Ub_{5-6}$ (96.66%, ΔAIC = 6.73). $Ub_2$ and $Ub_3$ had significantly different $t_{1/2}$ (99.64% support, ΔAIC = −5.735). $Ub_2$ levels were assumed to reach zero at t = 1024 min (~17h). Black: experimental data. Red dashed line: fitted curve.

**(C) Synthesis of the $diUb_{ATA}$ probe.** Synthetic scheme for $diUb_{ATA}$. Synthesis of **5** is shown in panel (E). Full synthesis details are provided in Materials and Methods.

**(D) LC-MS analysis of the intermediates and products from (C).** Samples for LC-MS analysis were obtained by treating a small amount of fully protected material (2-5 mg) with 50 – 75 uL TFA/$H_2O$/iPr$_3$SiH (92/5/3 v/v/v) for 2.5 hrs. Next, 1 mL diethyl ether was added and the precipitated material dissolved in 1 mL $CH_3CN$/$H_2O$/formic acid (50/50/1 v/v/v) for LC-MS analysis. Gradient: 30-80% B over 6.5 min (over 3.5 min for **Ub(1-75)-MMP**)

**(E) Chemical synthesis of ATA building block 5.** Full experimental details and analyses are provided in Mulder *et al.*, 2014[1] (open access).

**(F) Human BRCA1-A core adopts a conformation nearly identical to mouse BRCA1-A.** The human BRCA1-A complex in complex with $polyUb_{ATA}$ (State C+P, in colour) was superposed onto the mouse BRCA1-A complex (PDB ID: 6GVW[2], in white) using secondary structure matching (SSM) alignment in Coot[3], with BRCC36 (chain B) as the reference (indicated by red dot). Two views related by a 90° rotation are shown. The chain-to-chain average root mean square deviation (r.m.s.d.) across all Cα atoms is 1.27 Å.

Supplementary Figure 2

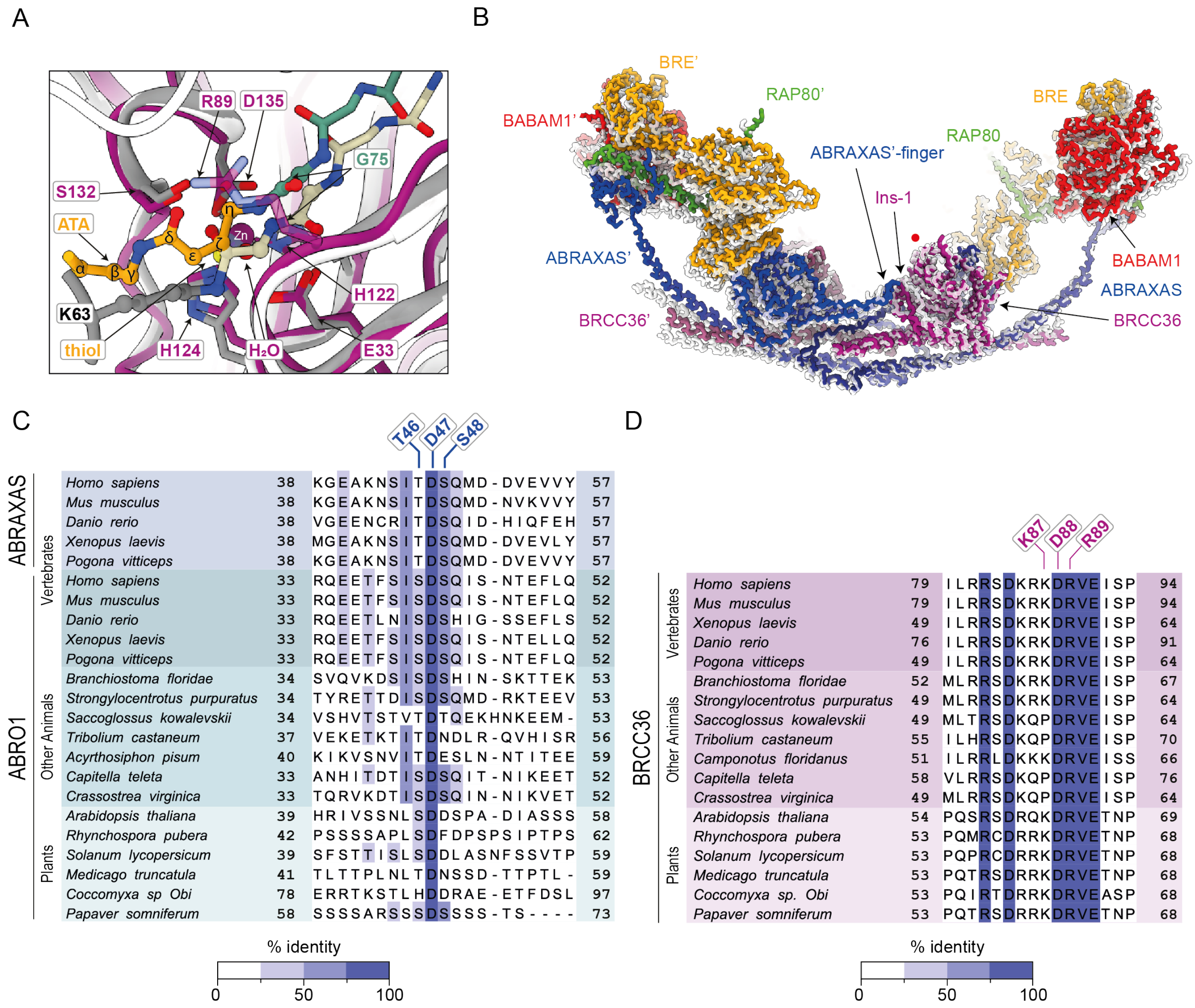

#### Supplementary Figure 2: Changes in active site upon substrate engagement.

**(A) The probe warhead mimics the native isopeptide bond.** Close-up of the BRCC36 active site (purple, State C) in complex with polyUb$_{ATA}$ (UbD: teal; ATA: orange), superposed to apo-AMSH-LP (PDB: 2ZNR[4] light grey), and to catalytically inactive AMSH-LP E292A (white) in complex with K63-diUb (UbD: beige; UbP K63: grey; PDB: 2ZNV[4]). Superposition was performed between BRCC36 and AMSH-LP using ChimeraX. Catalytically important residues of BRCC36 (His124, His122, Asp135, Glu33, Ser132) align closely with the corresponding residues of apo-AMSH-LP (His347, His349, Asp360, Glu292, and Ser357) and adopt a catalytically competent configuration. The C-terminus of UbD in our structure adopts a similar conformation to that of inactive AMSH-LP bound to K63-diUb. The thiol of the ATA probe chelates the zinc ion and occupies the position corresponding to the catalytic water, while the ζ-carbon of ATA correctly mimics the scissile carbon. The portion of ATA corresponding to the lysine side chain follows a different trajectory in the two structures, reflecting the distinct positioning of UbP in AMSH-LP and BRCC36. The carbonyl group of ATA interacts with the side chains of BRCC36 Ser132 and Arg89. The precise positioning of the native carbonyl group in the isopeptide bond remains unclear, as the AMSH-LP G292A mutant used for the diUb complex lacks the catalytic zinc, resulting in an apparent positioning of the isopeptide bond too close to the His-His-Asp triad. Given that Ser132 is thought to form the oxyanion hole that stabilizes the reaction intermediate, it is likely that, in the native state, the isopeptide carbonyl also interacts with Ser132 and Arg89 of BRCC36. Residue numbering is based on BRCC36.

**(B) Substrate engageme nt brings ABRAXAS and the opposing BRCC36 into closer proximity**. Apo- (white) and BRCA1-A:PolyUb$_{ATA}$ (State C+P, colour) BRCA1-A structures—shown as backbone traces—were superposed on the BRCC36 subunit in state C (indicated by the red dot). Engagement of BRCC36 Ins-1 with the opposing ABRAXAS-finger brings the two proteins into closer proximity, inducing slight bending of the coiled coil and partial closure of the arms. Ubiquitins are omitted for clarity.

**(C) The length and tip of the ABRAXAS-finger are conserved between ABRO1 and ABRAXAS, and in eukaryotes.** Sequences of ABRAXAS and ABRO1 from selected eukaryotic species were aligned using MSAProbs[5]. The region corresponding to the ABRAXAS (or ABRO1) finger is shown, coloured by percentage identity. In vertebrates, both ABRAXAS (BRCA1-A) and ABRO1 (BRISC) sequences are displayed; in other eukaryotes, which lack ABRAXAS, only ABRO1 is shown. The aspartic acid at the tip of the finger (Asp47 in human ABRAXAS) is conserved across all species.

**(E) The BRCC36 Ins-1 residues interacting with the ABRAXAS-finger are conserved in eukaryotes.** BRCC36 sequences from the same species in (B) were aligned using MSAProbs[5]. The Ins-1 region is shown, coloured by percentage identity. Arg89 is invariant in all species examined.

Supplementary Figure 3

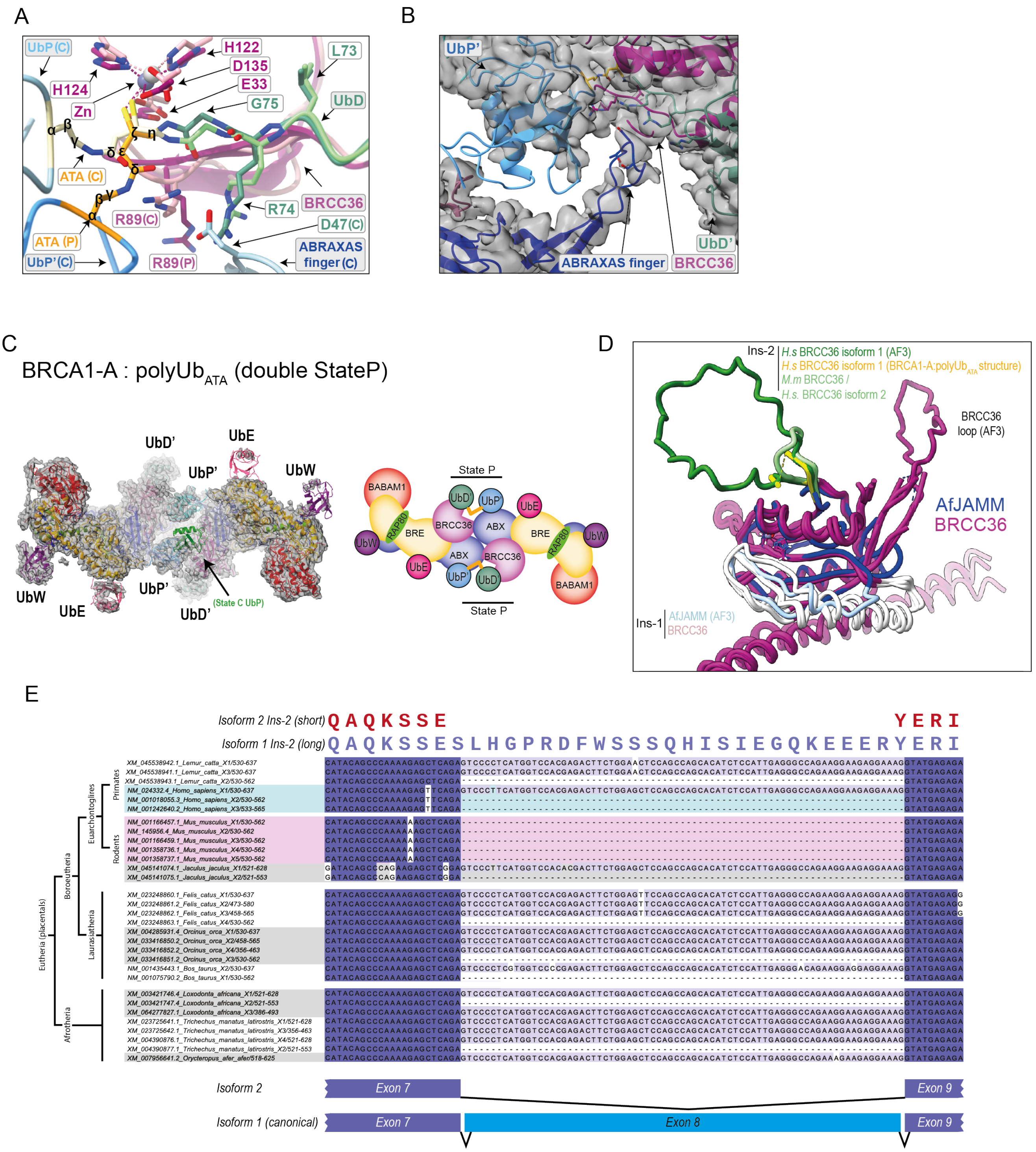

#### Supplementary Figure 3: Details of active site in State P and role of Ins-2.

**(A) Comparison of active site configuration between State C and State P.** The active sites in state C and state P were superposed via BRCC36 (State C: pink; State P: purple). No differences are observed in the active site of BRCC36 or the C-terminus of UbD (State C: light green; State P: dark green) between the two states. However, owing to the different conformation of UbP' in state P, the probe bends away from the active site by approximately 55 degrees at the thiol-bearing ζ-carbon. Proteins are shown in stick representation. Carbons of ATA are marked with Greek letters following the order of Fig. 1C. UbP and residues that differ in position between State C and State P are labelled with the corresponding state in parentheses.

**(B) The ABRAXAS-finger can interact with UbD in State P**. Particles for open BRCA1-A:polyUb$_{ATA}$ (State C+P) and open BRCA1-A:polyUb$_{ATA}$ (double State P) were combined, and focused classification was applied to the UbD':BRCC36':ABRAXAS interface of the active site in State P. A subclass is shown where the active site sees density for the ABRAXAS-finger extending towards the C-terminal arginines of UbD'.

**(C) Both active sites can simultaneously exist in state P.** Left: Sharpened electron density and fitted model of BRCA1-A with both active sites in state P (BRCA1-A:polyUb$_{ATA}$ double State P, Supplementary Table 1). A proximal ubiquitin in state C has been added in green for comparison. Right: schematic representation of the structure. ABX: ABRAXAS.

**(D) BRCC36 isoforms have short or long insertion 2**. Superposition of the crystal structure of *Af*JAMM (PDB:1R5X[6], blue) with the cryoEM structure of BRCA1-A:polyUb$_{ATA}$ (BRCC36 isoform 1 shown), and the crystal structure of mouse BRCC36 (PDB: 6GVW[1], equivalent to human BRCC36 isoform 2, purple) shows the location of Ins-1 (white) and Ins-2 (green). AfJAMM Ins-1 and BRCC36 Isoform 1 Ins-2 are disordered in the structure and are shown as predicted by Alphafold3. BRCC36 (human and mouse) bears a unique loop (BRCC36 loop) not present in *Af*JAMM (also modelled with Alphafold3). *H.s.: Homo sapiens; M.m: Mus musculus; Af: Archaeoglobus fulgidus;* AF3: AlphaFold 3.

**(E) BRCC36 exists with short and long insertion 2 in many placental mammals, but not mouse.** All BRCC36 transcripts for the indicated mammalian species were retrieved from NCBI, aligned using MSAProbs[5], and coloured by percentage identity. A portion of the alignment is shown, centred around Ins-2. Species are grouped according to taxonomy. Transcripts containing a long Ins-2 include an extra exon (exon 8 in humans) and are found across most major placental clades (highlighted in blue), demonstrating inheritance from a common ancestor. Although a long Ins-2 is present in some rodents (e.g. lesser jerboa, *Jaculus jaculus*), it is absent in mouse (*Mus musculus*, highlighted in pink). X# indicates the transcript number as defined by NCBI. Human and mouse transcripts are highlighted in blue and red, respectively.

### Supplementary Figure 4

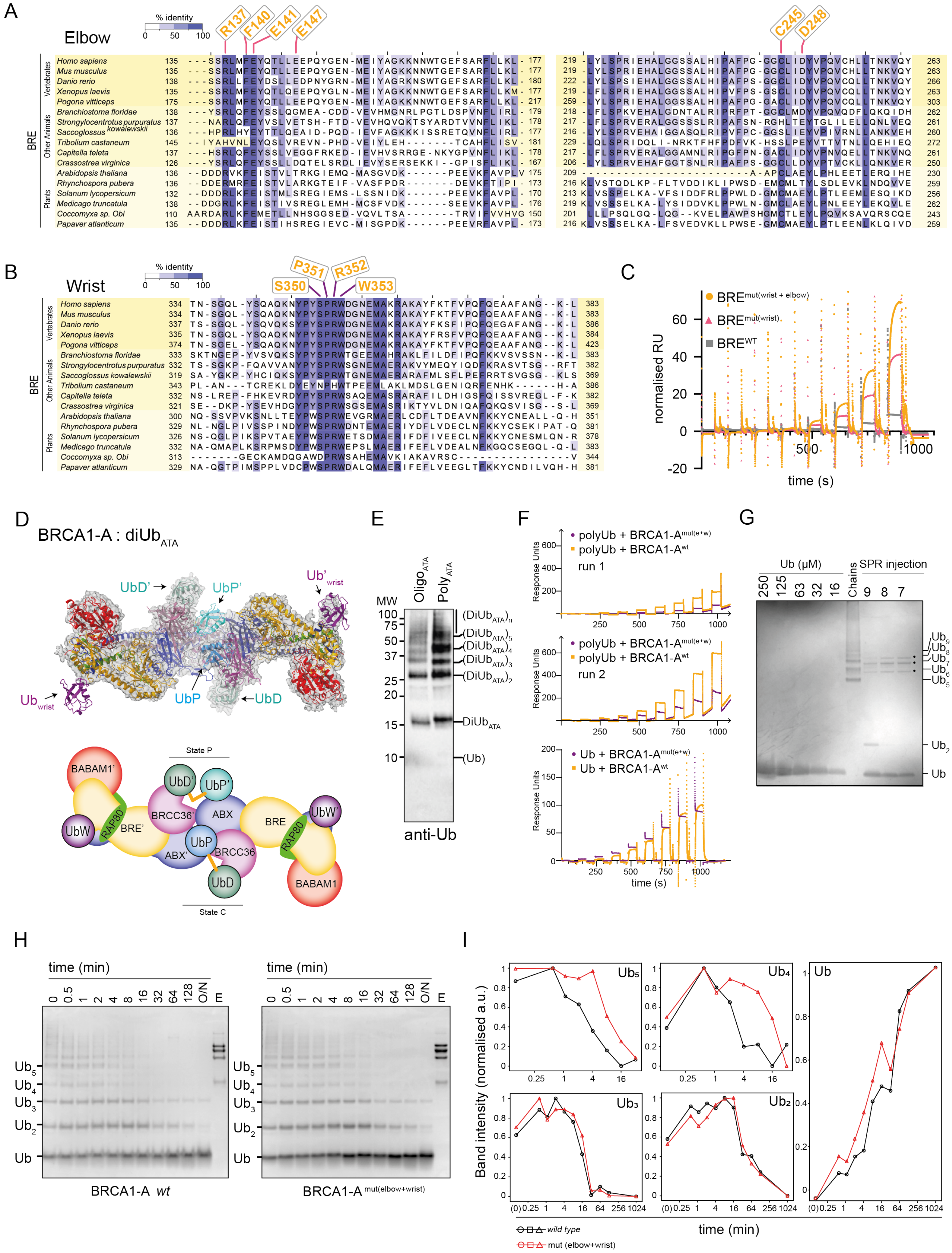

#### Supplementary Figure 4: BRE wrist and elbow site display conserved interfaces to ubiquitin.

**(A) BRE $Ub_{elbow}$ binding site is conserved across eukaryotes**. Sequences of BRE from selected eukaryotic species were aligned using MSAProbs[5]. The interfaces of BRE on helix 4 (left) and helix 5 (right) that bind $Ub_{elbow}$ are shown. The residues homologous to human Arg137 and Phe140 are conserved across eukaryotes. Main interacting residues are indicated.

**(B) BRE $Ub_{wrist}$ binding site is conserved across eukaryotes**. Same as in (A), but the region corresponding to the $Ub_{wrist}$ binding site is shown, coloured by percentage identity. The key Ub binding residues of BRE are conserved across all species.

**(C) The BRE wrist site accounts for most of the binding affinity of BRE for ubiquitin.** SPR response curves normalized to immobilization levels for the data shown in Fig. 4C. RU: response units.

**(D) Structure of BRCA1-A with $diUb_{ATA}$ displays Ub bound to the wrist site.** Top: Combined focused map and model of the BRCA1-A:$diUb_{ATA}$ structure (Supplementary Table 1). Both active sites of BRCA1-A are occupied by $diUb_{ATA}$ (state C+P). Additionally, a Ub moiety is observed at each wrist site (circled). Bottom: Schematic representation of the structure in D.

**(E) Oligo- and $polyUb_{ATA}$ do not contain detectable monoUb.** One microliter of oligo- or $polyUb_{ATA}$ were analysed by anti Ub western blot. The expected migration height for Ub is marked.

**(F) Response curves for the SPR analysis in Fig. 4F.** Top and middle panel: baseline-corrected binding responses from two experiments comparing BRCA1-A and BRCA1-$A^{mut(wrist+elbow)}$ binding to polyUb chains. Bottom panel: baseline-corrected binding responses of BRCA1-A and BRCA1-$A^{mut(wrist+elbow)}$ for monoUb.

**(G) Estimation of the concentration of Ub chains used in the SPR experiment of Fig. 4F.**
Ten microliters of K63 polyUb chains (analyte) from each of the 3 highest-concentration points of the SPR experiment were digested overnight with 1µM BRCA1-A. The digestion was loaded on SDS page and the amount of digested Ub was compared against a standard of purified Ub. 10 ul of undigested chains were loaded as a control. The total amount of monoUb digested in the highest concentration SPR sample is estimated to be 1.3 – 2.6 µg, equivalent to 0.13 – 0.26 µg/µl, or ~15 - 30 µM monoUb) . Conservatively assuming that all chains were $Ub_5$ leads to a concentration estimation of 3 – 6 µM. The value of 5 µM was used in the fitting of Fig. 4F. Black dots indicate the bands of the BRCA1-A enzyme used in the digestion. The migration height of some Ub species is indicated to the side of the graph.

**(H) BRCA1-$A^{mut(elbow+wrist)}$ displays wild-type activity toward short ubiquitin chains.** The activity of BRCA1-A WT and BRCA1-$A^{mut(elbow+wrist)}$ against short K63-linked ubiquitin chains (≤$Ub_{2-5}$) was assayed in a time course and analysed by SDS–PAGE. Lane E: five microliters of the 1 µM enzyme stocks used in the experiment were loaded as a control.

**(I) Quantification of the Ub species in (H).** Band intensities corresponding to the different ubiquitin species in (H) were quantified by densitometry and plotted over time. BRCA1-$A^{mut(elbow+wrist)}$ digests $Ub_4$ and $Ub_5$ at a slower rate compared to the *wild type* control, but exhibits comparable digestion rates for $Ub_2$ and $Ub_3$. Time is shown on a $log_2$ scale. Overnight digestion was assumed to correspond to 1024 min (~17 h). Time 0 was approximated as $2^{-4}$ minutes. A.u., arbitrary intensity units; O/N: overnight. All curves were independently normalised to their respective minima and maxima.

### Supplementary Figure 5

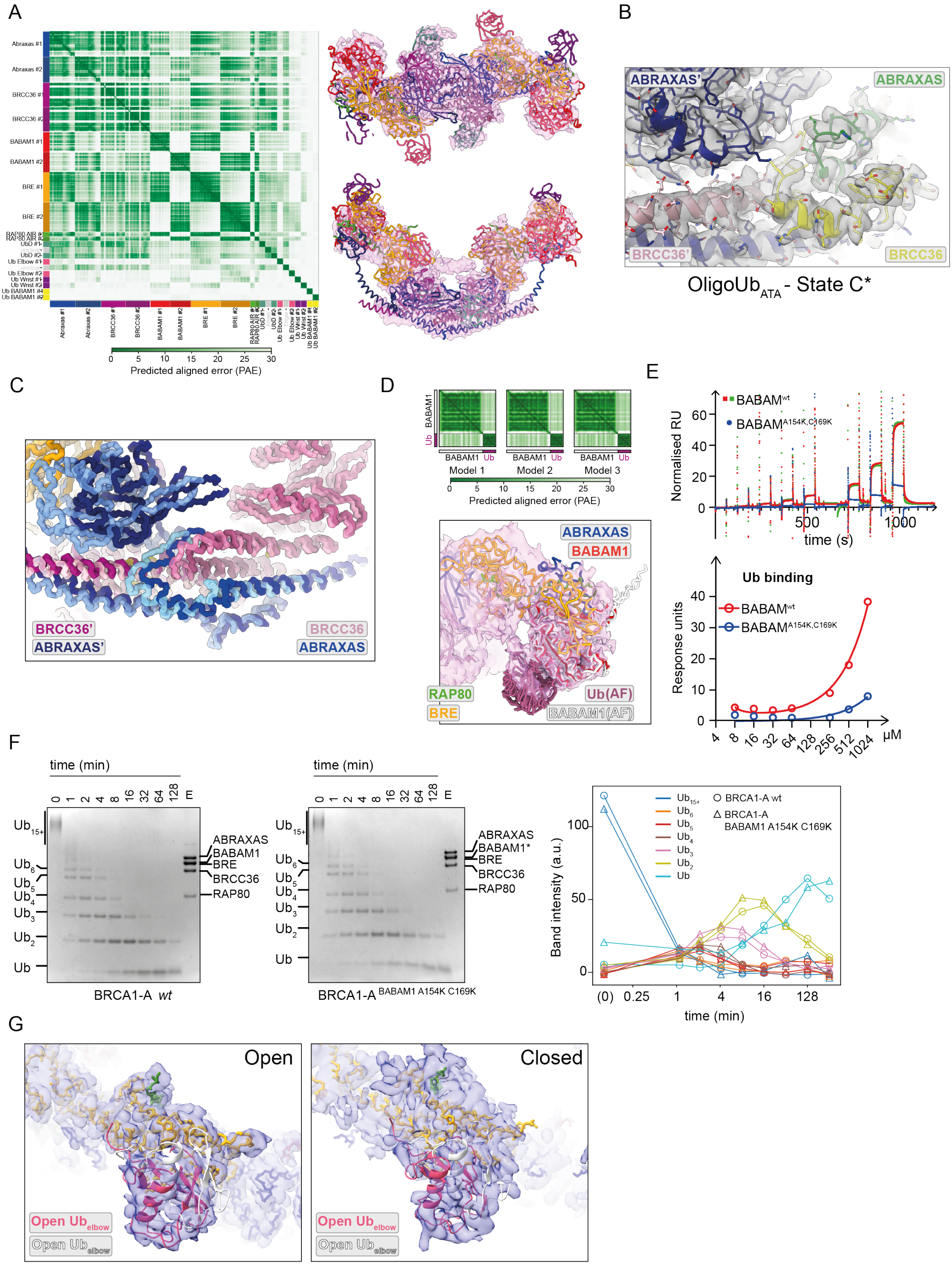

#### Supplementary Figure 5:
#### BRCA1-A exists in a closed form with specific ubiquitin binding properties.

**(A) AlphaFold 3 (AF3) predicts BRCA1-A in a closed conformation with ubiquitin anchoring points along the arm.** AF3 was used to model the full-length human BRCA1-A complex in the presence or absence of ten copies of monoUb. AF3 predicts BRCA1-A in a closed form, irrespective of Ub binding (not shown). When Ub is added, AF3 correctly places $Ub^{wrist}$, $Ub^{elbow}$, and UbD, but does not position UbP. AF3 also positions an Ub in front of BABAM1 in the same conformation shown in (C), but with a very low pAE score. Two ubiquitins are confidently predicted to bind the solvent-exposed side of the ABRAXAS-finger, adjacent to UbD, although we see no evidence for such binding. Left: Predicted aligned error (pAE) matrix, with chain assignments annotated. Centre: Rigid-body fitting of the AF3 model into the experimental density of the closed conformation of BRCA1-A:oligo$Ub_{ATA}$ (pink). The AF3 model of BRCA1-A:$Ub_{12}$ was deposited at ModelArchive with ID: ma-yb0lm.
**(B) Electron density map at dimerization interface of the closed form.** Close up of the dimerization interface of the closed form of BRCA1-A:oligo$Ub_{ATA}$ (Fig. 5B). DeepEMHancer[7]-sharpened density and fitted model are shown.
**(C) Closure of BRCA1-A brings changes in the dimerization interface.** Close-up of the dimerization region from Figure 5C. Open BRCA1-A:poly$Ub_{ATA}$ (light colours) and closed BRCA1-A:oligo$Ub_{ATA}$ (dark colours) were superposed, with main chains shown and carbonyl atoms omitted for clarity. Arm closure brings ABRAXAS' closer to BRCC36, leading to a rearrangement of the ABRAXAS and BRCC36 coiled coils at the dimerization interface.
**(D) BABAM1 binds ubiquitin.** Top: pAE matrix for the AF3 prediction of the ubiquitin:BABAM1 complex (three of five models). Ubiquitin binding to the solvent-exposed face of BABAM1 is predicted in three models, while the remaining two erroneously predict Ub binding at the BRE:BABAM1 interface (not shown). Bottom: The three AF3 predictions were superposed onto the model of closed BRCA1-A:oligo$Ub_{ATA}$ in Fig. 5A, via BABAM1, and fitted into the unsharpened density map of a closed-form subclass bearing extra density in front of BABAM1 (59,382 particles).
**(E) Mutations in the AF3-predicted Ub binding site of BABAM1 disrupt binding.** SPR analysis of binding of Ub to His-tagged BABAM1 (wild-type or double mutant A154K/C169K) immobilised via an anti-His antibody. Top: response curves normalised to immobilisation level (two lanes BABAM1 WT, green and red; one lane $BABAM1^{A154K/C169K}$, blue). Bottom: response levels (BABAM1 WT, average of two lanes, red; $BABAM1^{A154K/C169K}$, blue). Although the titration does not reach saturation, binding is estimated to occur in the 0.5–1 mM range. Mutation of the predicted interface residues ($BABAM1^{A154K/C169K}$) shows a very strong loss of interaction.
**(F) Loss of ubiquitin binding by BABAM1 does not impair catalytic activity.** Left: BRCA1-A complexes assembled with either wild-type BABAM1 or the A154K/C169K mutant were tested for their ability to cleave K63-linked polyUb chains. Right: Band intensities corresponding to the different ubiquitin species were quantified by densitometry and plotted over time. Digestion rates for all Ub species are comparable between the two BRCA1-A variants. Time is shown on a $\log_2$ scale. Time 0 was approximated as $2^{-4}$ minutes and notated as (0). A.u., arbitrary intensity units.
**(G) $Ub_{elbow}$ shifts forward upon arm closure.** The structure of open BRCA1-A:poly$Ub_{ATA}$ (state C+P) and the AF3 model of BRCA1-A:$Ub_{12}$ from (A) were fitted in the density for open BRCA1-A:poly$Ub_{ATA}$ (left) and closed BRCA1-A:poly$Ub_{ATA}$ (from Fig. 5E, right) using BRE. The $Ub_{elbow}$ configuration observed in the open form of BRCA1-A:poly$Ub_{ATA}$ does not fit the closed-form $Ub_{elbow}$ density because the density has shifted forward; the reverse is true for the AF3-predicted configuration of $Ub_{elbow}$.

#### Supplementary Figure 6

A

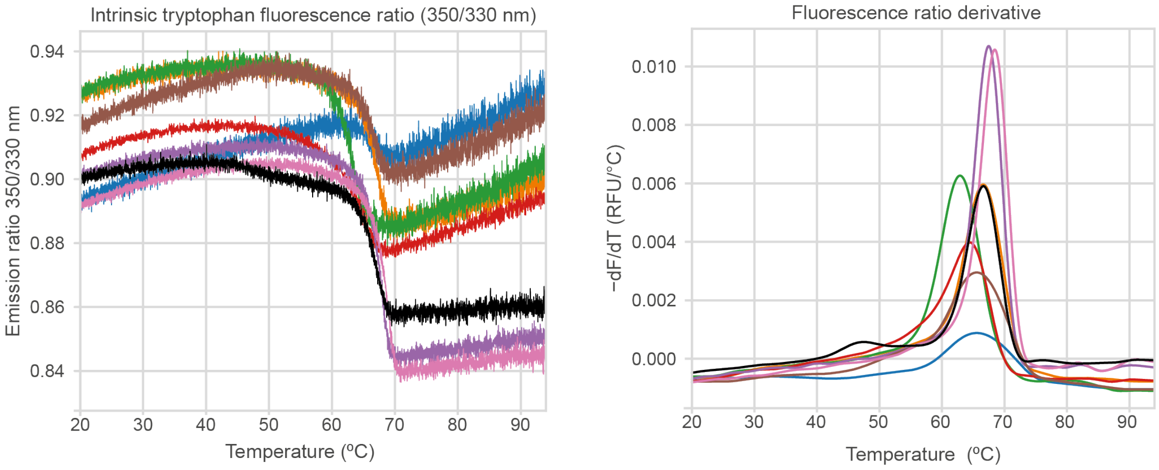


**Legend:**

| BRCA1-A variant | $T_m$ ± st.dev. | BRCA1-A variant | $T_m$ ± st.dev. |
|---|---|---|---|
| BRCA1-A (BRE mut wrist) | 65.6 ± 1.4 °C | BRCA1-A (BRCC36 isoform 2) | 65.1 ± 1.4 °C |
| BRCA1-A (BABAM1 A154K C169K) | 65.7 ± 0.1 °C | BRCA1-A (BRCC36 R89A) | 66.0 ± 0.7 °C |
| BRCA1-A (BRE mut elbow+wrist) | 62.8 ± 0.1 °C | BRCA1-A (BRE W165A) | 67.7 ± 0.7 °C |
| BRCA1-A *wild type* | 65.6 ± 0.9 °C | BRCA1-A (ABRAXAS 3A) | 64.0 ± 1.7 °C |

B

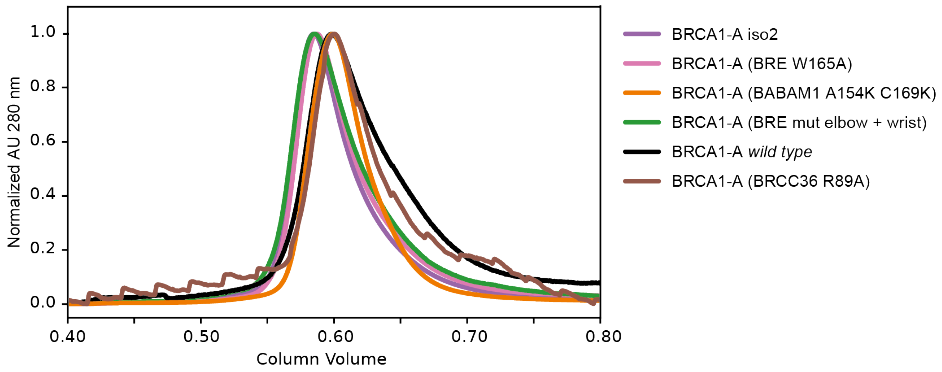


##### Supplementary Figure 6: BRCA1-A variants are stable and well folded.

**(A) BRCA1-A variants have melting temperatures comparable to wild type.** The indicated BRCA1-A variants were diluted to 1–5 µM in 150 mM NaCl, 20 mM HEPES (pH 7.5), and 5 mM TCEP, and their thermal stability was assessed using a Prometheus NT.48. Thermal denaturation (20–95 °C, gradient 1 °C/min) was monitored by intrinsic tryptophan fluorescence at 330 and 350 nm. Left: raw melting curves. Right: first derivatives of the melting curves. Table: calculated melting temperatures (mean ± standard deviation, n ≥ 2). **(B) BRCA1-A variants are well folded.** 30–100 µg (25 µl) of the indicated BRCA1-A variants were loaded onto a Superose 6 Increase 3.2/300 size-exclusion column. Absorbance at 280 nm is shown, normalized to peak intensity to account for differences in protein amount. All mutants elute at similar volumes, indicating comparable size and proper folding.
