## Supplementary Table 1 for "A ubiquitin chain-feeding mechanism for BRCA1-A"

**Cryo-EM data collection, refinement and validation statistics**

|  | Apo* | BRCA1-A  DiUb_ATA_ | BRCA1-A  OligoUb_ATA_  (closed) | BRCA1-A  PolyUb_ATA_ State C+P | BRCA1-A  PolyUb_ATA_  State double P | Ub_wrist_ focused refinement |
| --- | --- | --- | --- | --- | --- | --- |
| State | Open | Open | Closed | Open | Open | Open |
| EMDB id | EMD-55040 | EMD-55047 | EMD-55053 | EMD-55038 | EMD-55039 | EMD-55041 |
| PDB id | 9SMR | 9SNA | 9SO9 | 9SMN | 9SMP | 9SMS |
| **Data collection and processing** |  |  |  |  | | |
| Magnification | 81000 | 105,000 | 105,000 | 105,000 | | |
| Voltage (kV) | 300 | 300 | 300 | 300 | | |
| Electron exposure (e–/Å^2^) | 50 | 50 | 50 | 50 | | |
| Defocus range (μm) | -1.2-2.5 | -1.2-2.5 | -1.2-2.5 | -1.2-2.0 | | |
| Pixel size (Å)** | 0.56 (1.12) | 0.418 (0.836) | 0.418 (0.836) | 0.418 (0.836) | | |
| Symmetry imposed | C1 | C1 | C1 | C1 | | |
| Initial particle images (no.) | ~ 2 million | ~1 million | ~2 million | ~ 2 million | | |
| Final particle images (no.) | 278,973 | 195,047 | 74506 | 113,023 | 134,835 | 123929 |
| Map resolution (Å) | 3.2 | 3.1 | 3.4 | 3.2 | 3.1 | 3.3 |
| FSC threshold | 0.143 | 0.143 | 0.143 | 0.143 | 0.143 | 0.143 |
| Map resolution range (Å)*** | 4.1 – 2.9 | 4.2 - 2.8 | 11.9 – 3.0 | 3.5 - 2.8 | 3.3 - 2.8 | 3.3 – 2.8 |
| **Refinement** |  |  |  |  |  |  |
| Initial model used (PDB code) | 9SMP | 9SMP | ModelArchive  ma-yb0lm | 6GVW  1UBQ | 9SMN | 9SMP |
| Model resolution (Å)  FSC threshold | 3.2/3.2/3.6  0/0.143/0.5 | 3.1/3.2/3.8  0/0.143/0.5 | 3.3/3.4/6.6  0/0.143/0.5 | 3.1/3.3/3.8  0/0.143/0.5 | 3.0/3.2/3.7  0/0.143/0.5 | 2.9/3.1/3.7  0/0.143/0.5 |
| Map sharpening *B* factor (Å^2^) | 102.9 | 85.6 | 71.1 | 47.3 | 57.8 | 63.1 |
| Model composition  Non-hydrogen atoms  Protein residues  Ligands | 19528  2436  2 | 23487  2930  4 | 21051  2624  3 | 24768  3093  4 | 24573  3067  4 | 6410  800  0 |
| *B* factors (Å^2^)  Protein (min/max/mean)****  Ligand (min/max/mean) | 18.51/634.53/  167.10  125.62/160.35/  142.99 | 17.84/1009.3/191.37  48.61/177.31/109.17 | 30.00/999.99/  200.18  121.33/147.78/  135.76 | 13.33/479.81/  141.61  108.86/150.25/  129.88 | 12.43/1042.00/  156.13  84.97/143.94/  117.51 | 25.62/326.27/  133.06  - |
| R.m.s. deviations  Bond lengths (Å)  Bond angles (°) | 0.005  1.046 | 0.006  0.980 | 0.009  1.316 | 0.015  1.468 | 0.014  1.550 | 0.012  1.396 |
| Validation  MolProbity score  Clashscore  Poor rotamers (%) | 2.02  10.64  1.28 | 1.72  10.02  0.04 | 2.36  26.50  0.25 | 2.25  18.63  1.26 | 2.25  22.50  0.76 | 2.61  17.33  3.04 |
| Ramachandran plot  Favored (%)  Allowed (%)  Disallowed (%) | 94.09  5.82  0.08 | 96.77  3.19  0.03 | 92.99  7.01  0.00 | 93.82  6.15  0.03 | 93.95  6.02  0.03 | 91.62  8.25  0.13 |

^* This grid was frozen in presence of OligoUbATA but no ligand is present in the active site, although Ub is observed at the wrist site.
The active site appears aligned and intact.^

^** Super-resolution mode. Value in parenthesis indicates pixel size after in-camera binning.^

^*** Calculated by Cryosparc orientation diagnostics, with mask auto tightening.^

^**** Density for Ub^_elbow_^, Ub^_wrist_ ^and occasionally BABAM1 and UbP is clearly present but presents much lower resolution due to arm movement. These moieties were fitted as rigid bodies with 100% occupancy, resulting in elevated atomic displacement parameters (ADPs).^
