## Supplementary material for "A ubiquitin chain-feeding mechanism for BRCA1-A": EM processing supplement

### Table 1: Deposited structures and features of each dataset

| Dataset Apo (ligand: oligoUb$_{ATA}$) | | | | | | | |
|---|---|---|---|---|---|---|---|
| Map ID | Structure name | Ub$_{wrist}$ | Ub$_{elbow}$ | Active site 1 | Active site 2 | Arms | PDB |
| EMDB: 55040 | Apo | Present | Absent | Empty, aligned | Empty, aligned | Open | 9SMR |
| EMDB: 55055 | | Present | Absent | Empty | Empty | Closed | |
| **Dataset diUb (ligand: diUb$_{ATA}$)** | | | | | | | |
| EMDB: 55047 | diUb$_{ATA}$ | Present | Absent | State C | State P | Open | 9SNA |
| **Dataset oligoUb (ligand: oligoUb$_{ATA}$)** | | | | | | | |
| EMDB: 55054 | | Present | Weak | State C | UbP disordered | Open | |
| EMDB: 55053 | oligoUb$_{ATA}$ | Present | Weak | State C* | UbP disordered | Closed | 9SO9 |
| **Dataset polyUb (ligand: polyUb$_{ATA}$)** | | | | | | | |
| EMDB: 55038 | State C<br>State P | Present | Present | State C | State P | Open | 9SMN |
| EMDB: 55039 | Double P | Present | Present | State P | State P | Open | 9SMP |
| EMDB: 55056 | | Present | Present | State C* | UbP disordered | Closed | |
| Focused maps from this dataset: | | | | | | | |
| EMDB: 55053<br>PDB: 9SO9 | Ub$_{wrist}$ focused classification and refinement<br>(merging particles from open State C + State P, and open double State P structures) | | | | | | |
| EMD-55052 | Ub$_{elbow}$ focused classification and refinement<br>(merging particles from open State C + State P, and open double State P structures) | | | | | | |

### References:

1 - https://guide.cryosparc.com/processing-data/all-job-types-in-cryosparc/utilities/job-orientation-diagnostics

Figure 1: BRCA1-A polyUb$_{ATA}$ dataset processing workflow

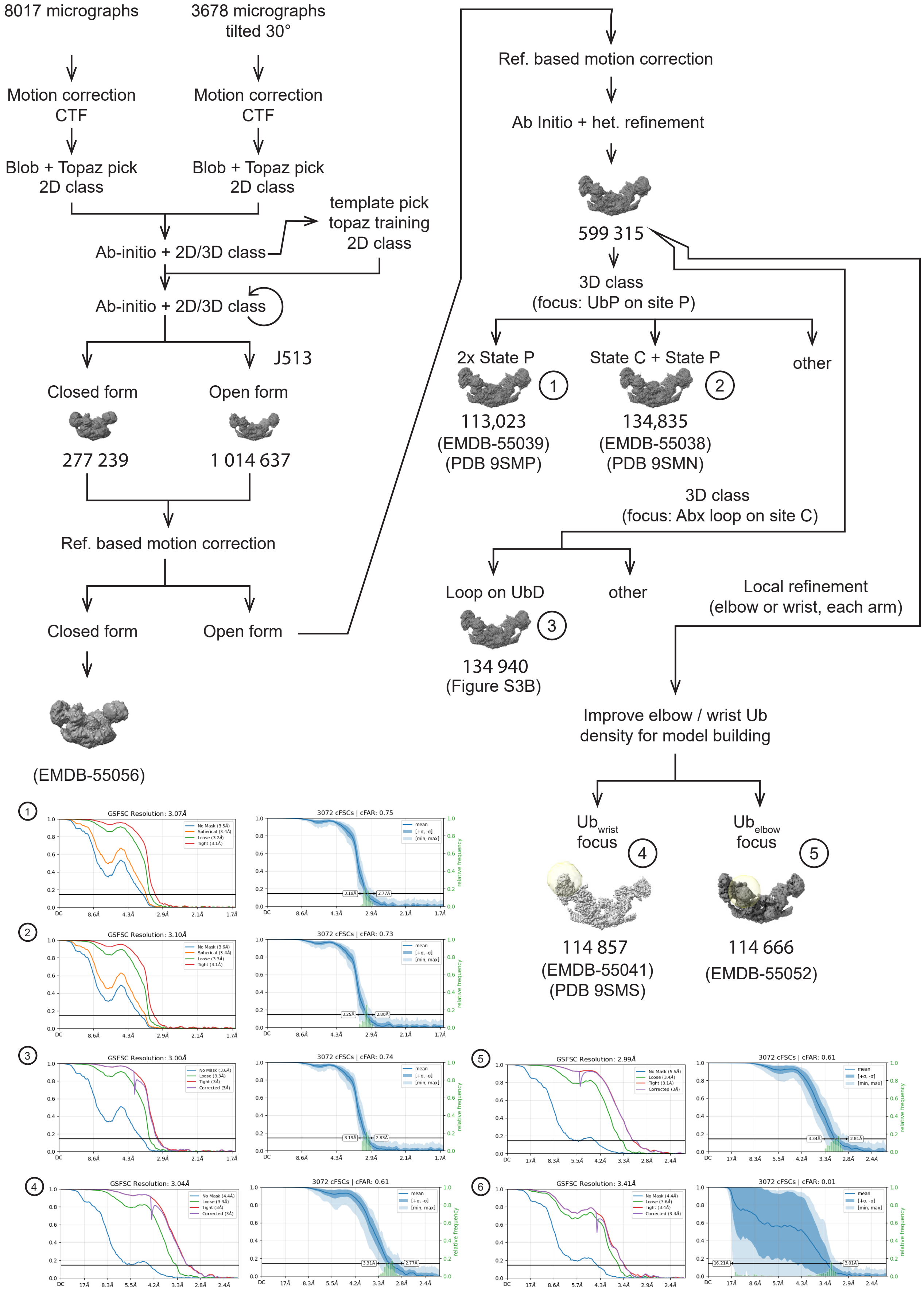

### Figure 1: EM processing workflow for the BRCA1-A: polyUb$_{ATA}$ dataset

The cryo-EM data processing workflow for the BRCA1-A complex in complex with polyUb$_{ATA}$ is shown. Two datasets (untilted and tilted by 30°) were collected from the same grid in separate sessions, using identical microscope settings. Maps and structures presented in the manuscript are highlighted, with particle counts and corresponding database accession numbers indicated. Map features (open/closed conformation and ligand-binding status) are summarised in Table 1 below. For each final map, the gold-standard Fourier shell correlation (GS-FSC) curve and the conical Fourier shell correlation (cFSC) plot are shown. cFSC plots were generated using the CryoSPARC orientation diagnostics tool[1].

Figure 2: 3D Reconstruction diagnostics for open BRCA1-A:polyUb$_{ATA}$ structure

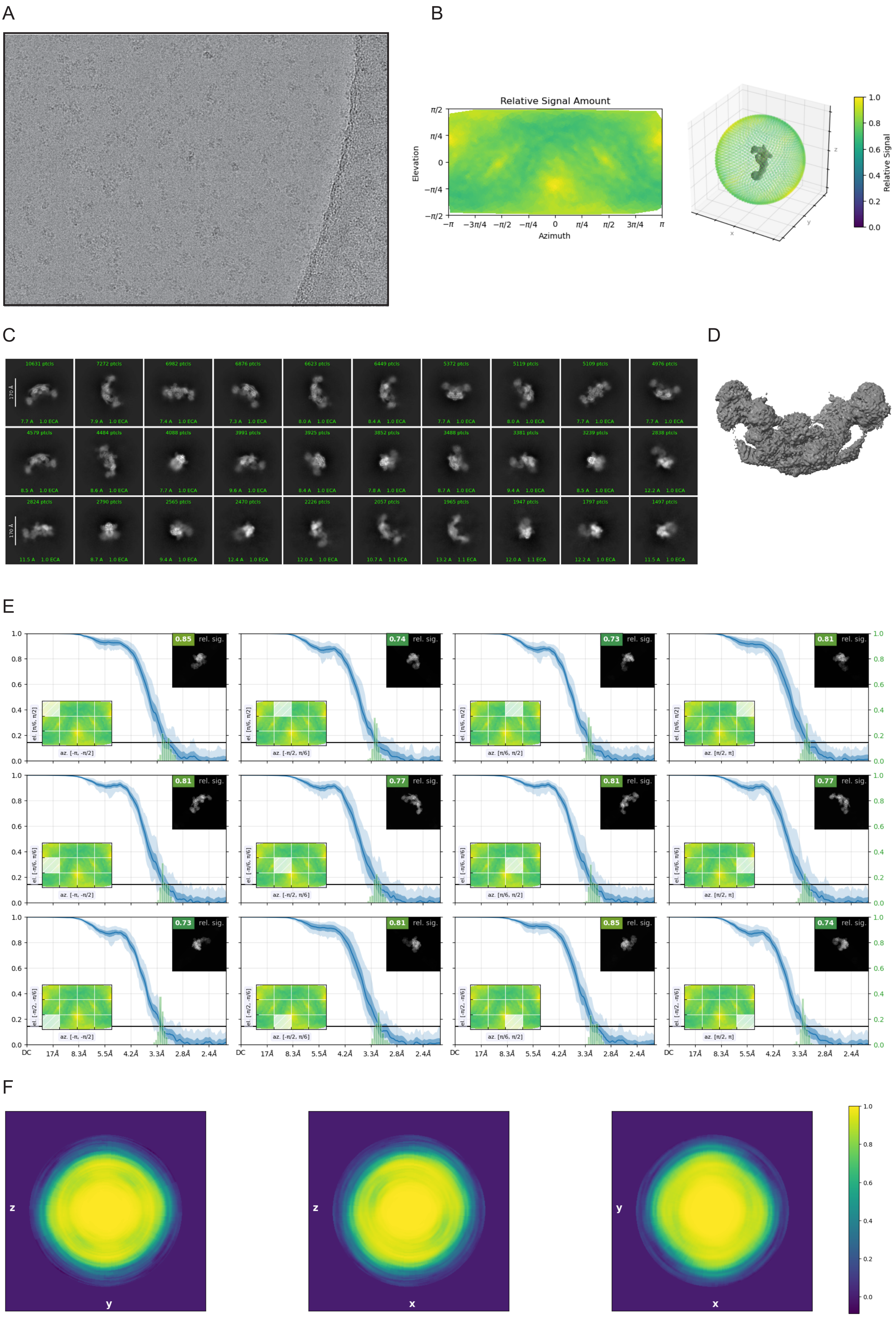

### Figure 2: Reconstruction diagnostics for open BRCA1-A:polyUb$_{ATA}$ 3D volume.

**(A)** Representative micrograph. **(B)** Relative signal amount vs. viewing direction. **(C)** 2D classification of the underlying particles. **(D)** Unfiltered reconstructed density **(E)** Average relative signal amount within azimuth-elevation viewing regions as computed by CryoSPARC orientation diagnostics tool[1]. Briefly, "the relative signal amount is computed by considering the FSC within a torus whose cross-sectional plane is orthogonal to a viewing vector. Concretely, relative signal is defined to be the weighted area under curve (AuC) of this toroidal FSC, normalized such that the maximum relative signal over the viewing sphere is 1"[1]. (F) 3DFSC volume central slices (clipped at 2.839 Å).

Figure 3: 3D Reconstruction diagnostics for closed BRCA1-A:polyUb$_{ATA}$ structure

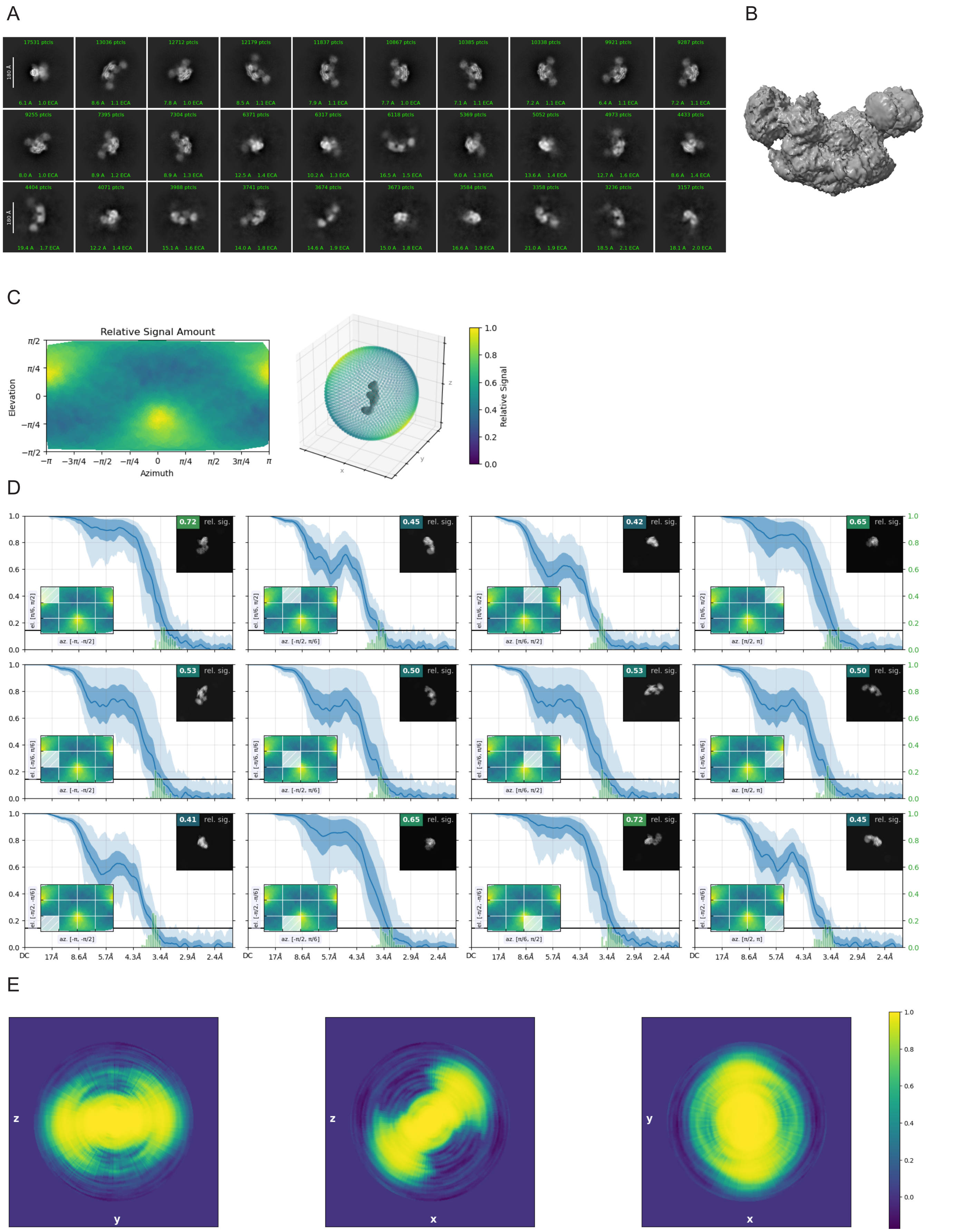


### Figure 3: Reconstruction diagnostics for closed BRCA1-A:polyUb$_{ATA}$ 3D volume.

**(A)** 2D classification of the underlying particles. **(B)** Unfiltered reconstructed density. **(C)** Relative signal amount vs. viewing direction. **(E)** Average relative signal amount within azimuth-elevation viewing regions as computed by CryoSPARC orientation diagnostics tool[1]. **(F)** 3DFSC volume central slices (clipped at 2.916 Å). The particles belong to the same dataset as the open BRCA1-A:polyUb$_{ATA}$ 3D volume.

Figure 4: BRCA1-A oligoUb$_{ATA}$ dataset processing workflow

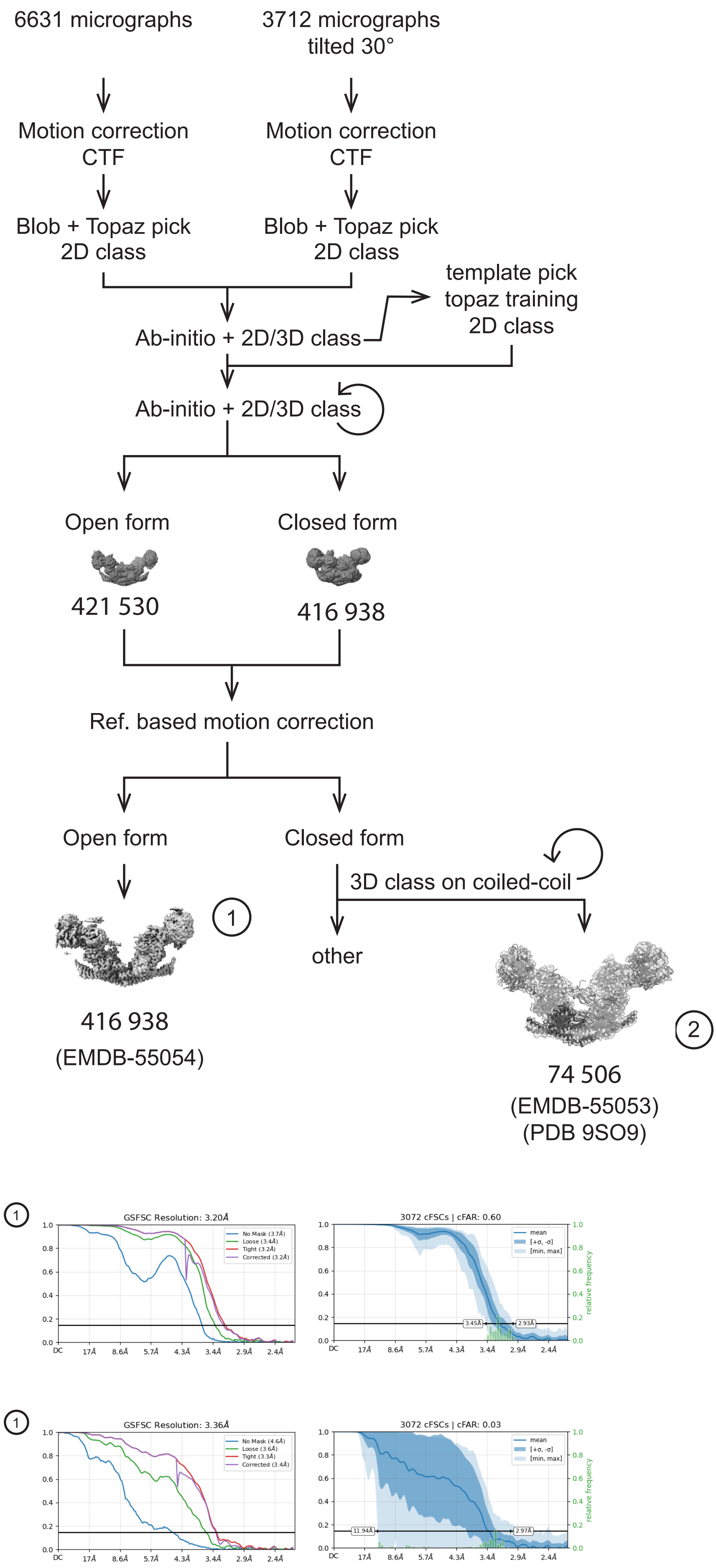


### Figure 4: EM processing workflow for the BRCA1-A: oligoUb$_{ATA}$ dataset

As in Figure 1, the cryo-EM processing workflow is shown for the BRCA1-A:oligoUb$_{ATA}$ dataset, with highlighted maps, particle counts, database accessions, and corresponding gold standard Fourier shell correlation (GS-FSC) and conical Fourier shell correlation (cFSC) plots. Two datasets (untilted and tilted by 30°) were collected from different grids prepared in the same freezing session. Map features are summarised in Table 1.

Figure 5: BRCA1-A $diUb_{ATA}$ dataset processing workflow

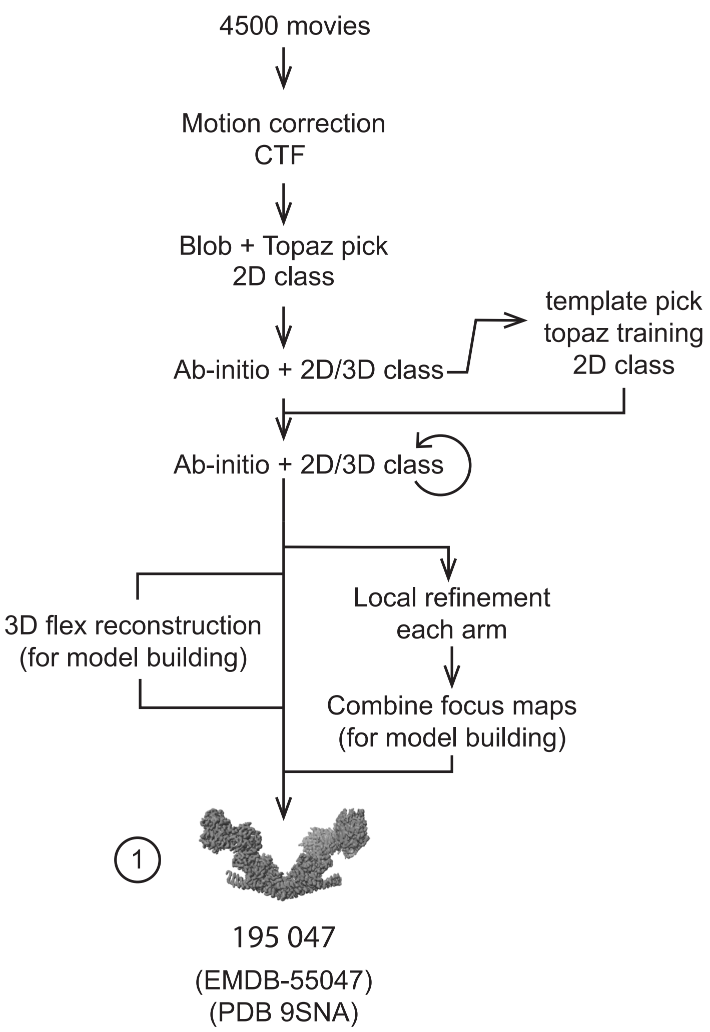


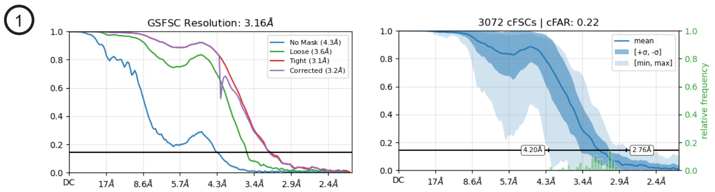


### Figure 5: EM processing workflow for the BRCA1-A: $diUb_{ATA}$ dataset

As in Figure 1, the cryo-EM processing workflow is shown for the BRCA1-A:$diUb_{ATA}$ dataset, with highlighted maps, particle counts, database accessions, and corresponding GS-FSC and cFSC plots. A closed form of this complex was not observed. Map features are summarised in Table 1.

Figure 6: BRCA1-A Apo dataset processing workflow

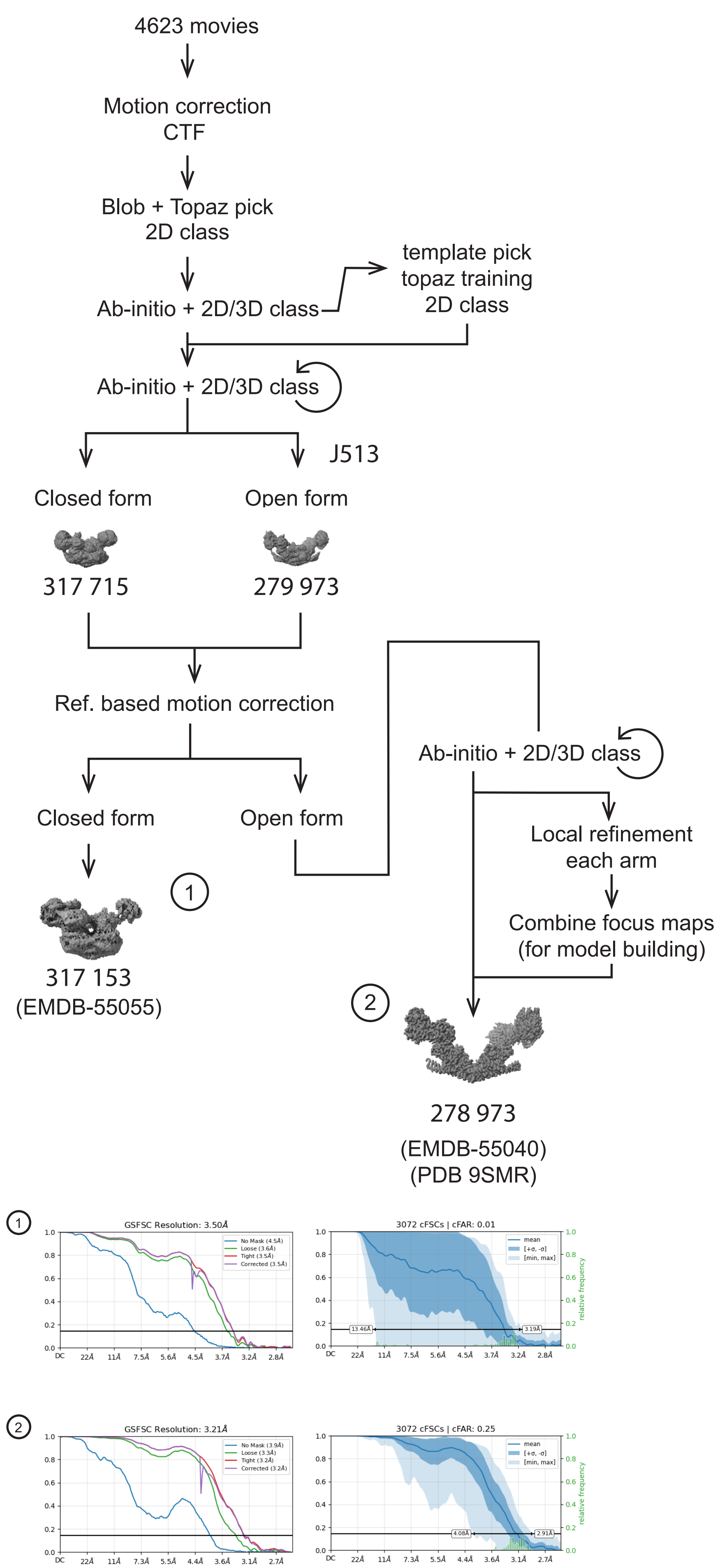


### Figure 6: EM processing workflow for the apo BRCA1-A dataset

As in Figure 1, the cryo-EM processing workflow is shown for the apo BRCA1-A dataset. In this dataset, oligoUb$_{ATA}$ was present in the sample, but no substrate was detected in the active site. Maps highlighted in the paper are shown, together with particle counts, database accessions, and corresponding GS-FSC and cFSC plots. Map features are summarised in Table
