## Supplementary code for "A ubiquitin chain-feeding mechanism for BRCA1-A": Figure_3D.ipynb

````
# %% [markdown]
## Note about this sandbox

This notebook implements an interactive pipeline that:

1. Subtracts imager-dependent background from scanned gels  
2. Allows selection of lanes and extraction of 1D lane profiles  
3. Normalizes signal intensity based on marker intensity  
4. Allows manual definition of integration gates on the 1D lane profiles (i.e. band selection)  
5. Plots the time-dependent intensity of ubiquitinated species  

The interactive components of the notebook have been disabled by setting `reprocess = False`.

Instead, checkpoints are provided containing:
- background-subtracted gels,
- selected lanes,
- and integration gates used for Figure 3D.

The same pipeline was used for all gel quantifications presented in the paper.

# %% [markdown]
## Environment Setup

This notebook is intended to be run in a Jupyter environment (e.g., Jupyter Notebook or JupyterLab).

### 1. Create a virtual environment (recommended)

Using `venv`:

```bash
python -m venv .venv
source .venv/bin/activate   # macOS / Linux
.venv\Scripts\activate      # Windows
```

Or using `conda`:

```bash
conda create -n myenv python=3.14
conda activate myenv
```

### 2. Install required dependencies

Install core libraries:

```bash
pip install numpy pandas matplotlib scipy pillow
```

Install Jupyter and widgets support:

```bash
pip install notebook jupyterlab ipywidgets ipympl
```

* If interactive plots do not render, ensure `ipympl` is installed.

# %% [code]
# imports
import pickle
import copy
from itertools import cycle
from functools import partial
from pathlib import Path

import matplotlib.pyplot as plt
import numpy as np
import pandas as pd

from matplotlib import patches
from matplotlib.colors import Colormap, to_hex
from matplotlib.gridspec import GridSpec
from PIL import Image, ImageOps
from scipy import sparse
from scipy.sparse.linalg import spsolve
from scipy.ndimage import gaussian_filter

# %% [code]
# Set the working directory containing the gel image series.
# Code Ocean capsule was set up with raw gel images are located in
# the "data/raw_tif" subdirectory of the capsule.
# Modify this path as needed.
# 
folder = Path.cwd().parent/"data"
assert folder.exists()

# Tif files exported by imageLab (Biorad Inc)
raw_file_folder = folder / "raw_tif"
pattern = "*.tif"
tif_files = sorted(raw_file_folder.glob(pattern))
assert tif_files
# Blank image obtained by imaging the empty tray
blank_image = raw_file_folder / "blank.tif"
assert blank_image.exists()

# remove blank image from list if present
tif_files = [p for p in tif_files if p != blank_image]
# keep only raw images - remove possible processing intermediates
image_paths = [
    p
    for p in tif_files
    if "output" not in p.name and "cropped" not in p.name and "subtracted" not in p.name
]
assert image_paths

# convert to grayscale, invert, and load as np.array-like matrix
raw_images = [
    np.array(ImageOps.invert(Image.open(p).convert("L"))) for p in image_paths
]
# same for blank image
blank = np.array(ImageOps.invert(Image.open(blank_image).convert("L")))

# %% [code]

# Explore how different background scaling factors (alpha) affect subtraction.
# The user should visually inspect the outputs and select the alpha that best
# removes background without suppressing real signal.

%matplotlib inline
plt.close("all")

def subtract_scaled_blank(raw, blank_field, alpha):
    raw = raw.astype(np.float32)
    corrected = raw - alpha * blank_field * np.mean(raw)
    return np.clip(corrected, 0, None)

alphas = np.linspace(1.0, 0.0, 10)
blank_smooth = gaussian_filter(blank.astype(np.float32), sigma=30)
blank_smooth /= np.mean(blank_smooth)

fig, axes = plt.subplots(len(raw_images), len(alphas) + 2, figsize=(40, len(raw_images)*5))
for ax_row, raw, path in zip(axes, raw_images, image_paths):
    ax_row[0].imshow(blank, cmap="Greys")
    ax_row[0].set_title("blank")
    ax_row[1].imshow(raw, cmap="Greys")
    ax_row[1].set_title(Path(path).name)
    for j, alpha in enumerate(alphas, start=2):
        corrected = subtract_scaled_blank(raw, blank_smooth, alpha)
        ax_row[j].imshow(corrected, cmap="magma")
        ax_row[j].set_title(f"α={alpha:.2f}")
        ax_row[j].axis("off")

analysis_folder = folder/"analysis"
analysis_folder.mkdir(exist_ok=True)
plt.savefig(analysis_folder /"background_subtraction_coefficients.png", format="png")
plt.tight_layout()
plt.show()

# %% [code]
# Select the desired alpha coefficient based on previous figure
# Verify that the chosen alpha removes background without introducing artifacts
# Background levels outside the gel should look uniformly grey.

alpha = 0.7


%matplotlib inline
plt.close("all")
blank_smooth = gaussian_filter(blank.astype(np.float32), sigma=30)
blank_smooth /= np.mean(blank_smooth)
blanked = [subtract_scaled_blank(img, blank_smooth, alpha) for img in raw_images]

fig, axes = plt.subplots(len(raw_images), 2, figsize=(10, len(raw_images)*5))
for i, ax in enumerate(axes):
    ax[0].imshow(raw_images[i], cmap="Greys")
    ax[0].set_title("Original")
    ax[0].axis("off")
    ax[1].imshow(blanked[i], cmap="Greys")
    ax[1].set_title(f"Background corrected (α={alpha})")
    ax[1].axis("off")
plt.tight_layout()
plt.savefig(analysis_folder / "background_correction.png", format="png")
plt.show()

# %% [code]
# Export background-subtracted images as [image_name]_subtracted.tif
background_sub_folder = analysis_folder / f"background_subtracted_gels"
background_sub_folder.mkdir(exist_ok=True)
for img, img_path in zip(blanked, image_paths):
    image = Image.fromarray(img).convert("L")
    image = ImageOps.invert(image)
    fileout = background_sub_folder / f"{img_path.name}_subtracted.tif"
    image.save(fileout, format="TIFF")

# %% [markdown]
Next step: straighten and crop the background-subtracted images in Photoshop or another suitable program.

Ensure all images have consistent width to height ratio and the same resolution (recommended: 600 × 500 pixels at 300 dpi).

Place the cropped files in the same folder as the background-subtracted images.

Append `_cropped` to each filename to distinguish them (e.g. `[image]_cropped.tif`).

# %% [code]
# cropped images are loaded from data/cropped_gels
images_by_longname = {}
cropped_gels_folder = folder/"cropped_gels"
for i in cropped_gels_folder.glob("*_cropped.tif"):
    imagename = i.name
    imagedata = ImageOps.invert(Image.open(i))
    images_by_longname[imagename] = imagedata
images_by_longname

# %% [code]
# Match sample IDs to the corresponding images.
# Ensure that the order of the sample IDs matches the order of the images listed above.
# Verify correctness using the output of of the cell.

sample_ids = [
    "deltains",
    "wt",
]

# map shorthand name to filename
image_name_mapping = {
    shortname: longname
    for shortname, longname in zip(sample_ids, images_by_longname.keys())
}
# index images by shortname
images = {
    shortname: image_data
    for shortname, image_data in zip(
        image_name_mapping.keys(), images_by_longname.values()
    )
}
# record the order in which images are displayed in the interactive plots.
image_index_mapping = {
    shortname: index for shortname, index in zip(images.keys(), range(len(images)))
}
index_to_name = {value: key for key, value in image_index_mapping.items()}

# prompt user to check that the files and the shortnames match
print("Please check correctness, or modify image shortname list above:")
for shortname, longname in image_name_mapping.items():
    print(f'"{shortname}" refers to original image "{longname}"')

# %% [code]
# set the desired width for lane extraction
WIDTH = 35

# %% [code]
# define functions to help with lane selection, plotting, etc.
def remove_all_axvlines(ax):
    for patch in ax.patches:
        if isinstance(patch, patches.Rectangle):
            patch.remove()


# function to drive vertical lines on ax at line_x_values
# uses rectangles to ensure that width scales properly
# since axvline's width is in points and does not
def redraw_all_lines(
    ax, line_x_values, color=None, cmap=plt.cm.tab10, clear_previous=True
):
    if clear_previous:
        remove_all_axvlines(ax)
    # input check
    if (color is None and cmap is None) or (color is not None and cmap is not None):
        raise ValueError("Provide either a single color or a cmap")

    for i, x_value in enumerate(line_x_values):
        if color is not None:
            facecolor = color
        else:  # we have a color map - we try to keep a direct correspondence between lane # and color
            if hasattr(cmap, "colors"):  # ListedColormap (tab10, tab20, sns palettes)
                palette = cmap.colors
                facecolor = palette[i % len(palette)]
            else:  # continuous colormap (viridis, magma, etc.)
                facecolor = cmap(i / max(len(line_x_values) - 1, 1))
        draw_lane(ax, x_value, facecolor)
    fig.canvas.draw()


def draw_lane(
    ax,
    x_value,
    facecolor=None,
    edgecolor=None,
    y_min=None,
    y_max=None,
    width=WIDTH,
    linewidth=0,
    alpha=0.5,
):
    if facecolor is None and edgecolor is None:
        raise ValueError("Please provide either facecolor or edgecolor")
    if facecolor is None:
        facecolor = "none"  # weird convention but whatever
    if edgecolor is None:
        edgecolor = "none"  # ditto
    if edgecolor != "none" and linewidth == 0:
        import logging

        logging.getLogger(__name__).warning(
            f"Attempting to set edgecolor {edgecolor} with linewidth == 0. Setting linewidth = 3 for visibility"
        )
        linewidth = 3
    mn, mx = ax.get_ylim()
    if y_min is None:
        y_min = mn
    if y_max is None:
        y_max = mx
    rect = patches.Rectangle(
        (x_value - width / 2, y_min),
        width,
        y_max - y_min,
        linewidth=linewidth,
        edgecolor=edgecolor,
        facecolor=facecolor,
        alpha=alpha,
    )
    ax.add_patch(rect)


def on_click(event):
    if event.xdata is None:
        return lanes
    clicked_ax = event.inaxes
    if clicked_ax is None:
        return lanes

    # Determine subplot index
    axes_list = list(fig.axes)
    ax_index = axes_list.index(clicked_ax)
    # Get stable key (image name)
    image_key = index_to_name[ax_index]
    ix = int(event.xdata)
    # Initialize list if not present
    if image_key not in lanes:
        lanes[image_key] = []
    if event.button == 2:  # middle click → delete closest
        if lanes[image_key]:
            closest_value = min(lanes[image_key], key=lambda x: abs(x - ix))
            lanes[image_key].remove(closest_value)
    elif event.button == 1:  # left click → add
        lanes[image_key].append(ix)
    # Redraw
    remove_all_axvlines(clicked_ax)
    redraw_all_lines(clicked_ax, lanes[image_key])
    return lanes


def get_palettes(number_of_palettes):
    palettes = cycle([plt.cm.Blues, plt.cm.Reds, plt.cm.Greens, plt.cm.Greys])
    return [next(palettes) for i in range(number_of_palettes)]


def get_gradient_colors(
    cmap: Colormap, number: int, darker: bool = True, inverted: bool = True
):
    # darker: lightest color of the gradient is a bit darker than default
    if darker:
        start = 50
    else:
        start = 0
    gradient_positions = np.round(np.linspace(start, 255, number)).astype(int)
    if inverted:
        gradient_positions = gradient_positions[::-1]
    return [cmap(i) for i in gradient_positions]


lane_select_instructions = "Left click on images to select lanes. Middle click removes lane. \n Drag triangle in lower right corner to increaase image size"

# %% [code]
# This cell normally supports interactive gel lane annotation.
# In this capsule, interactivity is disabled and lanes are loaded from saved checkpoints.
# Interactive code is retained for reference.

# Interactives: mark lanes for analysis. Include marker and loading control.
# Resize the interactive figure as needed before clicking.
# Include the enzyme lane and one marker lane.
# Ensure all gels have the same number of selected lanes.

# Set `reprocess = True` to redo lane selection.
# Otherwise, previously picked lanes will be loaded in the next cell.

reprocess_lanes = False

plt.close("all")
if reprocess_lanes:
    %matplotlib ipympl
    lanes = {}
    fig, axes = plt.subplots(1, len(images))
    fig.canvas.header_visible = False #type:ignore
    fig.canvas.footer_visible = False #type:ignore
    fig.canvas.resizable = True #type:ignore
    fig.canvas.toolbar_visible = False #type:ignore
    if len(images) == 1: #nnot subscriptable
        axes.imshow(img, cmap="Greys")
    else:
        for index, data in enumerate(images.items()):
            imgname, img = data
            axes[index].imshow(img, cmap="Greys")
            axes[index].set_title(f"{imgname}")
    if reprocess_lanes:
        # Connect click events to the function
        fig.canvas.mpl_connect("button_press_event", on_click)
        # Display instructions for lane selection on on image
        fig.text(
            0.5, 0.97, lane_select_instructions,
            ha="center",
            va="top",
            fontsize=11,
            bbox=dict(boxstyle="round", facecolor="white", alpha=0.8)
        )
    figure_axes = plt.gcf().axes
    plt.show()

# %% [code]
# save selected lanes 
# or load previously selected lanes from checkpoint
checkpoint_file = folder / "checkpoints/lanes.pkl"

plt.close("all")
if reprocess_lanes:
    for key, lines in lanes.items():
        # sort the lanes left to right
        lanes[key] = sorted(lines)
    # save the selected lanes
    # so that we can reproduce this flow
    with open(checkpoint_file, "wb") as f:
        pickle.dump(lanes, f)
else:
    with open(checkpoint_file, "rb") as f:
        lanes = pickle.load(f)

ordered = False  # guard flag to avoid repeat execution of cell below

# %% [code]
# Lane annotation. This is done manually.

# are time points t(start) -> t(end) in left to right order (True) or mirrored (False)
natural_order = False

#Define which lanes correspond to the marker and loading control on each gel.
#Lanes are indexed starting from 0 (leftmost lane = 0), with each gel indexed independently.

marker_index = {
    "deltains": 0,
    "wt": -1,
}
# note which selected lanes are enzyme loading control in each gel
enzyme_index = {
    "deltains": 1,
    "wt": 0,
}

# specify labels for time points
# 1024 ~= 17.1 hrs ~= O/N
time_points_labels = [
    0,
    0.5,
    1,
    2,
    4,
    8,
    16,
    32,
    64,
    128,
    1024,
]

if not natural_order:
    time_points_labels = time_points_labels[::-1]
x_positions = lanes

# sort lanes by type and order
marker_x = {}
load_control_x = {}
time_points_x = {}
if not ordered:  # guard clause to prevent data corruption in case of repeated execution
    for image, lane_x_positions in lanes.items():
        marker_x[image] = lane_x_positions[marker_index[image]]
        load_control_x[image] = lane_x_positions[enzyme_index[image]]
        lane_x_positions.remove(marker_x[image])
        lane_x_positions.remove(load_control_x[image])
        # remove marker and enzyme lanes from time points ensuring order
        time_points_x[image] = lane_x_positions
        number_of_lanes = len(lane_x_positions)
        number_of_time_points = len(time_points_labels)
        assert (
            number_of_lanes == number_of_time_points
        ), f"Image {image} has {number_of_lanes} lanes but {number_of_time_points} timepoints are specified"
    ordered = True

# %% [code]
# check that time series are correctly assigned
# by plotting the lanes in color gradient (light->dark = earlier to later time points)

palettes = {
    sample: palette for sample, palette in zip(images.keys(), get_palettes(len(images)))
}
gradients = {
    sample: get_gradient_colors(
        palettes[sample], len(time_points_labels), darker=True, inverted=natural_order
    )
    for sample in images.keys()
}

%matplotlib inline
plt.close("all")
fig2, axes = plt.subplots(1, len(images), figsize=(20, 6))
if len(images) == 1:
    axes = [axes]  # ensure this is iterable if there is only a single plot
for index, ax in enumerate(axes):
    ax.imshow(images[index_to_name[index]], cmap="Greys")
for sample, lines_x in time_points_x.items():
    cmap = palettes[sample]
    colors = iter(gradients[sample])
    img_index = image_index_mapping[sample]  # which image index is the sample on
    for x_value in lines_x:
        color = next(colors)
        draw_lane(axes[img_index], x_value, facecolor=color)
    draw_lane(axes[img_index], load_control_x[sample], facecolor="yellow")
    draw_lane(axes[img_index], marker_x[sample], facecolor="grey")
    axes[img_index].set_title(f"{index_to_name[img_index]}")
#add user instructions
info = "Check that every lane is highlighted, and that lanes follow the color gradient light->dark (early->late)"
info += "\nMarkers are grey, loading controls are yellow"
fig2.text(
    0.5, 1, info,
    ha="center",
    va="top",
    fontsize=11,
    bbox=dict(boxstyle="round", facecolor="white", alpha=0.8)
)
plt.savefig(analysis_folder/"selected_lanes.png")
plt.show()

# %% [code]
# For each user-defined lane position we extract a vertical stripe centered on that x-coordinate.
# The stripe has width WIDTH and spans the full height of the gel (optionally cropped at the top
# or bottom using min_x / max_x).

# For every row in this stripe we compute the mean pixel intensity across the stripe width.
# This produces a 1-D vertical intensity profile for the lane.

# Rationale:
# 1. Gel bands are approximately rectangular. Their intensity is therefore nearly constant
#    across the horizontal axis of the band. Averaging across a stripe gives a good proxy
#    for the total band signal while being insensitive to the exact lane boundaries.
# 2. Minor horizontal bleed from neighbouring lanes affects only a few pixels and is
#    largely suppressed by averaging across the stripe width.


def get_profile(image_array, x, width=WIDTH, min_x=None, max_x=None):
    # gets a column stripe of width=width from an image array
    # then averages row-wise to return a single average column
    # minx_x and max_x are the percentage of x-axis to cut from each end
    if width % 2:  # odd
        width += 1
    if min_x is not None and max_x is not None:
        total = image_array.shape[0]
        left_x = int(total * min_x)
        right_x = int(total * max_x)
    else:
        left_x = 0
        right_x = image_array.shape[0]
    halfwidth = width // 2
    stripe = image_array[
        left_x:right_x, x - halfwidth : x + halfwidth
    ]  # Extract width-wide stripe centered at x
    averaged_column = stripe.mean(axis=1)  # average over row
    return averaged_column


def plot_curve_series(series, ax, color_generator):
    # A bit of a hack, but I mixed my data structures downstream and I do not want
    # to refactor everything.
    # series might be a List[np.array] or a Dict[str:np.array]
    if isinstance(series, dict):
        iterator = series.values()
    elif isinstance(series, list):
        iterator = series
    else:
        raise ValueError("Expecting a list or a dict")
    for curve in iterator:
        ax.plot(curve, color=next(color_generator))


# calculate time point curves
time_curves = {key: [] for key in images.keys()}
for sample, img in images.items():
    for x in lanes[sample]:
        time_curves[sample].append(
            get_profile(
                np.array(img),
                x,
                width=WIDTH,
            )
        )
# calculate marker curves
marker_curves = {}
for sample, img in images.items():
    marker_curves[sample] = get_profile(
        np.array(img),
        marker_x[sample],
        width=WIDTH,
    )
# calculate loading control curves
loading_curves = {}
for sample, img in images.items():
    loading_curves[sample] = get_profile(
        np.array(img), load_control_x[sample], width=WIDTH
    )

fig, axes = plt.subplots(len(images) + 2, figsize=(20, 10))
index = 0
for ax, sample in zip(axes, images.keys()):  # +2 ignored via zip
    ax.set_title(sample)
    cmap = palettes[sample]
    colors = iter(gradients[sample])
    plot_curve_series(time_curves[sample], ax, colors)
    index += 1
# plot marker_x
for index, sample in enumerate(images.keys()):
    cmap = plt.cm.tab10  # discrete colors
    # marker_x
    axes[-2].plot(marker_curves[sample], color=cmap(index))
    axes[-1].plot(loading_curves[sample], color=cmap(index))
axes[-2].set_title("marker_x")
axes[-1].set_title("loading")
plt.tight_layout()
plt.show()

# %% [code]
# The spike observed at the end of the curves is an imaging artifact
# caused by the raised lip of the precast gels used in the experiment,
# which consistently appears as a very dark region in the image.
# This region is trimmed prior to analysis because it can interfere
# with baseline correction procedures.
TRIM_RIGHT = 25
fig3, axes3 = plt.subplots(3, figsize=(20, 10))
trimmed_marker_curves = copy.deepcopy(marker_curves)
trimmed_loading_curves = copy.deepcopy(loading_curves)
trimmed_time_curves = copy.deepcopy(time_curves)
for index, sample in enumerate(images.keys()):
    trimmed_marker_curves[sample] = marker_curves[sample][:-TRIM_RIGHT]
    trimmed_loading_curves[sample] = loading_curves[sample][:-TRIM_RIGHT]
    for index, curve in enumerate(time_curves[sample]):
        trimmed_time_curves[sample][index] = curve[:-TRIM_RIGHT]
    axes3[0].plot(trimmed_marker_curves[sample])
    axes3[0].set_title("marker lanes")
    axes3[1].plot(trimmed_loading_curves[sample])
    axes3[1].set_title("loading controls")
    colors = iter(gradients[sample])
    plot_curve_series(trimmed_time_curves[sample], axes3[2], colors)
    axes3[2].set_title("All time points")

# %% [code]
# Apply a simple vertical shift to each intensity profile for ease of visualisation
# We estimate the baseline by averaging the last ~500 data points of the curve
# (a region assumed to contain no bands). This value is subtracted from the
# entire profile so that the tail of the curve is approximately zero.
#
# This step compensates for small lane-to-lane differences in baseline image
# intensity. A more rigorous baseline correction will be performed later.

fig5, axes5 = plt.subplots(len(images) + 2, figsize=(20, 10))
shifted_time_curves = {}
shifted_marker_curves = {}
shifted_loading_curves = {}
for index, sample in enumerate(images.keys()):
    shifted_time_curves[sample] = {}
    for time_point, curve in zip(time_points_labels, trimmed_time_curves[sample]):
        shift = np.mean(curve[-500:])
        shifted = curve - shift
        shifted_time_curves[sample][time_point] = shifted
    shifted_marker = np.mean(marker_curves[sample][-500:])
    shifted_marker_curves[sample] = trimmed_marker_curves[sample] - shifted_marker
    shifted_load = np.mean(loading_curves[sample][-500:])
    shifted_loading_curves[sample] = trimmed_loading_curves[sample] - shifted_load
    axes5[index].set_title(f"{sample} shifted")
    plot_curve_series(
        shifted_time_curves[sample],
        axes5[index],
        color_generator=iter(gradients[sample]),
    )
    color = palettes[sample](128)
    axes5[-2].plot(shifted_marker_curves[sample], color=color)
    axes5[-2].set_title("markers shifted")
    axes5[-1].plot(shifted_loading_curves[sample], color=color)
    axes5[-1].set_title("loading shifted")


plt.tight_layout()
plt.show()

# %% [code]
# check that lanes match profile curves (sample 1) and save figures for all gels.

%matplotlib inline
plt.close("all")

for i, sample in enumerate(images.keys()):
    rows = len(time_points_x[sample]) + 2  # marker, loading

    fig = plt.figure(figsize=(30, 16))
    gs = GridSpec(rows, 2, figure=fig, width_ratios=[1, 2])

    colors = gradients[sample]

    # left column: timepoint curves
    for index, time_point in zip(range(rows), time_points_labels):
        ax = fig.add_subplot(gs[index, 0])
        ax.plot(
            shifted_time_curves[sample][time_point],
            label=f"{time_points_labels[index]} minutes",
            color=to_hex(colors[index]),
        )
        ax.legend()

    # marker curve
    ax = fig.add_subplot(gs[-2, 0])
    ax.plot(trimmed_marker_curves[sample], label="marker", color="yellow")
    ax.legend()

    # loading control curve
    ax = fig.add_subplot(gs[-1, 0])
    ax.plot(trimmed_loading_curves[sample], label="enzyme", color="grey")
    ax.legend()

    # right column: gel image
    ax_image = fig.add_subplot(gs[:, 1])
    ax_image.imshow(images[sample], cmap="gray_r")

    # timepoint lanes
    for index, x in enumerate(x_positions[sample]):
        draw_lane(
            ax_image,
            x,
            facecolor=None,
            edgecolor=to_hex(colors[index]),
            linewidth=3,
        )

    # marker lane
    draw_lane(
        ax_image,
        marker_x[sample],
        facecolor=None,
        edgecolor="orange",
        linewidth=3,
    )

    # loading control lane
    draw_lane(
        ax_image,
        load_control_x[sample],
        facecolor=None,
        edgecolor="black",
        linewidth=3,
        width=WIDTH
    )

    ax_image.legend(labels=time_points_labels + ["marker", "enzyme"])

    fig.savefig(analysis_folder / f"lanes_vs_curves_comparison_{sample}.png", dpi=300, bbox_inches="tight")

# %% [code]
# Define baseline correction, peak detection and peak integration functions

# Prior to peak integration we perform baseline correction.
#
# We use the Asymmetric Least Squares (AsLS) approach described in:
#
# @techreport{Eilers2005AsLS,
#   author  = {Eilers, Paul H. C. and Boelens, Hans F. M.},
#   title   = {Baseline Correction with Asymmetric Least Squares Smoothing},
#   institution = {Leiden University Medical Centre},
#   year    = {2005},
#   month   = {October}
# }
#
# AsLS methods are widely used to remove slowly varying baselines from
# chromatographic and spectroscopic signals.
#
# More sophisticated baseline algorithms exist (see e.g.
# https://pybaselines.readthedocs.io/en/latest/algorithms_2d/whittaker_2d.html),
# but AsLS provides a simple and robust solution for our use case.
#
# The implementation below is a minimal reimplementation to avoid adding a
# dependency on the pybaselines library.


def baseline_asls(y, lam=1e6, p=0.01, niter=10):
    """
    Asymmetric least squares baseline.

    lam: smoothness parameter (larger -> smoother baseline)
    p: asymmetry parameter (smaller -> baseline stays below peaks)
    """
    y = np.asarray(y, dtype=float)
    n = y.size

    D = sparse.diags([1.0, -2.0, 1.0], [0, 1, 2], shape=(n - 2, n), dtype=float)
    w = np.ones(n)
    dtd = (D.T @ D).tocsc()

    for _ in range(niter):
        W = sparse.diags(w, 0, shape=(n, n))
        Z = (W + lam * dtd).tocsc()
        z = spsolve(Z, w * y)
        w = p * (y > z) + (1 - p) * (y <= z)

    return z


# peak detection and integration use scipy

from scipy import signal


def detect_peaks(curve, height=None, prominence=None, distance=None, width=None):
    peaks, props = signal.find_peaks(
        curve, height=height, prominence=prominence, distance=distance, width=width
    )
    return peaks, props


def integrate_peaks(curve, peaks, left_bases, right_bases):
    integrals = {}
    for i, p in enumerate(peaks):
        left = int(left_bases[i])
        right = int(right_bases[i])
        integrals[p] = np.trapezoid(curve[left : right + 1])
    return integrals
# baseline correct and integrate marker lanes for normalization


def baseline_correct(curve, lam=1e6, p=0.01, niter=10):
    baseline = baseline_asls(curve, lam=lam, p=p, niter=niter)
    corrected = curve - baseline
    return corrected, baseline


def integrate_region(curve, l, r):
    x = np.arange(l, r + 1)
    return np.trapezoid(curve[l : r + 1], x)


def detect_marker_peaks(marker_curves, prominence=5, height=2, baseline_kwargs=None):
    if baseline_kwargs is None:
        baseline_kwargs = dict(lam=1e6, p=0.01, niter=10)

    results = {}
    for sample, curve in marker_curves.items():
        corrected, baseline = baseline_correct(curve, **baseline_kwargs)

        peaks, props = detect_peaks(corrected, height=height, prominence=prominence)

        peak_data = []
        for k, p in enumerate(peaks):
            l = int(props["left_bases"][k])
            r = int(props["right_bases"][k])
            integral = integrate_region(corrected, l, r)
            peak_data.append(dict(peak=p, left=l, right=r, integral=integral))

        results[sample] = dict(
            baseline=baseline,
            corrected=corrected,
            peaks=peaks,
            properties=props,
            peak_data=peak_data,
        )

    return results


def plot_marker_curves(marker_results, palettes):
    n = len(marker_results)
    fig, axes = plt.subplots(1, n, figsize=(4 * n, 4), sharey=True)
    if n == 1:
        axes = [axes]

    for ax, (sample, res) in zip(axes, marker_results.items()):
        y = res["corrected"]
        baseline = res["baseline"]
        cmap = palettes[sample]
        color = cmap(0.6)
        x = np.arange(len(y))

        ax.plot(y, color=color)
        ax.plot(baseline, linestyle="--", color="grey", linewidth=1)
        ax.axhline(0, linestyle="--", color="grey", linewidth=1)

        for k, d in enumerate(res["peak_data"]):
            p, l, r = d["peak"], d["left"], d["right"]
            xr = np.arange(l, r + 1)

            ax.scatter(p, y[p], color="red", s=25)
            ax.text(p, y[p], str(k), fontsize=9, ha="center", va="bottom")
            ax.fill_between(xr, 0, y[l : r + 1], color=color, alpha=0.35)

        ax.set_title(sample)

    fig.supxlabel("index")
    fig.supylabel("signal")
    fig.tight_layout()
    return fig


# %% [code]
# Correct staining intensity by normalizing each sample to the molecular weight marker intensity.
# The PageRuler unstained protein ladder (Thermo Fisher) was used here, though any suitable marker can be used.
# Four microliters of marker were loaded per gel.

# Integrate marker peaks and inspect results
marker_results = detect_marker_peaks(marker_curves, prominence=3, height=1)
fig = plot_marker_curves(marker_results, palettes)
plt.savefig(analysis_folder / "marker_intensity_comparison.tif", format="TIFF")
plt.show()

# %% [code]
# Calculate normalization factors using the intensities of the 25, 20, and 15 kDa marker bands.
# Peak integration may occasionally detect ghost peaks.
# Use the image above to identify the peaks corresponding to the marker bands in each sample.

peak_ids = {"wt": [9, 10, 11],
            "deltains":[9, 10, 11]}
band_names = [25, 20, 15]
mean_intensities = {}
for sample in marker_results.keys():
    mean_intensities[sample] = 0
    # we average over the intensity of the three bands
    for index, id_ in enumerate(peak_ids[sample]):
        mean_intensities[sample] += marker_results[sample]["peak_data"][id_]["integral"]
    mean_intensities[sample] = np.round(mean_intensities[sample] / 3, 3)
#find which gel has the highest intensity of staining
max_value = max(mean_intensities.values())
#calculate intensity ratios
norm_factor = {}
for sample, value in mean_intensities.items():
    norm_factor[sample] = round(max_value / value, 3)

# apply normalisation factor to every sample
normalized_curves = {}
for sample, curves in shifted_time_curves.items():
    normalized_curves[sample] = {}
    for tp, data in curves.items():
        normalized_curves[sample][tp] = data * norm_factor[sample]

print(f"Mean intensities: {mean_intensities}")
print(f"Normalization factor: {norm_factor}")

# %% [code]
# inspect effect of lambda and p values of AsLS on sample baseline correction
# reordering samples by substrate for plotting convenience
samples = list(image_name_mapping.keys())

plt.close("all")

lams = [1e04, 1e05, 1e06, 1e07, 1e08]
niter = 10

n_tp = len(time_points_labels)
n_lam = len(lams)

fig, axes = plt.subplots(n_tp, n_lam, figsize=(4 * n_lam, 3 * n_tp), sharex=True)

if n_tp == 1:
    axes = np.array([axes])

for i, tp in enumerate(time_points_labels):
    for j, lam in enumerate(lams):
        ax = axes[i, j]
        y = normalized_curves[samples[0]][tp]  # choose sample to inspect
        ax.plot(y, alpha=0.35, label="raw")
        for p, color in zip([0.1, 0.01, 0.001], ["red", "green", "blue"]):
            b = baseline_asls(y, lam=lam, p=p, niter=niter)
            y_corr = y - b
            ax.plot(y_corr, label=f"corrected_P={p}", color=color)
            ax.axhline(0, linestyle="dotted", linewidth=1)
            if i == 0:
                ax.set_title(f"λ = {lam:.1e}")
            if j == 0:
                ax.set_ylabel(tp)
            if i == 0 and j == 0:
                ax.legend()

    fig.supxlabel("index")
    fig.supylabel("signal")
    fig.tight_layout()
    plt.savefig(analysis_folder / f"{sample}_baseline_parameter_estimation.tif", format="TIFF")


# %% [code]
# choose appropriate lambda and p
lam = 1e07
p_asls = 0.01

baselines = {}
baselined_curves = {}
# reordering samples for convenience

for sample in samples:
    baselines[sample] = {}
    baselined_curves[sample] = {}
    curves = normalized_curves[sample]
    n = len(curves)
    ncols = 4
    nrows = int(np.ceil(n / ncols))
    fig, axes = plt.subplots(
        nrows, ncols, figsize=(4 * ncols, 3 * nrows), sharex=True, sharey=True
    )
    axes = axes.flatten()
    for i, (tp, data) in enumerate(curves.items()):
        baseline = baseline_asls(data, lam=lam, p=p_asls)
        baselines[sample][tp] = baseline
        baselined_curves[sample][tp] = data - baseline
        ax = axes[i]
        ax.plot(data, label="curve")
        ax.plot(baseline, label="baseline")
        ax.plot(baselined_curves[sample][tp], label="subtracted", color="red")
        ax.axhline(0, linestyle="--", color="grey", linewidth=1)
        ax.set_title(str(tp))
    for j in range(i + 1, len(axes)):
        fig.delaxes(axes[j])
    fig.suptitle(sample)
    handles, labels = ax.get_legend_handles_labels()
    fig.legend(handles, labels, loc="upper right")
    plt.savefig(analysis_folder / f"{sample}_baseilne_correction.tif", format="TIFF")
    plt.tight_layout()
    plt.show()

# %% [code]
# extract lanes for plotting
def extract_lane(image, x, width=WIDTH, top=None, bottom=None):
    img = np.asarray(image)
    if not width % 2:
        width -= 1
    if img.ndim == 3:
        raise ValueError(f"Expected 2D matrix: {img.ndim}D matrix found instead.")
    h, w = img.shape
    l = max(0, int(x - width // 2))
    r = min(w, int(x + width // 2))
    stripe = img[:, l:r]
    if top is not None or bottom is not None:
        t = 0 if top is None else top
        b = h if bottom is None else bottom
        stripe = stripe[t:b, :TRIM_RIGHT]
    return stripe


extracted_lanes = {}

for sample, img in images.items():
    extracted_lanes[sample] = {}
    for tp, x in zip(time_points_labels, time_points_x[sample]):
        stripe = extract_lane(img, x)
        extracted_lanes[sample][tp] = stripe

# %% [code]
# define some functions for interactive gate selection and plotting
# # relies on global objects extracted_lanes, baselined_curves, gradients


class AxData:
    def __init__(self, sample, time_point, index, axis_type):
        self.sample = sample
        self.time_point = time_point
        self.index = index
        self.axis_type = axis_type


def plot_sample_set(samples, dpi=100, narrow=False, supernarrow=False):
    lanes0 = extracted_lanes[samples[0]]
    tps = list(lanes0.keys())

    n_tp = len(tps)
    n_samples = len(samples)

    h, w = next(iter(lanes0.values())).T.shape

    if narrow:
        x_factor = 1
    else:
        x_factor = 2
    if supernarrow:
        x_factor = 0.25
    fig, axs = plt.subplots(
        2 * n_tp,
        n_samples,
        figsize=(x_factor * n_samples * w / dpi, 4 * n_tp * h / dpi),
        dpi=dpi,
        gridspec_kw={"height_ratios": [2, 1] * n_tp},
    )

    ax_table = {}

    for j, sample in enumerate(samples):
        lanes = extracted_lanes[sample]
        curves = baselined_curves[sample]

        for i, tp in enumerate(tps):
            ax_curve = axs[2 * i, j]
            ax_img = axs[2 * i + 1, j]

            ax_table[ax_curve] = AxData(sample, tp, i, "curve")
            ax_table[ax_img] = AxData(sample, tp, i, "image")

            stripe = lanes[tp]
            profile = curves[tp]
            L = len(profile)
            stripe = stripe[:L, :]
            x = np.arange(L)

            ax_curve.plot(x, profile, lw=3, color=gradients[sample][-5])
            ax_curve.set_xlim(0, L)
            ax_curve.set_xticks([])
            ax_curve.set_yticks([])

            if i == 0:
                ax_curve.set_title(sample)
            if j == 0:
                ax_curve.set_title(tp, loc="left")

            ax_img.imshow(
                1 - stripe.T,
                cmap="gray",
                origin="lower",
                interpolation="none",
                aspect="auto",
            )
            ax_img.set_xticks([])
            ax_img.set_yticks([])

    plt.tight_layout()
    return fig, axs, ax_table


# function to help with gate plotting
# gates: Dict[sample, Dict[time_point, List[Gate]]]


class Gate:
    def __init__(self, x: int, width: int = WIDTH):
        self.x = x
        self.width = width

        # per-axes artist storage
        # {ax: {"center": Line2D, "left": Line2D, "right": Line2D}}
        self._artists = {}

        # stable styling for this gate instance
        self._style = {"c1": None, "c2": None}

    def __repr__(self):
        return f"Gate at x: {self.x}, width = {self.width}"


def on_click(event, ax_table, gates):
    if event.inaxes not in ax_table or event.xdata is None:
        return

    x = int(event.xdata)
    if event.button == 1:
        add_gate(event.inaxes, ax_table, gates, x)
    elif event.button == 3:
        delete_gate(event.inaxes, ax_table, gates, x)


def on_key(event, ax_table, gates):
    if event.inaxes not in ax_table or event.xdata is None:
        return

    x = int(event.xdata)

    if event.key == "left":
        move_gate(event.inaxes, ax_table, gates, x, -2)
    elif event.key == "right":
        move_gate(event.inaxes, ax_table, gates, x, 2)
    elif event.key == "+":
        adjust_gate_width(event.inaxes, ax_table, gates, x, 10)
    elif event.key == "-":
        adjust_gate_width(event.inaxes, ax_table, gates, x, -10)


def on_scroll(event, ax_table, gates):
    if "shift" not in event.modifiers:
        return
    if event.inaxes not in ax_table or event.xdata is None:
        return

    x = int(event.xdata)
    adjust_gate_width(event.inaxes, ax_table, gates, x, event.step * 4)


def find_nearest_gate(gates, sample, tp, x):
    lst = gates[sample][tp]
    if not lst:
        return None
    return min(range(len(lst)), key=lambda i: abs(lst[i].x - x))


cycle_colors = cycle(plt.cm.tab10.colors)


def _gate_positions(gate):
    x = gate.x
    w = gate.width
    return x, x - w / 2, x + w / 2


def _draw_gate_on_axis(ax, gate, c1, c2):
    x, l, r = _gate_positions(gate)

    center = ax.axvline(x, color=c1, lw=1)
    center._is_gate = True

    left = ax.axvline(l, color=c2, lw=1, ls="--")
    left._is_gate = True

    right = ax.axvline(r, color=c2, lw=1, ls="--")
    right._is_gate = True

    gate._artists[ax] = {
        "center": center,
        "left": left,
        "right": right,
    }


def _update_gate_artists(gate):
    x, l, r = _gate_positions(gate)

    touched_figures = set()

    for ax, artists in gate._artists.items():
        artists["center"].set_xdata([x, x])
        artists["left"].set_xdata([l, l])
        artists["right"].set_xdata([r, r])
        touched_figures.add(ax.figure)

    for fig in touched_figures:
        if fig.canvas is not None:
            fig.canvas.draw_idle()


def _remove_gate_artists(gate):
    touched_figures = set()

    for ax, artists in list(gate._artists.items()):
        for line in artists.values():
            try:
                line.remove()
            except ValueError:
                pass
        touched_figures.add(ax.figure)

    gate._artists.clear()

    for fig in touched_figures:
        if fig.canvas is not None:
            fig.canvas.draw_idle()


def redraw_gates(ax_table, gates, rainbow=False, dark=False):
    # full refresh fallback only
    for sample in gates:
        for tp in gates[sample]:
            for gate in gates[sample][tp]:
                _remove_gate_artists(gate)

    for sample in gates:
        for tp in gates[sample]:
            for gate in gates[sample][tp]:
                draw_gate(ax_table, sample, tp, gate, rainbow=rainbow, dark=dark)


def adjust_gate_width(ax, ax_table, gates, x, dw):
    axd = ax_table[ax]
    sample = axd.sample
    tp = axd.time_point

    idx = find_nearest_gate(gates, sample, tp, x)
    if idx is None:
        return

    gate0 = gates[sample][tp][idx]
    new_width = gate0.width + dw
    if new_width < 1:
        return

    # width is shared across corresponding gates at the same time point
    for s in gates:
        gate = gates[s][tp][idx]
        gate.width = new_width
        _update_gate_artists(gate)


def move_gate(ax, ax_table, gates, x, dx):
    axd = ax_table[ax]
    sample = axd.sample
    tp = axd.time_point

    idx = find_nearest_gate(gates, sample, tp, x)
    if idx is None:
        return

    # x is independent per sample, so update only this one gate
    gate = gates[sample][tp][idx]
    gate.x += dx
    _update_gate_artists(gate)


def delete_gate(ax, ax_table, gates, x):
    axd = ax_table[ax]
    sample = axd.sample
    tp = axd.time_point

    idx = find_nearest_gate(gates, sample, tp, x)
    if idx is None:
        return

    # corresponding gate index is removed for all samples at this time point
    for s in gates:
        gate = gates[s][tp].pop(idx)
        _remove_gate_artists(gate)


def add_gate(ax, ax_table, gates, x, rainbow=False, dark=False):
    axd = ax_table[ax]
    sample = axd.sample
    tp = axd.time_point

    ref = gates[sample][tp]
    insert_idx = next((i for i, g in enumerate(ref) if g.x > x), len(ref))

    # use width from neighbors if available, otherwise default WIDTH
    if ref:
        if insert_idx > 0:
            width = ref[insert_idx - 1].width
        else:
            width = ref[0].width
    else:
        width = WIDTH

    # add corresponding gate to every sample at this time point
    # same width, same initial x on insertion; x can diverge later
    for s in gates:
        gate = Gate(x, width=width)
        gates[s][tp].insert(insert_idx, gate)
        draw_gate(ax_table, s, tp, gate, rainbow=rainbow, dark=dark)


def draw_gate(ax_table, sample, tp, gate, rainbow=False, dark=False):
    if gate._style["c1"] is None:
        c1 = "black" if dark else "white"
        c2 = next(cycle_colors) if rainbow else "red"
        gate._style["c1"] = c1
        gate._style["c2"] = c2
    else:
        c1 = gate._style["c1"]
        c2 = gate._style["c2"]

    touched_figures = set()

    for ax, axd in ax_table.items():
        if axd.sample != sample or axd.time_point != tp:
            continue

        if ax in gate._artists:
            x, l, r = _gate_positions(gate)
            gate._artists[ax]["center"].set_xdata([x, x])
            gate._artists[ax]["left"].set_xdata([l, l])
            gate._artists[ax]["right"].set_xdata([r, r])
        else:
            _draw_gate_on_axis(ax, gate, c1, c2)

        touched_figures.add(ax.figure)

    for fig in touched_figures:
        if fig.canvas is not None:
            fig.canvas.draw_idle()


def redraw_gates_new_figure(ax_table, gates, rainbow=False, dark=False):
    """
    Recreate all gate artists on a newly created figure.

    Uses only gate geometry (x, width) plus sample/tp matching.
    Any existing gate._artists references are discarded and replaced.
    """
    valid_axes = set(ax_table.keys())

    # Remove any existing gate lines already present on this figure
    for ax in valid_axes:
        for line in list(ax.lines):
            if getattr(line, "_is_gate", False):
                line.remove()

    for sample in gates:
        for tp in gates[sample]:
            for gate in gates[sample][tp]:
                # reset runtime artist refs for this gate
                gate._artists = {}

                # ensure style exists
                if not hasattr(gate, "_style"):
                    gate._style = {"c1": None, "c2": None}

                if gate._style["c1"] is None:
                    c1 = "black" if dark else "white"
                    c2 = next(cycle_colors) if rainbow else "red"
                    gate._style["c1"] = c1
                    gate._style["c2"] = c2
                else:
                    c1 = gate._style["c1"]
                    c2 = gate._style["c2"]

                x = gate.x
                w = gate.width
                l = x - w / 2
                r = x + w / 2

                for ax, axd in ax_table.items():
                    if axd.sample != sample or axd.time_point != tp:
                        continue

                    center = ax.axvline(x, color=c1, lw=1)
                    center._is_gate = True

                    left = ax.axvline(l, color=c2, lw=1, ls="--")
                    left._is_gate = True

                    right = ax.axvline(r, color=c2, lw=1, ls="--")
                    right._is_gate = True

                    gate._artists[ax] = {
                        "center": center,
                        "left": left,
                        "right": right,
                    }

    figs = {ax.figure for ax in valid_axes}
    for fig in figs:
        if fig.canvas is not None:
            fig.canvas.draw_idle()


def add_keybinding_panel(fig):
    help_text = (
        "Controls   |   "
        "Left click: add   |   "
        "Right click: delete   |   "
        "← / →: move   |   "
        "+ / -: width   |   "
        "Shift + scroll: width"
    )

    # create space above subplots
    fig.subplots_adjust(top=0.88)

    fig.text(
        0.5,
        0.95,
        help_text,
        ha="center",
        va="top",
        fontsize=10,
        family="monospace",
        bbox=dict(boxstyle="round", facecolor="white", alpha=0.9, edgecolor="gray"),
    )


def redraw_gates_new_figure(ax_table, gates, rainbow=False, dark=False):
    """
    Recreate all gate artists on a newly created figure.

    Uses only gate geometry (x, width) plus sample/tp matching.
    Any existing gate._artists references are discarded and replaced.
    """
    valid_axes = set(ax_table.keys())

    # Remove any existing gate lines already present on this figure
    for ax in valid_axes:
        for line in list(ax.lines):
            if getattr(line, "_is_gate", False):
                line.remove()

    for sample in gates:
        for tp in gates[sample]:
            for gate in gates[sample][tp]:
                # reset runtime artist refs for this gate
                gate._artists = {}

                # ensure style exists
                if not hasattr(gate, "_style"):
                    gate._style = {"c1": None, "c2": None}

                if gate._style["c1"] is None:
                    c1 = "black" if dark else "white"
                    c2 = next(cycle_colors) if rainbow else "red"
                    gate._style["c1"] = c1
                    gate._style["c2"] = c2
                else:
                    c1 = gate._style["c1"]
                    c2 = gate._style["c2"]

                x = gate.x
                w = gate.width
                l = x - w / 2
                r = x + w / 2

                for ax, axd in ax_table.items():
                    if axd.sample != sample or axd.time_point != tp:
                        continue

                    center = ax.axvline(x, color=c1, lw=1)
                    center._is_gate = True

                    left = ax.axvline(l, color=c2, lw=1, ls="--")
                    left._is_gate = True

                    right = ax.axvline(r, color=c2, lw=1, ls="--")
                    right._is_gate = True

                    gate._artists[ax] = {
                        "center": center,
                        "left": left,
                        "right": right,
                    }

    figs = {ax.figure for ax in valid_axes}
    for fig in figs:
        if fig.canvas is not None:
            fig.canvas.draw_idle()

# %% [code]
#Interactive selection of integration gates.
#Note that a checkpoint is provided with the gates selected for figure 3D
#change reprocess to True to allow gate selection


reprocess=False
plt.close("all")

from functools import partial
if reprocess:
    %matplotlib ipympl
    try:
        fig.canvas.mpl_disconnect(cid)
    except NameError:
        pass
    # Protecting gates from overwrite and ensuring they are initialised
    long_samples = samples
    try:
        need_init = not long_gates
    except NameError:
        need_init = True
    if need_init:
        long_gates = {}
        for sample in long_samples:
            long_gates[sample] = {}
            for tp in extracted_lanes[sample].keys():
                long_gates[sample][tp] = []

    fig, axs, axtable = plot_sample_set(long_samples, narrow=True)
    if not need_init:
        redraw_gates_new_figure(axtable, long_gates, )
    cid = fig.canvas.mpl_connect(
        "button_press_event",
        partial(on_click, ax_table=axtable, gates=long_gates),
    )

    cid = fig.canvas.mpl_connect(
        "key_press_event",
        partial(on_key, ax_table=axtable, gates=long_gates),
    )
    fig.canvas.mpl_connect(
        "scroll_event",
        partial(on_scroll, ax_table=axtable, gates=long_gates),
    )
    plt.show()

# %% [code]
plt.close("all")
dest = folder / "checkpoints/gates.pkl"

if reprocess:
    with open(dest, "wb") as f:
        pickle.dump(long_gates, f)
else:
    gates = {}
    with open(dest, "rb") as f:
        long_gates = pickle.load(f)

# %% [code]
#double check that everything is A-OK with non interactive figure
tweak = False
savefig =True
to_check = samples
gates_check = long_gates
if len(to_check)<=2:
    narrow = False
    supernarrow = False
elif len(to_check) <=4:
    narrow=True
    supernarrow = False
else:
    narrow=False
    supernarrow=True
if not tweak:
    %matplotlib inline
    plt.close("all")
elif tweak:
    %matplotlib ipympl
fig, axs, axtable = plot_sample_set(to_check, narrow = narrow, supernarrow=supernarrow)
redraw_gates_new_figure(axtable, gates_check, )
if tweak:
    cid = fig.canvas.mpl_connect(
        "button_press_event",
        partial(on_click, ax_table=axtable, gates=new_long_gates),
    )

    cid = fig.canvas.mpl_connect(
        "key_press_event",
        partial(on_key, ax_table=axtable, gates=new_long_gates),
    )
    fig.canvas.mpl_connect(
        "scroll_event",
        partial(on_scroll, ax_table=axtable, gates=new_long_gates),
    )
fig.savefig(analysis_folder/f"peak_vs_lanes_{'_'.join(to_check)}.png", format="PNG")
plt.show()

# %% [code]
#define functions for integration

def plot_peaks_with_integration(
    samples,
    curves,
    peak_results,
    peak_integrals,
    time_points_labels,
    palettes,
    show_text=True,
    show_boundaries=True,
    show_area=True,
    layout="grid",  # "grid" or "rows"
    sharey=False,
):
    n_tp = len(time_points_labels)
    n_s = len(samples)

    if layout == "grid":
        fig, axes = plt.subplots(
            n_tp,
            n_s,
            figsize=(20, 4 * n_tp),
            sharex=True,
        )
        if n_tp == 1:
            axes = np.array([axes])

    elif layout == "rows":
        fig, axes = plt.subplots(
            n_tp * n_s, 1, figsize=(14, 3 * n_tp * n_s), sharex=True, sharey=sharey
        )
        axes = np.atleast_1d(axes)

    idx = 0

    for i, tp in enumerate(time_points_labels):
        for j, sample in enumerate(samples):

            if layout == "grid":
                ax = axes[i, j]
            else:
                ax = axes[idx]

            y = curves[sample][tp]
            cmap = palettes[sample]
            color = cmap(0.6)

            ax.plot(y, color=color)

            peaks = peak_results[sample][tp]["peaks"]
            props = peak_results[sample][tp]["properties"]
            integrals = peak_integrals[sample][tp]

            for k, p in enumerate(peaks):
                left = int(props["left_bases"][k])
                right = int(props["right_bases"][k])
                x = np.arange(left, right + 1)

                gate_color = plt.cm.tab20(k % 20)

                ax.scatter(p, y[p], color=gate_color, s=25)

                if show_text:
                    ax.text(p, y[p], str(k), fontsize=9, ha="center", va="bottom")

                if show_area:
                    ax.fill_between(
                        x,
                        0,
                        y[left : right + 1],
                        color=gate_color,
                        alpha=0.30,
                    )

                if show_boundaries:
                    ax.axvline(left, linestyle="--", linewidth=1.5, color=gate_color)
                    ax.axvline(right, linestyle="--", linewidth=1.5, color=gate_color)

                area = integrals[k]

                if show_text:
                    ax.text(
                        p,
                        y[p] * 1.05,
                        f"{area:.2f}",
                        fontsize=8,
                        ha="center",
                        va="bottom",
                    )

            if layout == "grid":
                if i == 0:
                    ax.set_title(sample)
                if j == 0:
                    ax.set_ylabel(tp)
            else:
                ax.set_title(f"{sample} | {tp}")

            idx += 1

    fig.supxlabel("index")
    fig.supylabel("signal")
    fig.tight_layout()
    return fig


def reconstruct_peaks_and_integrals(gates, moved):
    peak_results = {}
    peak_integrals = {}

    for sample, times in gates.items():
        peak_results[sample] = {}
        peak_integrals[sample] = {}

        for tp, gate_list in times.items():
            curve = moved[sample][tp]
            n = len(curve)

            peaks = []
            left_bases = []
            right_bases = []
            integrals = []

            for g in gate_list:
                left = int(round(g.x - g.width / 2))
                right = int(round(g.x + g.width / 2))

                left = max(0, left)
                right = min(n - 1, right)

                seg = curve[left : right + 1]
                p = left + int(np.argmax(seg))

                area = np.trapezoid(seg)

                peaks.append(p)
                left_bases.append(left)
                right_bases.append(right)
                integrals.append(area)

            peak_results[sample][tp] = {
                "peaks": np.asarray(peaks),
                "properties": {
                    "left_bases": np.asarray(left_bases),
                    "right_bases": np.asarray(right_bases),
                },
            }

            peak_integrals[sample][tp] = integrals

    return peak_results, peak_integrals

# %% [code]
# calculate integrals and find peaks within gates
peak_results, peak_integrals = reconstruct_peaks_and_integrals(
    long_gates, baselined_curves
)
# make figures
for i in [samples]:
    samples_to_plot = i
    fig = plot_peaks_with_integration(
        samples=samples_to_plot,
        curves=baselined_curves,
        peak_results=peak_results,
        peak_integrals=peak_integrals,
        time_points_labels=time_points_labels,
        palettes=palettes,
        sharey=True,
    )
    fig.savefig(
        analysis_folder / f"integrated_peaks_{'_'.join(samples_to_plot)}.png", format="PNG"
    )

# %% [code]
# plot the integration series.
column_names = ["Ub10+", "Ub6", "Ub5", "Ub4", "Ub3", "Ub2", "Ub"]

dataframes = {}

for sample in samples:
    rows = peak_integrals[sample]
    columns = column_names
    df = pd.DataFrame.from_dict(rows, orient="index", columns=columns)
    df.index.name = "time_point"
    df = df.sort_index()
    if not natural_order:
        df = df.iloc[::-1]
    dataframes[sample] = df

# %% [code]
overwrite=False
%matplotlib inline
def plot_peak_timeseries(samples, dataframes, log_x=True, last_point=True, sharey=False):

    ref_df = dataframes[samples[0]]
    peaks = ref_df.columns
    n_peaks = len(peaks)

    fig, axes = plt.subplots(1, n_peaks, figsize=(4 * n_peaks, 4), sharey=sharey)

    if n_peaks == 1:
        axes = [axes]

    colors = dict(zip(samples, ["black","#e52521"]))
    markers = dict(zip(samples, ["o", "^"]))

    for sample in samples:
        df = dataframes[sample]
        for ax, peak in zip(axes, peaks):
            x = df.index + 0.1 #log axis and time 0
            y = df[peak]
            ax.plot(x, y, color=colors[sample], linewidth=1)
            ax.scatter(x, y, color=colors[sample], label=sample, marker=markers[sample], facecolor='none')

            if log_x:
                ax.set_xscale("log", base=2)

            ax.set_title(peak)
            ax.set_xlabel("time_point (s)")
            if not last_point:
                if not natural_order:
                    tps = x[::-1]
                else:
                    tps = x
                ax.set_xlim(0, tps[-2] + 10)
    axes[0].set_ylabel("intensity")
    axes[-1].legend()
    fig.tight_layout()
    return fig, axes


fig, axes = plot_peak_timeseries(samples, dataframes, log_x=True, sharey=0)#, last_point=False)
fig.savefig(analysis_folder / f"curves{"_".join(samples_to_plot)}.png", format="PNG")
fig.savefig(analysis_folder / f"curves{"_".join(samples_to_plot)}.pdf", format="PDF")
plt.show()
plt.close("all")

# %% [code]
````
