## Supplementary code for "A ubiquitin chain-feeding mechanism for BRCA1-A": Licence.txt

The MIT License (MIT)

Copyright (c) 2026 Andrea G Murachelli
ORCID: 0000-0002-7048-5758

Permission is hereby granted, free of charge, to any person obtaining a
copy of this software and associated documentation files (the
"Software"), to deal in the Software without restriction, including
without limitation the rights to use, copy, modify, merge, publish,
distribute, sublicense, and/or sell copies of the Software, and to
permit persons to whom the Software is furnished to do so, subject to
the following conditions:

The above copyright notice and this permission notice shall be included
in all copies or substantial portions of the Software.

THE SOFTWARE IS PROVIDED "AS IS", WITHOUT WARRANTY OF ANY KIND, EXPRESS
OR IMPLIED, INCLUDING BUT NOT LIMITED TO THE WARRANTIES OF
MERCHANTABILITY, FITNESS FOR A PARTICULAR PURPOSE AND NONINFRINGEMENT.
IN NO EVENT SHALL THE AUTHORS OR COPYRIGHT HOLDERS BE LIABLE FOR ANY
CLAIM, DAMAGES OR OTHER LIABILITY, WHETHER IN AN ACTION OF CONTRACT,
TORT OR OTHERWISE, ARISING FROM, OUT OF OR IN CONNECTION WITH THE
SOFTWARE OR THE USE OR OTHER DEALINGS IN THE SOFTWARE.
