## Supplementary code for "A ubiquitin chain-feeding mechanism for BRCA1-A": README.txt

Zip File Contents
=================

Figure_3D.ipynb
----------------
This Jupyter notebook contains the analysis pipeline used for processing
the gels shown in Figure 3D of the manuscript and for generating the
densitometry curves presented in Figure 3E.

Environment setup instructions are provided in the first cell of the
notebook. Once the environment is configured, running the notebook will
reproduce the analysis used to generate the figures.

The notebook includes several interactive processing steps, including:
- image background subtraction
- image cropping
- lane selection
- integration gate selection

These interactive sections are disabled by default using the flag:

    reprocess = False

Results from the interactive steps used to generate the manuscript
figures were saved as checkpoint files using pickle and are loaded
automatically when the notebook is executed.

The code for the interactive sections is included for reference and can
be enabled by setting:

    reprocess = True

Note that enabling reprocessing requires the interactive steps to be
performed manually for the notebook to run successfully. Doing so will
also overwrite the precalculated checkpoint results.


Folders
=======

background_subtraction
----------------------
Contains the processed analysis results used for Figures 3D and 3E.

figures
-------
Contains gel images with lane selections and the corresponding generated
curves.

raw_tif
-------
Contains the original TIFF images and background images used for
background subtraction.


Additional Notes
================

Analysis for the other figures in the manuscript was performed using the
same processing pipeline.
