## Supplementary code for "A ubiquitin chain-feeding mechanism for BRCA1-A": README.TXT

Files
=====

deltains_long_chains.tif
------------------------
Raw scan of the gel corresponding to the long-chain digestion experiment
performed with BRCA1-A (Iso2).

wt_long_chains.tif
------------------
Raw scan of the gel corresponding to the long-chain digestion experiment
performed with wild-type BRCA1-A.

blank.tif
---------
Scan of the empty gel tray used for background subtraction.


License and Usage
=================

Files in this directory are provided for reproducibility purposes only.

These materials are not distributed under the MIT License. All rights
remain with the respective copyright holders.
