## Supplementary figures and images for "A ubiquitin chain-feeding mechanism for BRCA1-A"

### background_correction.png

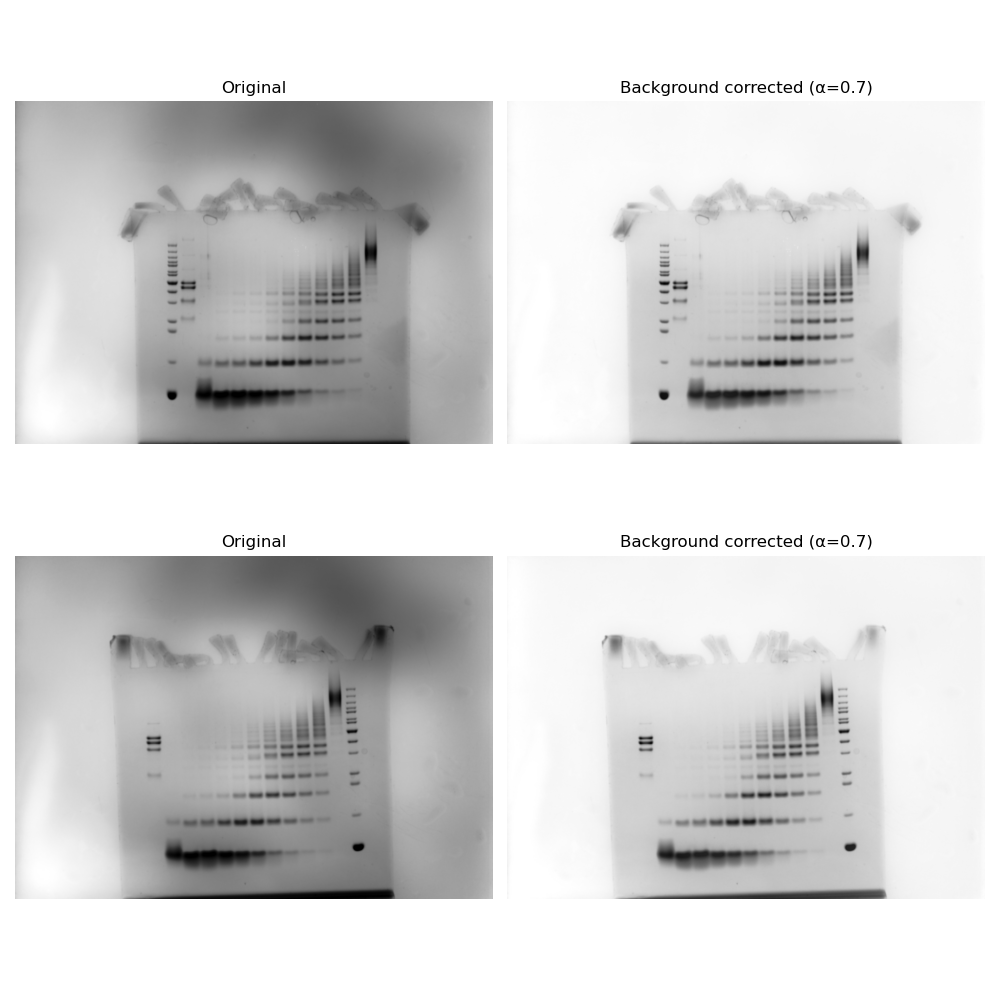

### background_subtraction_coefficients.png

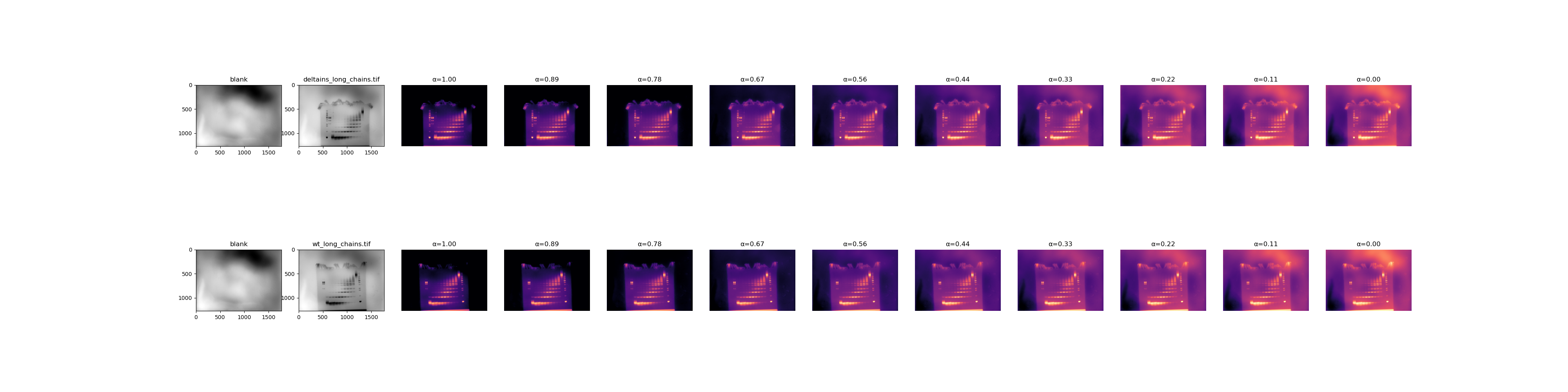

### blank.tif

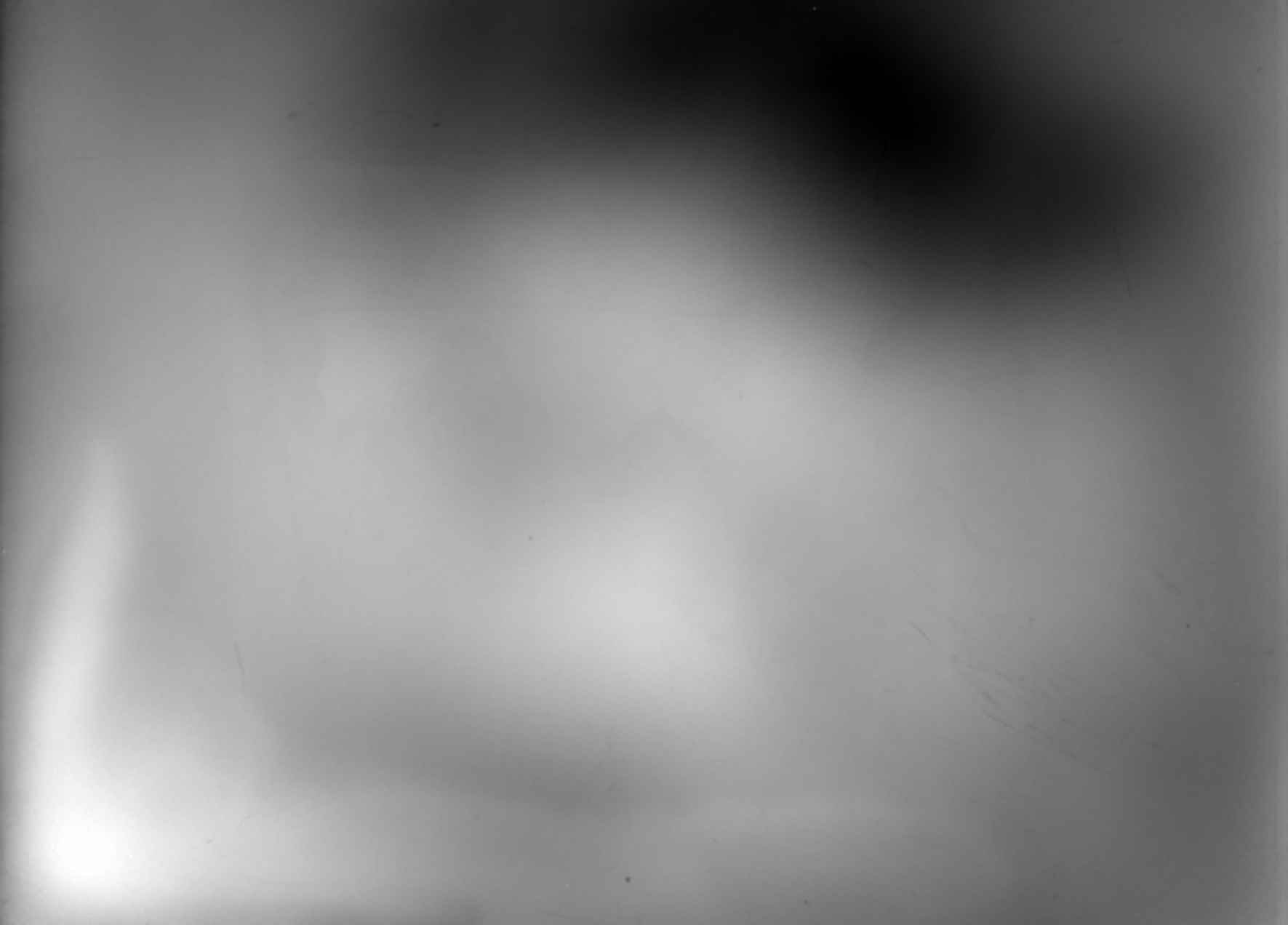

### curvesdeltains_wt.pdf

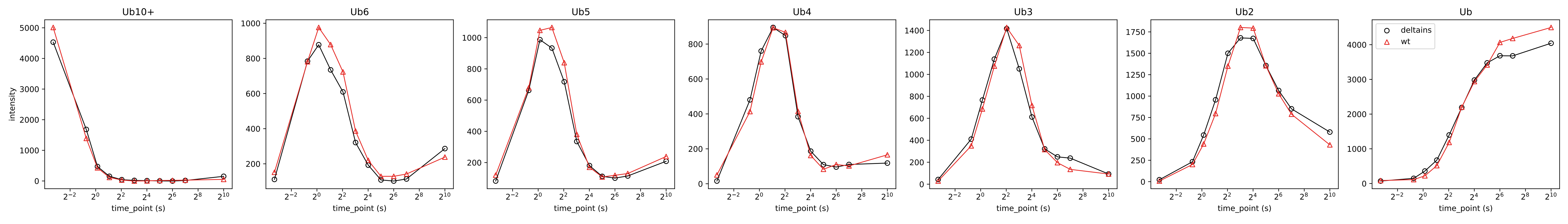

### curvesdeltains_wt.pdf

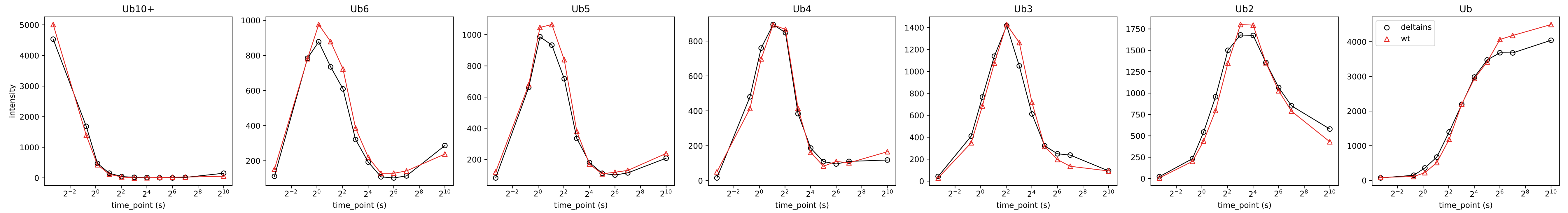

### curvesdeltains_wt.png

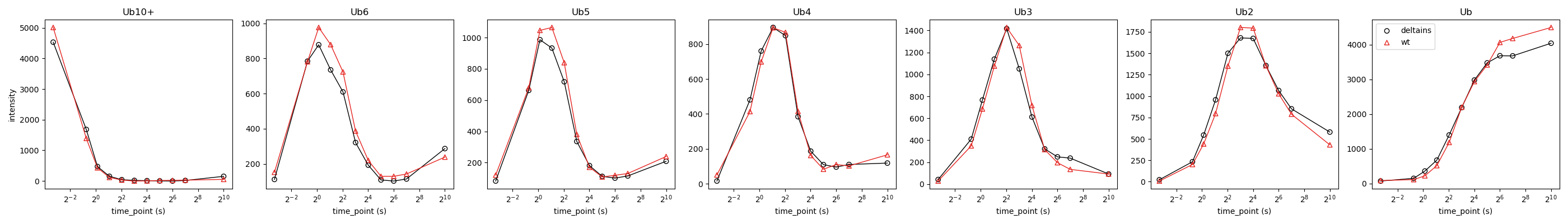

### deltains_baseilne_correction.tif

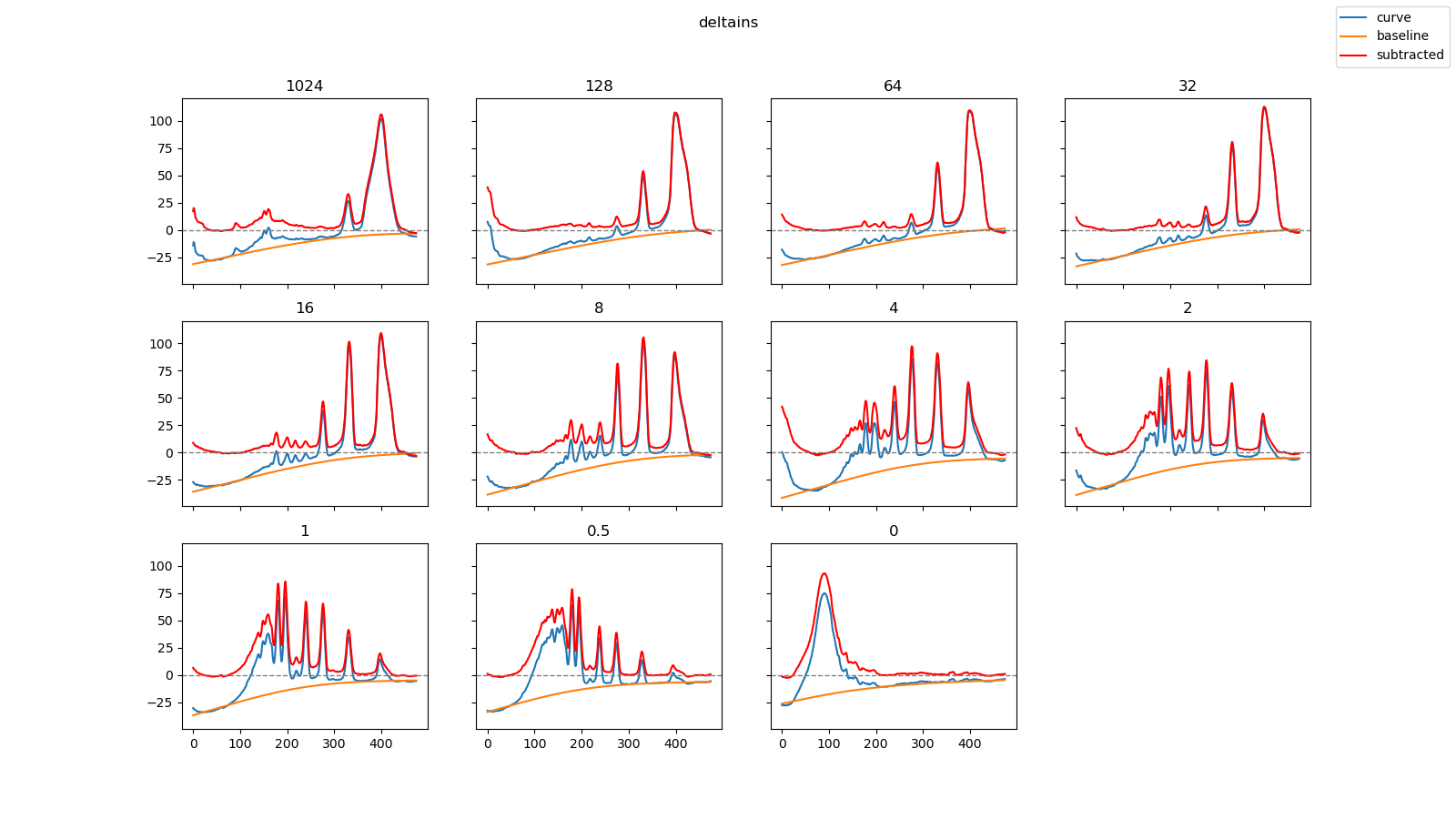

### deltains_long_chains.tif

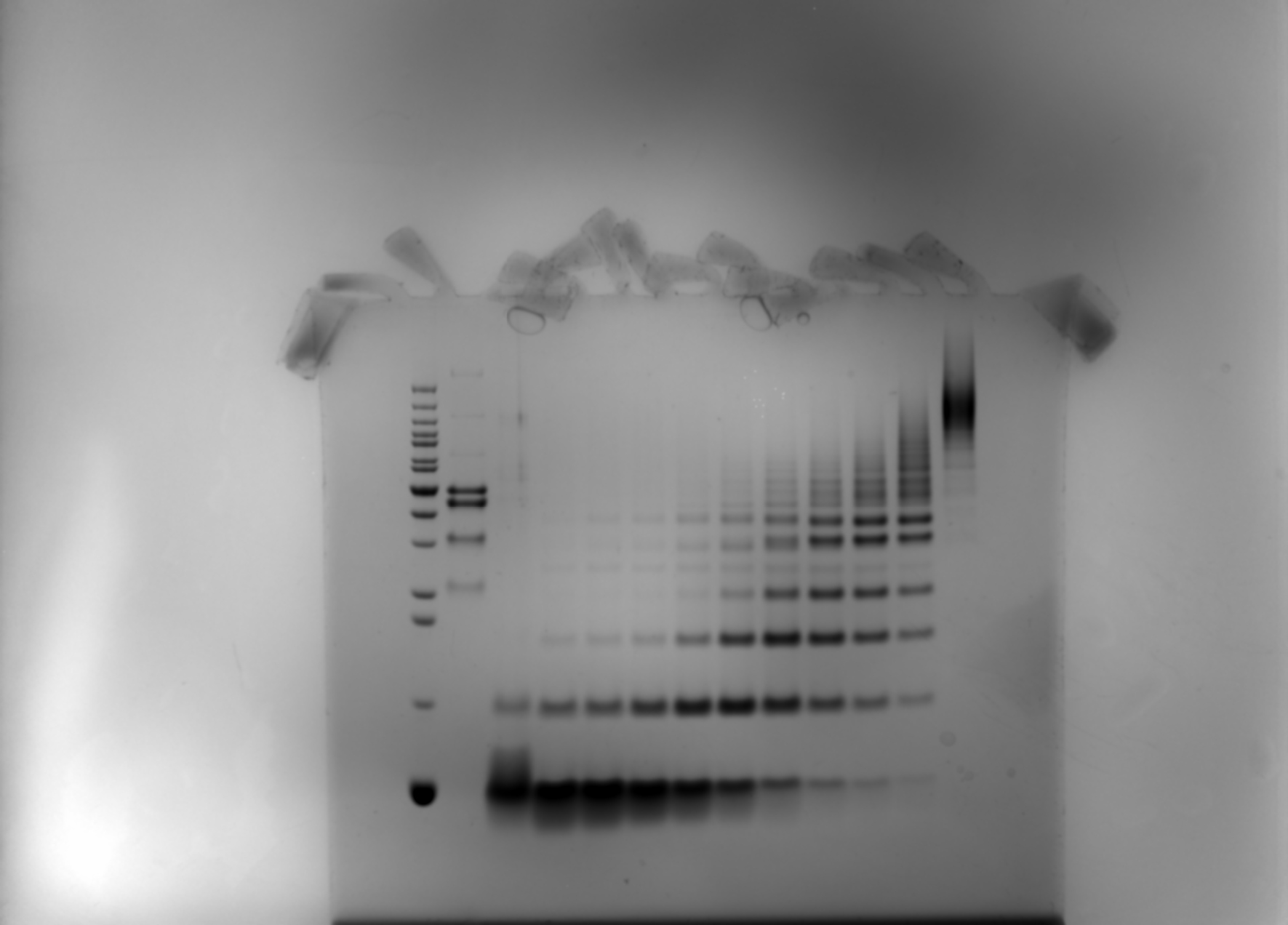

### deltains_long_chains.tif_subtracted.tif

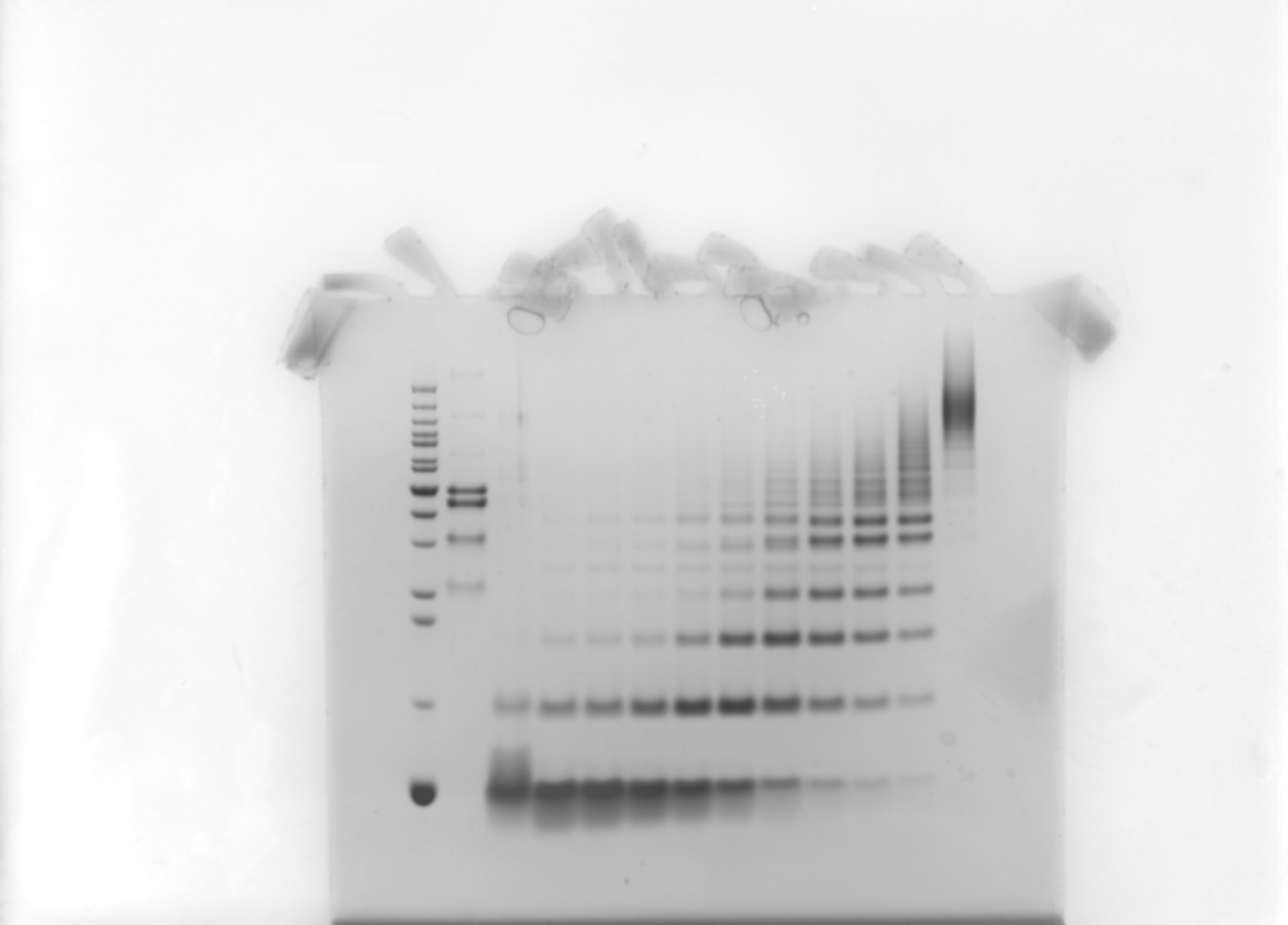

### deltains_long_chains.tif_subtracted_cropped.tif

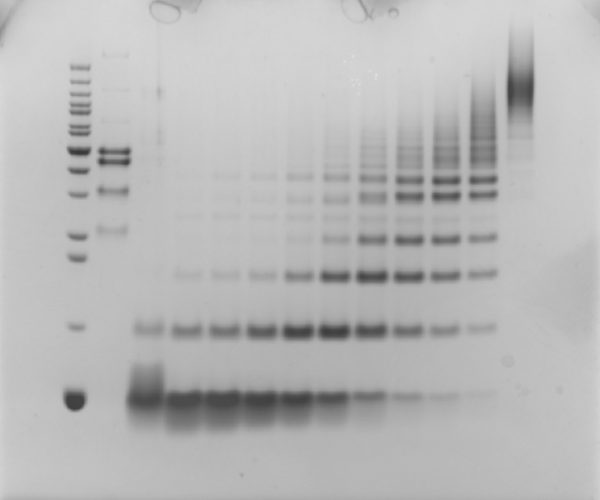

### integrated_peaks_deltains_wt.png

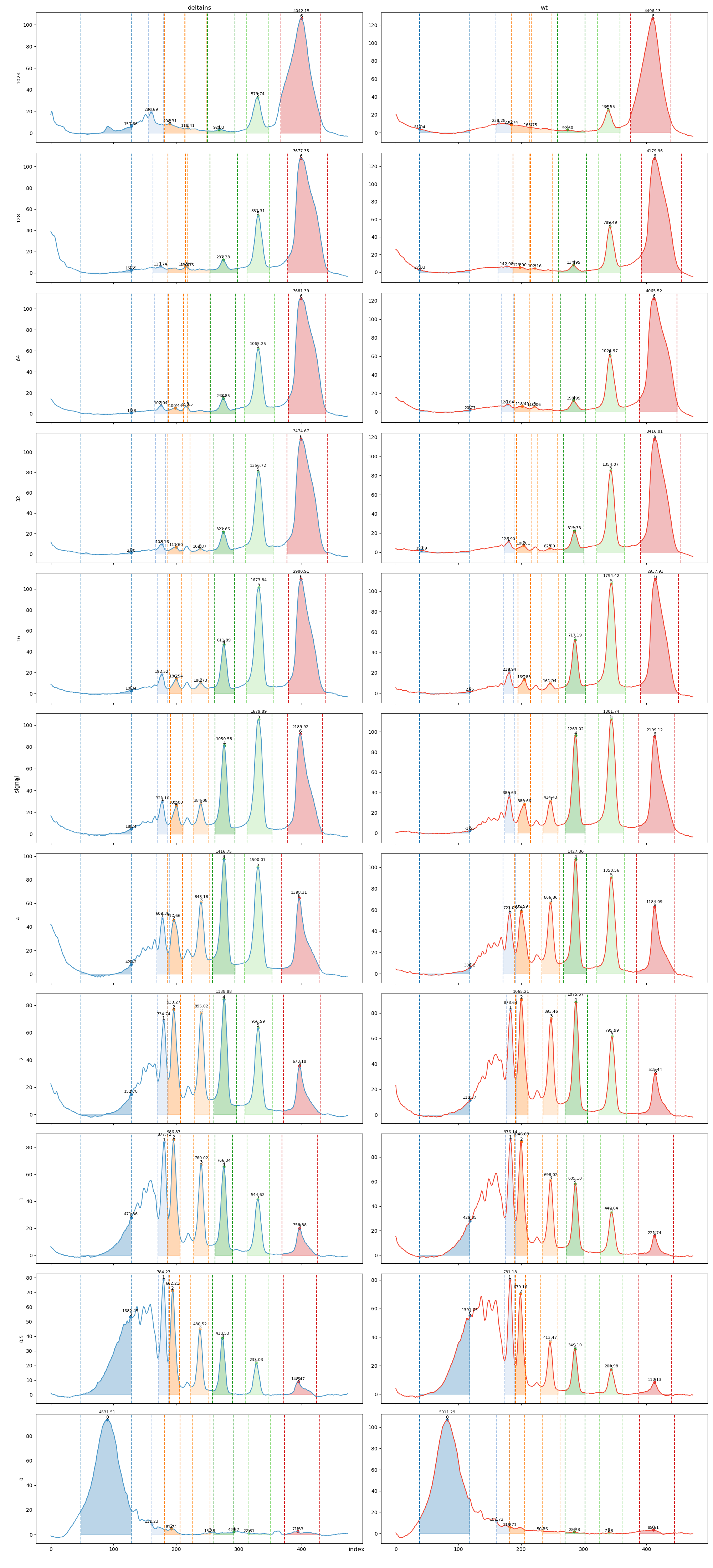

### lanes_vs_curves_comparison_deltains.png

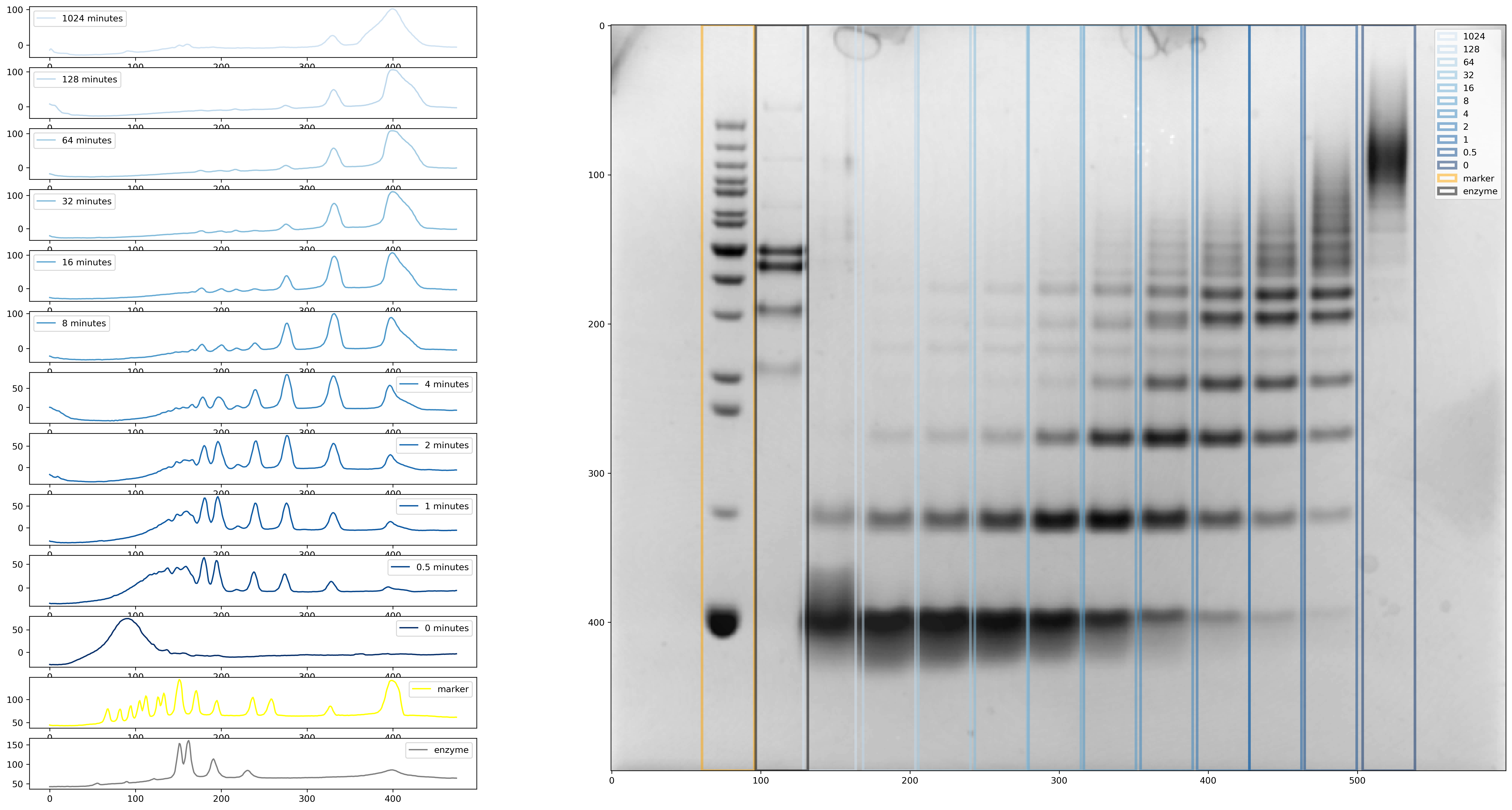

### lanes_vs_curves_comparison_wt.png

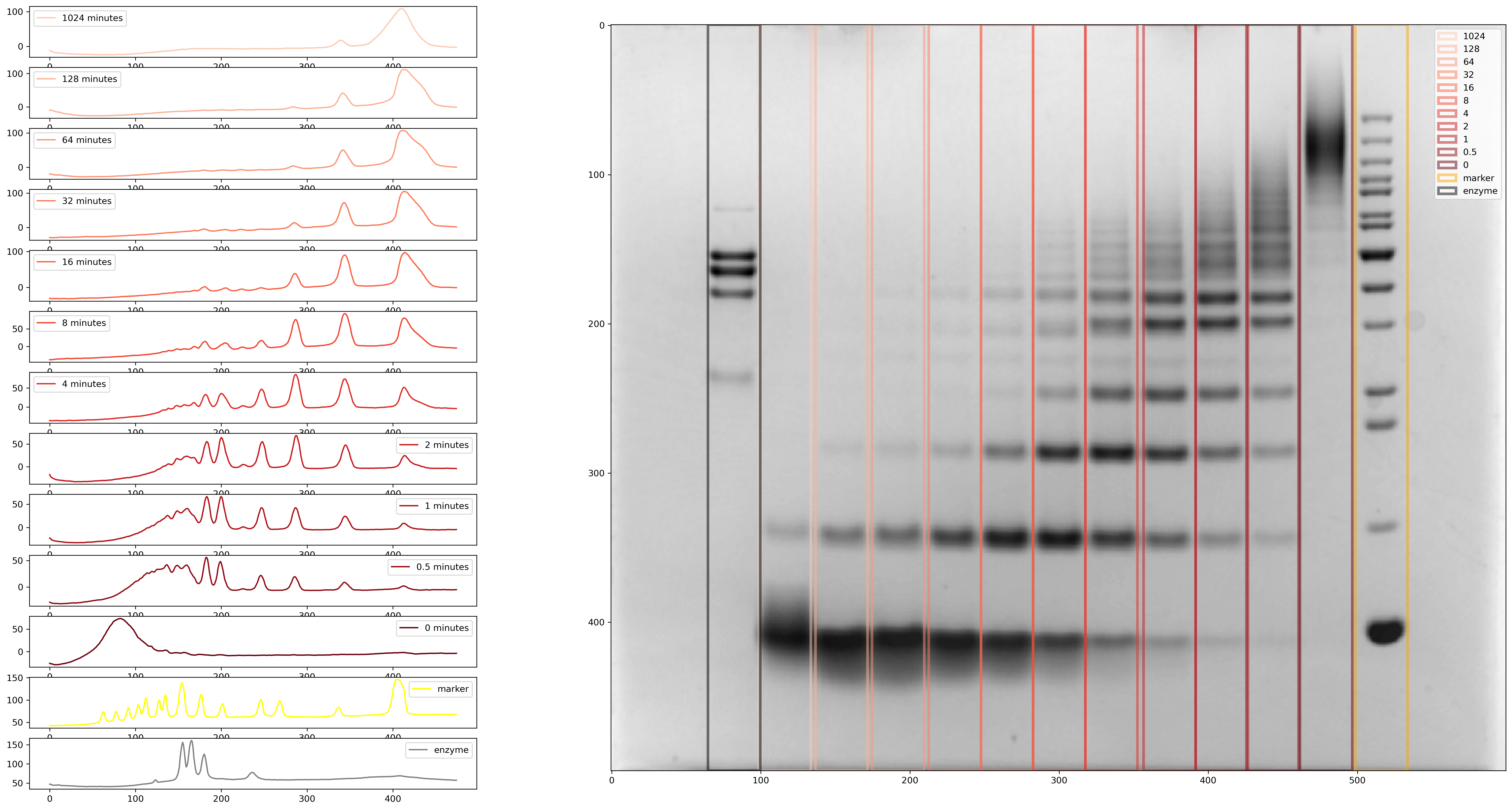

### marker_intensity_comparison.tif

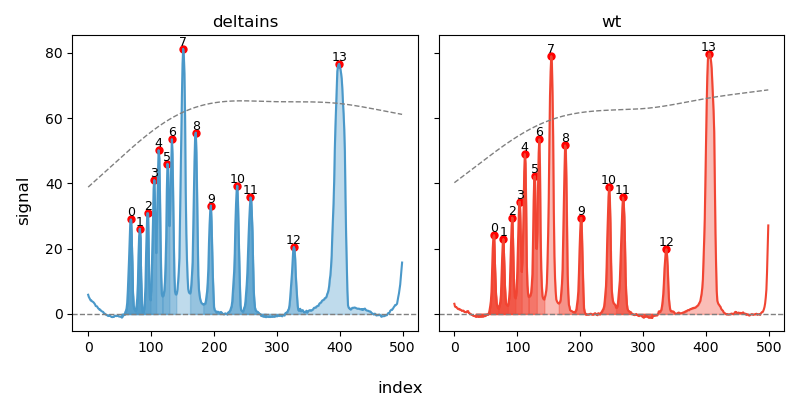

### peak_vs_lanes_deltains_wt.png

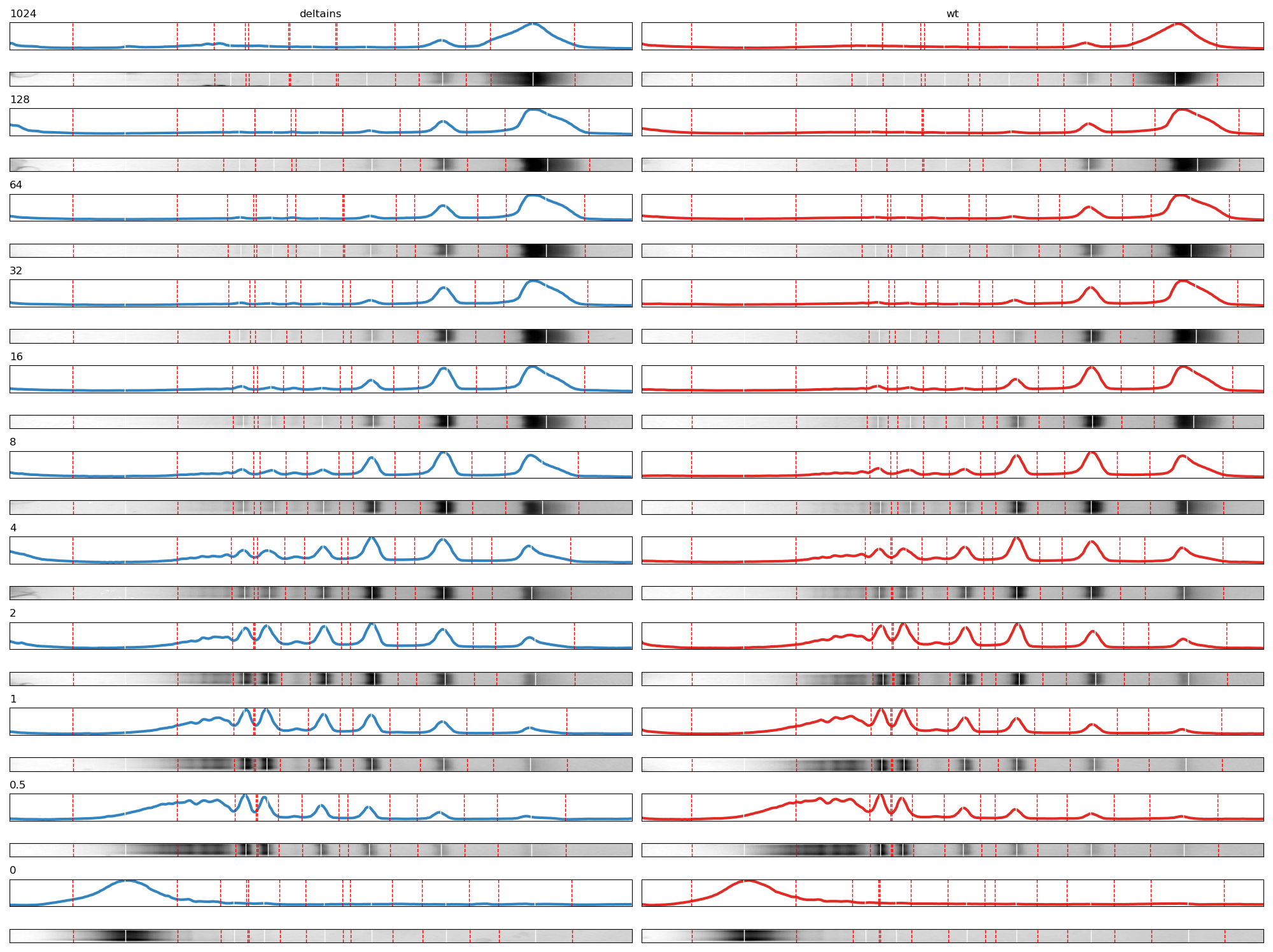

### selected_lanes.png

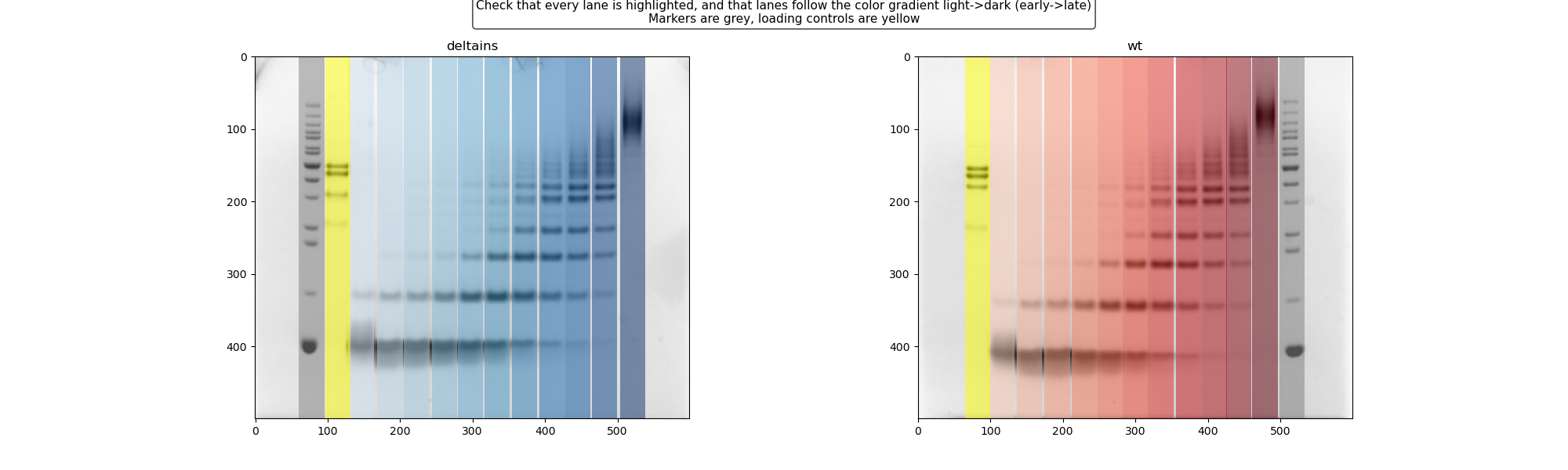

### wt_baseilne_correction.tif

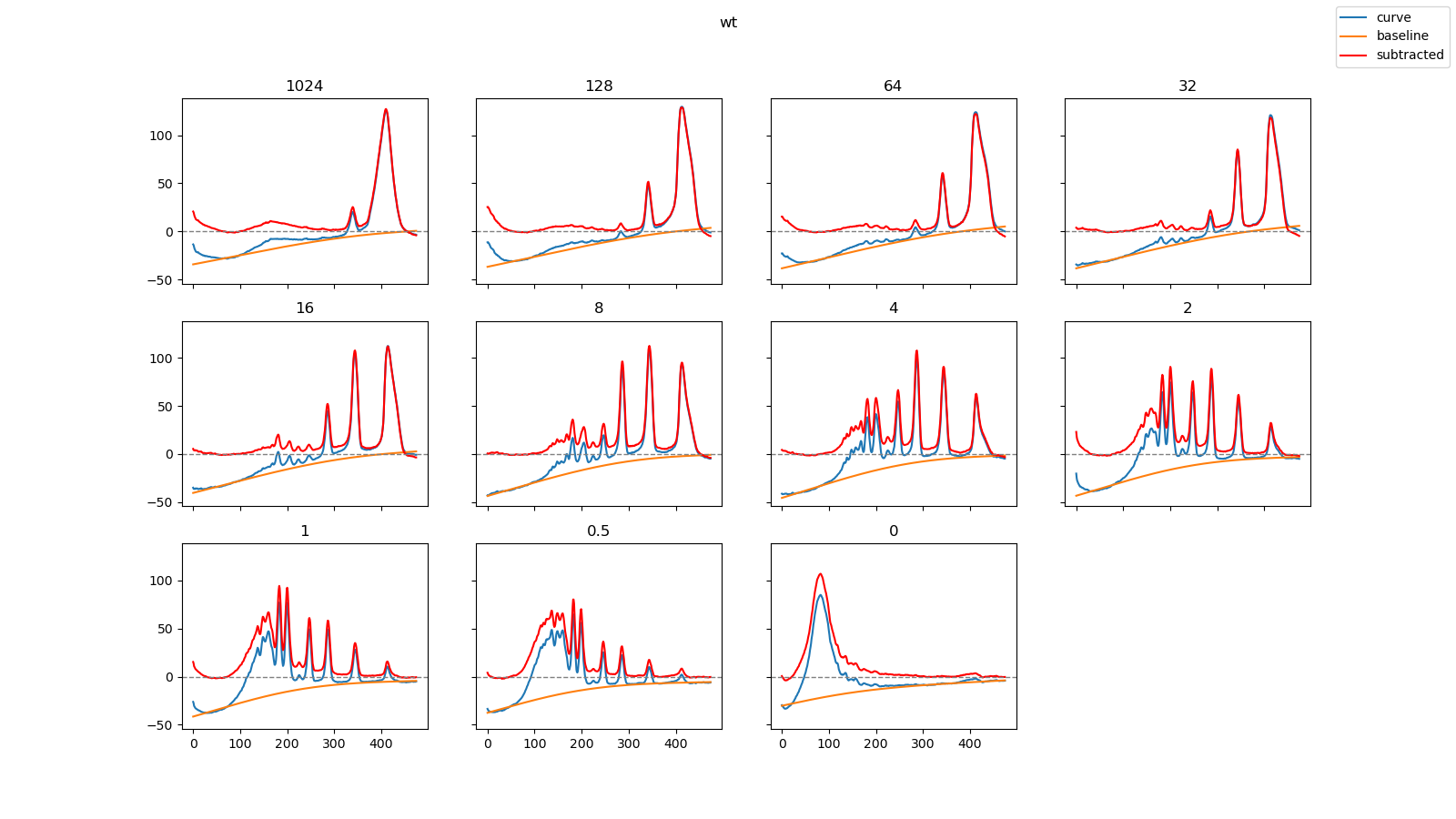

### wt_baseline_parameter_estimation.tif

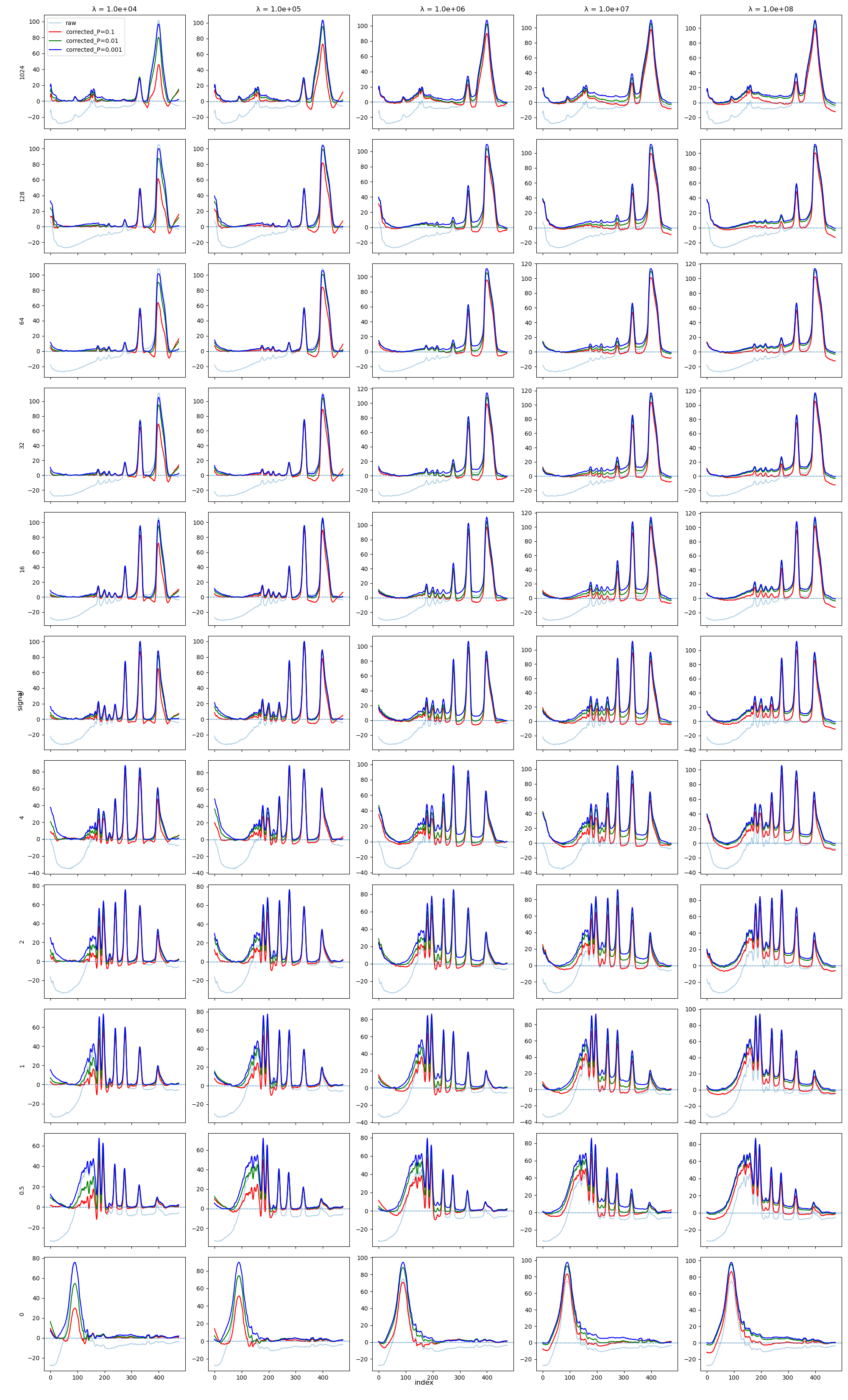

### wt_long_chains.tif

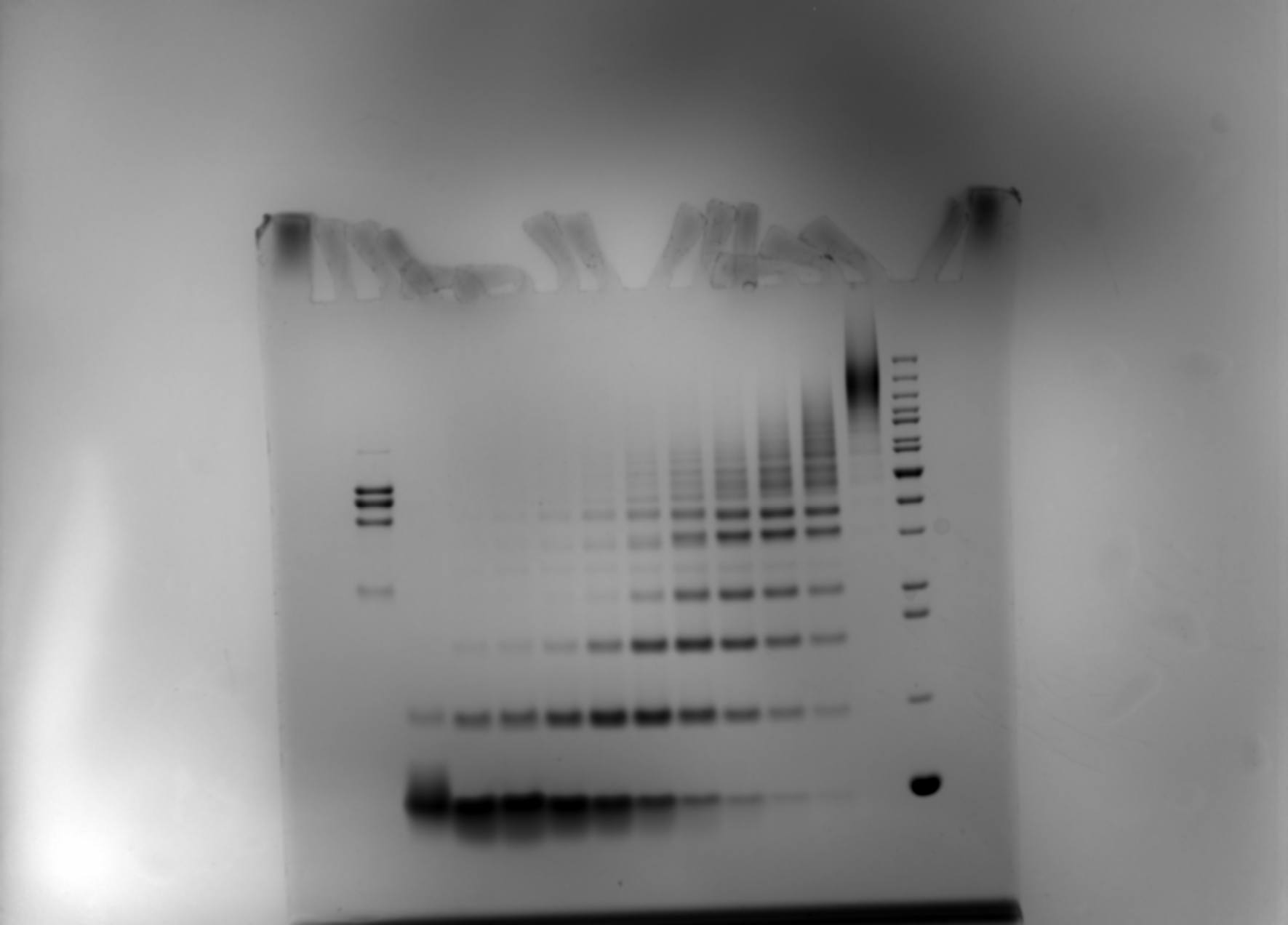

### wt_long_chains.tif_subtracted.tif

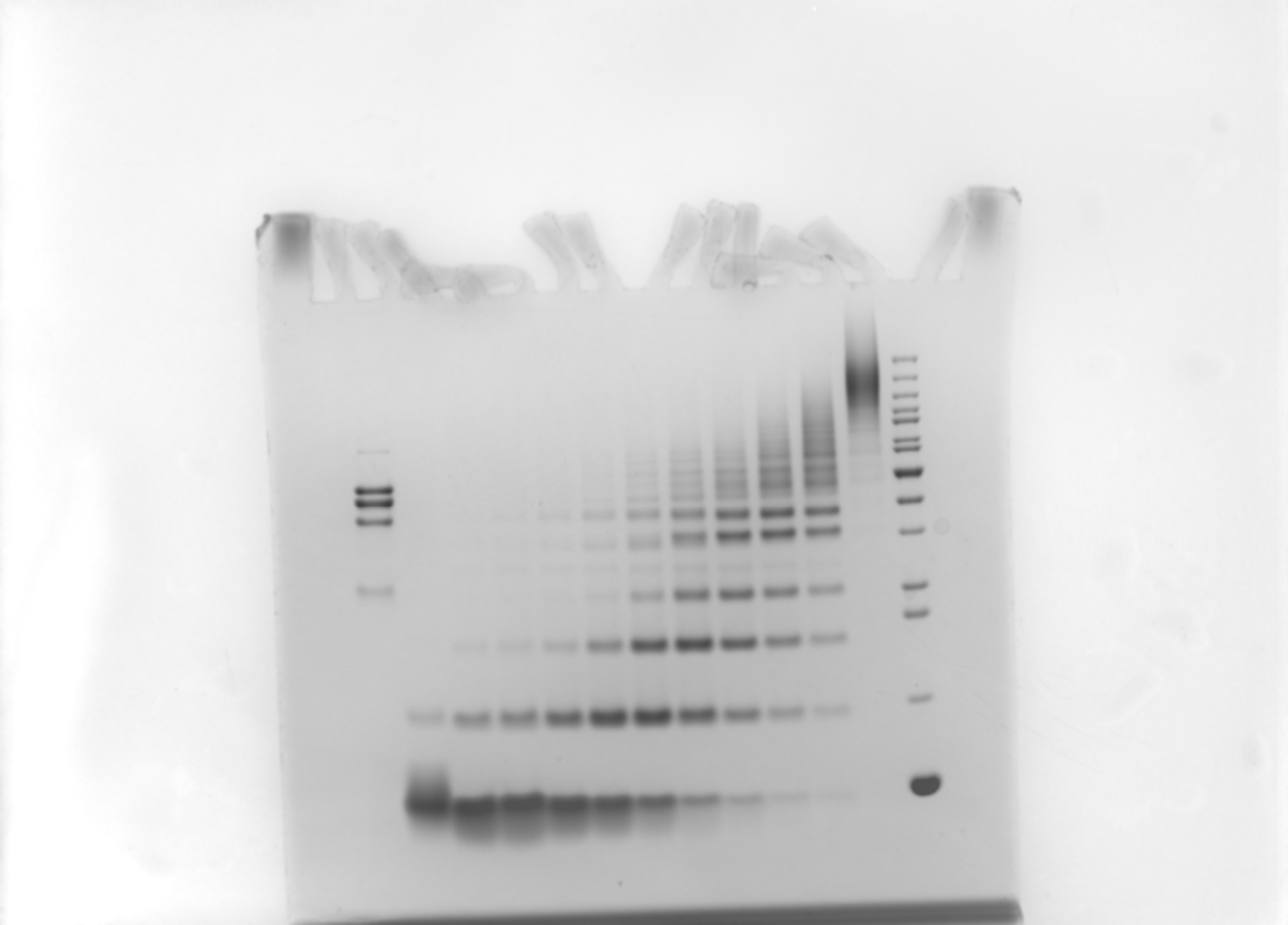

### wt_long_chains.tif_subtracted_cropped.tif

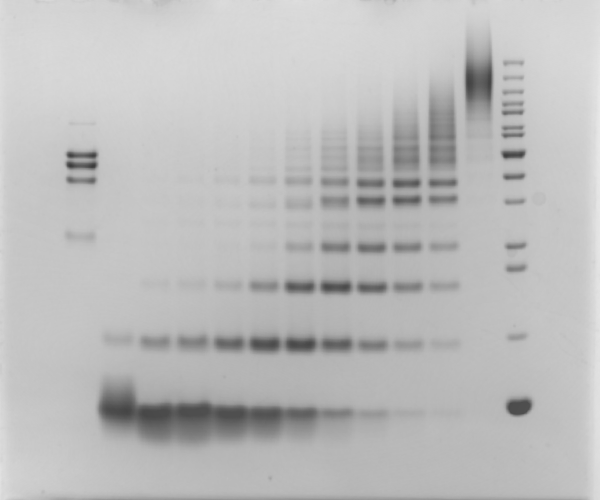
